# Computational Discovery of an Allosteric Pocket on APOBEC3B and the Impact of Metal Contamination on Inhibitor Validation

**DOI:** 10.1101/2024.04.25.591187

**Authors:** Mackenzie K. Wyllie, Katherine F. M. Jones, Özlem Demir, Rahul Dadabhau Kardile, Michael J. Grillo, Clare K. Morris, Brandon B. Schuldt, Farzana Kabir, Sophia P. Hirakis, Michael A. Walters, Reuben S. Harris, Rommie E. Amaro, Daniel A. Harki

## Abstract

The APOBEC3 (A3) family of enzymes are zinc metalloenzymes that catalyze the conversion of 2′- deoxycytidine to 2′-deoxyuridine in single stranded DNA. APOBEC3B (A3B), a member of the A3 family, has emerged as a key driver of genomic instability in many cancer types by mutating host DNA to drive tumorigenesis and therapy resistance. Small molecule inhibitors of A3B would extend the durability of current therapies by limiting mutations that promote tumor escape and therapy resistance. Consequently, a computer aided drug discovery (CADD) campaign was employed to identify inhibitors of A3B. Through molecular dynamics (MD) simulations and computational solvent mapping analysis, we identified a novel putative allosteric pocket on the c-terminal domain of A3B and virtually screened the ChemBridge Diversity Set (∼110,000 small molecules) against both the active and predicted allosteric sites. High scoring compounds were selected for *in vitro* testing and triage, resulting in 13 candidate inhibitors. Using cysteine reactive probes, one of the original hit compounds was mapped to the computationally identified allosteric pocket. However, after resynthesis of representative chemotypes and further investigation using *in vitro* assays, none of the chemotypes retained inhibitory activity. Further analysis revealed that most of the inhibition observed in the primary assay was due to metal contamination in the screening sample, which resulted in the removal of the catalytic zinc from the enzyme active site. Although validated A3B inhibitors were not discovered in this study, we report a ligandable allosteric site on A3B and several cautionary insights for researchers developing small molecule inhibitors of zinc metalloenzymes.

**TOC graphic:** 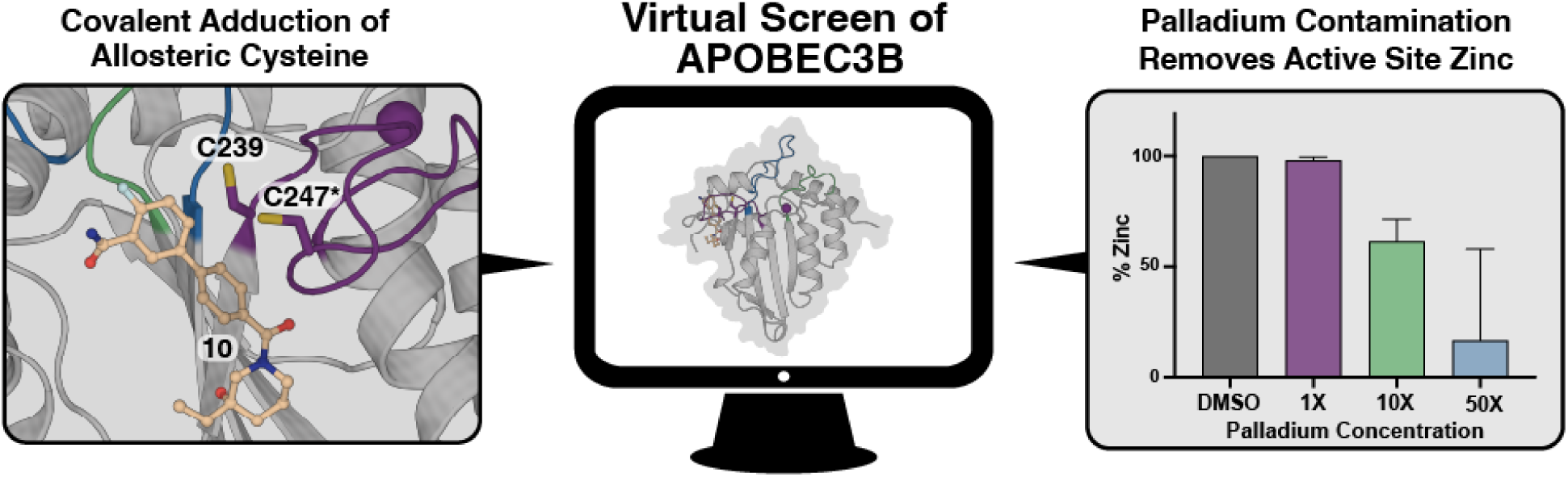

## Introduction

The apolipoprotein B mRNA editing catalytic polypeptide-like 3B (APOBEC3B, A3B) enzyme has emerged as a key endogenous driver of genomic instability in many cancers.^1–3^ A3B is one of seven mammalian A3 cytidine deaminases (A3A-H, excluding E) responsible for converting cytosine to uracil in single stranded (ss)DNA (**Figure 1A**). The enzymatic conversion of C-to-U by APOBEC3 (A3) enzymes occurs through a zinc-mediated hydrolysis mechanism where a carbonyl is installed at the 4- position of cytosine, resulting in the loss of ammonia (**Figure 1B**). Through various base repair pathways, these C-to-U lesions can then be immortalized as genomic C-to-T and C-to-G mutations.^4, 5^ The A3 enzymes were initially discovered as viral restriction factors, however, the use of next generation sequencing has identified APOBEC3 mutational signatures in over 70% of all cancer genomes.^4, 6^ Although all A3 enzymes are capable of deaminating cytidine to various degrees, A3B stands out as the most highly expressed family member in bladder, breast, cervical, head/neck, and lung cancer, where it correlates with poor prognosis and increased mutational burden.^1,7^ A3B-induced genomic instability has been shown to promote tumor evolution, therapeutic resistance, and immune evasion.^3,8–10^ Consequently, A3B inhibition has been proposed as a strategy to suppress mutagenic processes and enhance cancer treatment efficacy.

**Figure 1.**
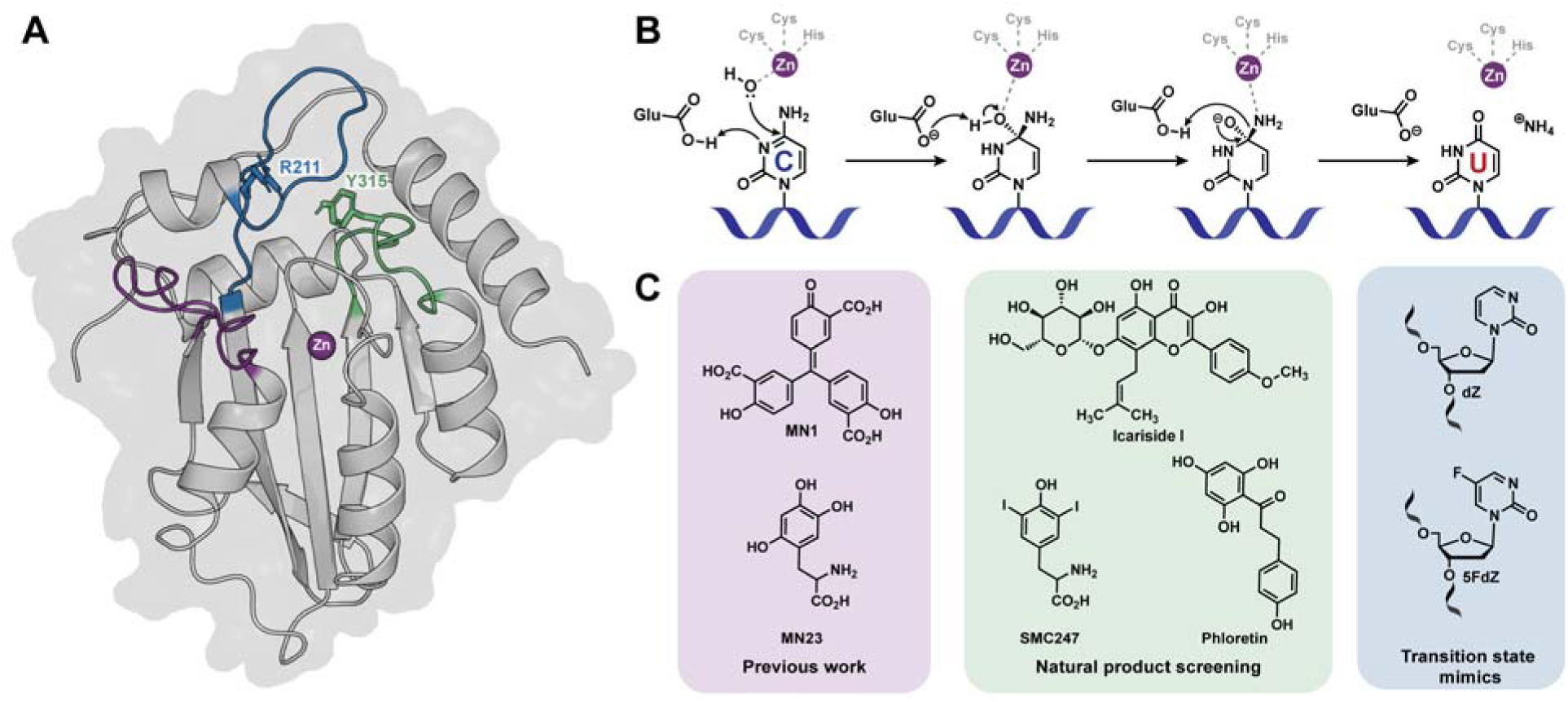
A) Structure of the c-terminal domain of A3B based on PDB ID 5TD5 with mutations reverted to wildtype. Loop one highlighted in blue, loop three highlighted in purple, and loop seven highlighted in green. Dynamic residues, R211 and Y315, implicated in the opening and closing of the active site are labeled. B) Mechanism of cytidine deamination by A3B. C) Previously reported inhibitors of A3B.

Despite its biological significance, A3B has remained a challenging target for inhibitor discovery. The conformational dynamics of the protein and high sequence homology across the A3 family has complicated the development of selective A3B inhibitors; more progress has been made in identifying pan-A3 inhibitors (**Figure 1C**).^11,12^ Previously, our group reported first-in-class small molecule inhibitors of APOBEC3G (A3G) identified through high-throughput screening, which also exhibited inhibitory activity against A3B (MN1 and MN23).^13^ However, these molecules possessed chemical liabilities and were highly reactive, precluding further investigation. Other groups have explored the potential of natural products for A3B inhibition, such as icariside, phloretin, and diiodotyrosine (SMC247); however, those molecules are not particularly suitable for drug development.^14, 15^ The incorporation of nucleosides that mimic the cytosine deamination transition state, such as zebularine, into nucleic acids have yielded ssDNA inhibitors with impressive potency.^16–19^ Such DNA-based approaches require further optimization to address cellular stability and delivery, which are common issues for oligonucleotide-based therapeutics.^20^

Small molecule A3B inhibitors with reasonable potency and specificity are needed to validate the mutagenic properties of the deaminase and to explore its potential as a therapeutic target. Consequently, we employed computer-aided drug discovery (CADD) to identify inhibitors of A3B. With the increase in computing power and improvements in algorithms, *in silico* screening can explore a larger chemical space than is feasible with traditional high-throughput screening. *In silico* modeling and docking provided the ability to screen a large library of molecules against a structure representative of wildtype A3B. In this study, we report the virtual screen of the ChemBridge Diversity Set library (110,000) against the active site in the c-terminal domain of A3B, as well as a novel putative allosteric site identified through computational methods. Following initial docking and scoring, and *in vitro* validation, several compounds appeared to inhibit A3B deaminase activity through a novel allosteric mechanism. However, subsequent resynthesis and analysis did not recapitulate the inhibition data produced from the secondary screen. Profiling of these (putative) A3B inhibitory compounds using several techniques coincided, at least with one compound, on palladium contamination in the screening sample that expels the zinc atom coordinated in the A3B active site. Nonetheless, mechanistic studies with CADD-identified ligands subsequently equipped with an electrophile result in the validation of a computationally identified allosteric binding site on A3B that is suitable for ligand-targeting.

## Results

### Molecular dynamics simulations of A3Bctd identify a putative allosteric site

We previously explored dynamics of wildtype A3Bctd, both in apo and single strand DNA (ssDNA)- bound forms, via MD simulations totaling 16 µs.^21^ In this prior work, the heavily mutated ssDNA-bound crystal structure (PDB ID 5TD5) was modeled back to wild type by reverting all mutations to the wild type sequence, as well as adding back in the deleted loop 3 residues. All A3Bctd snapshots from those MD simulations were clustered based on their active-site shape using the POVME3.0 program (**Figure 2A**).^22^ While the general structure is similar between the eight different clusters, differences can be observed in flexible loop regions, as well as in the positioning of the α-helices.

**Figure 2.**
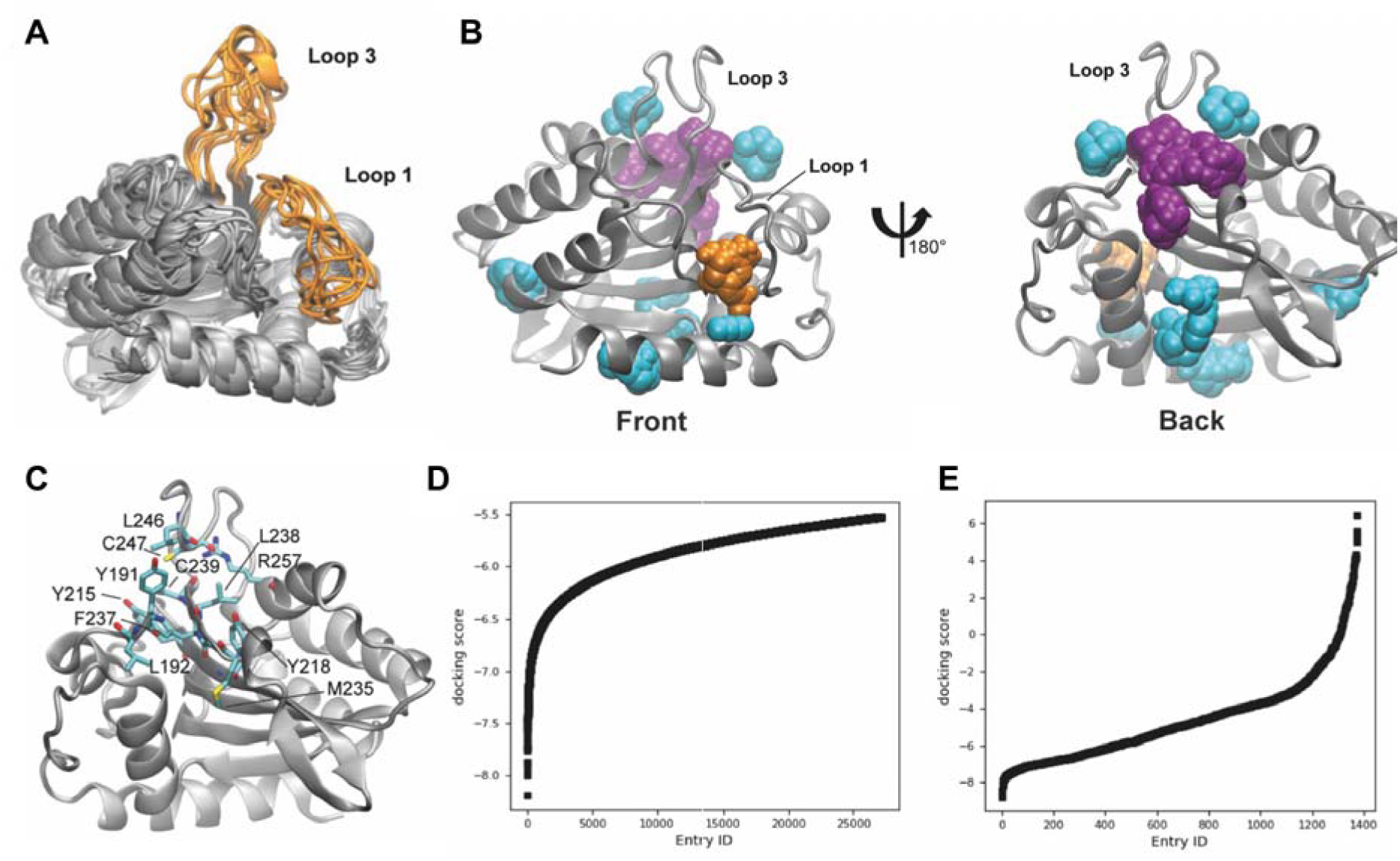
Molecular dynamics simulations and virtual screening of the A3Bctd active site and putative allosteric pocket. A) Eight cluster representative structures from MD simulations shown in silver ribbons, loops 1 and 3 highlighted in orange. B) Binding hot spots of Cluster6 representative structure depicted with FTMap probes as spheres. The front view highlights the FTMap probes binding into the active site, colored in orange. The back view highlights the location of the putative allosteric site, where the FTMap probes are colored in purple. C) Amino acids in 4 proximity of FTMap probes predicted to bind the putative allosteric site. This is the same view as the one shown on the right side of panel B. D) and E) Docking scores of compounds (in kcal/mol) from the virtual screen targeting the D) active site and E) putative allosteric site.

All eight representative A3Bctd snapshots – one from each of the eight clusters POVME3.0 detected – were then assessed for binding hot spots using FTMap, a computational solvent mapping program.^23^ FTMap utilizes small organic probe molecules and globally searches the protein surface for druggable sites. Consensus sites on the protein are locations where the highest number of probes dock. In addition to the A3Bctd active site (**Figure 2B**), a consensus site hot spot was detected by FTMap at the back of loop 3 distal to the active site (**Figure 2C**). FTProd, a program that compares and classifies FTMap-detected binding sites across multiple protein structures, identified this new site as a binding hot spot in 7 out of 8 cluster representative A3Bctd structures (**Figure S1**).^24^ Due to the challenge of finding small molecules that bind the active site of A3Bctd, we decided to assess both sites through virtual screening. Cluster0 and Cluster6 representative A3Bctd snapshots were selected for virtual screening as they had the first and second largest number of FTMap probes at the active site and the putative allosteric site, respectively.

Schrodinger Glide was employed to virtually screen the ChemBridge Diversity Set of 110,000 compounds for the active site (Cluster0) as well as the putative allosteric site (Cluster6).^25–27^ The compounds were ranked based on their docking scores and separately ranked on their ligand efficiency for the active site and putative allosteric site. Targeting the active site, 387 compounds were selected based on a docking score cutoff of -6.8 kcal/mol and another 311 compounds were selected with a ligand efficiency cutoff of -0.37 kcal/mol (**Figure 2D**). For the putative allosteric site, 712 compounds were selected using a docking score cutoff of -3.5 kcal/mol (**Figure 2E**). Further filtering based on compound availability resulted in a final set of 1408 compounds for *in vitro* testing in the secondary screen.

### *In vitro* screening yields thirteen primary hits

The 1408 compounds selected from docking were screened in biological duplicate at 500 µM in our established deaminase assay, using the previously characterized MN1 and MN23 as positive controls.^13^ A total of 23 small molecules inhibited A3Bctd deaminase activity below 30% in both biological replicates, resulting in a hit rate of approximately 2% (**Figure 3A**). Previous high-throughput screening campaigns against A3 enzymes generated hit rates between 0.02 - 0.8%,^28^ and we expected a higher hit rate than traditional HTS as compounds were pre-selected with a primary *in silico* screen. The plated compounds that elicited single-point responses were then tested in dose-response assays to further confirm activity (**Figure S2-3**). Of the 23 hits, 10 did not reconfirm in follow-up testing and three contained known PAINS-like moieties and were excluded,^29^ which yielded a total of 10 compounds for further study.

**Figure 3.**
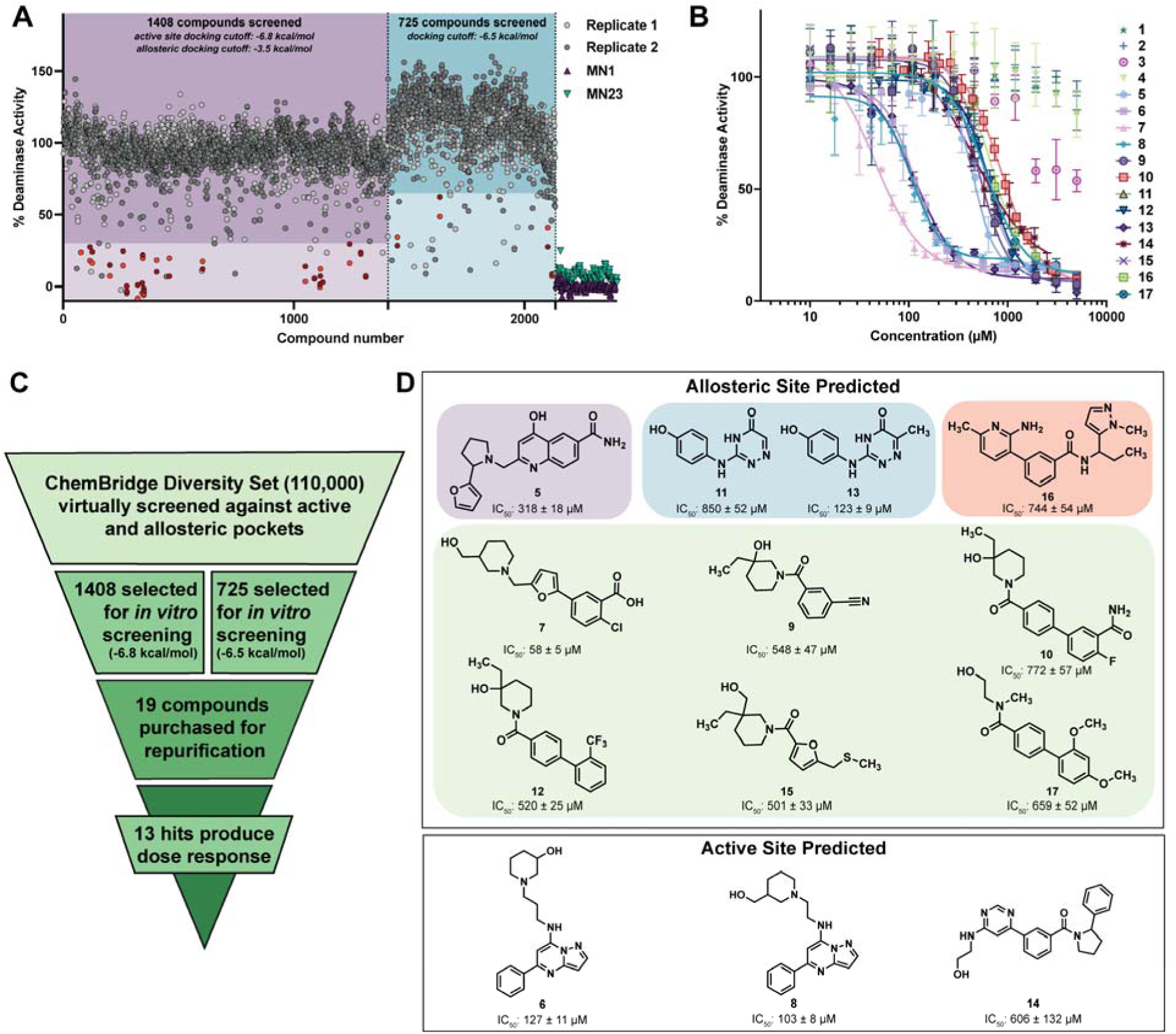
**A**) Primary *in vitro* screen in biological duplicate. First screen of 1408 compounds tested at 500 µM highlighted in purple, with light purple box indicating 30% deaminase activity cutoff. Second screen of additional 725 compounds tested at 100 µM highlighted in blue, with light blue box indicated 60% deaminase activity cutoff. MN1 and MN23 run as positive controls, indicated by green and purple triangles. **B**) Activity assay against A3Bctd after compound purification. Graph is one representative replicate of two biological replicates. Error bars are the standard error of the mean (SEM) of three technical replicates. **C**) Overview of screening triage to identify 13 hit compounds. **D**) Structure of 13 hit compounds with IC_50_ value below, grouped by predicted binding site, active site (bottom) and allosteric (top). Allosteric molecules are further grouped into representative chemotypes indicated by colored squares, with the most potent molecule from each group being selected for resynthesis (**5**, **7**, **13**, and **16**).

One of the first confirmed hits, **10**, had a docking score of -6.84 kcal/mol, which was low for the cutoff we selected (-6.8 to -8.1 kcal/mol), yet still performed well in the initial deaminase screen. Thus, we hypothesized that the original docking score cutoff may have been too stringent and promising scaffolds may have been excluded. Therefore, an additional 627 compounds with a lowered binding score cutoff of -6.5 kcal/mol were selected for testing, as well as an addition 98 compounds that were selected based on structural and 2D/3D similarity to **10** (**Figure 3A**). These additional 725 compounds were screened in the same manner, but at a lower concentration of 100 µM. Our goal was to identify more potent hits than those identified in the first screen, which yielded four additional hits that reduced deaminase activity below 60%. In aggregate, 19 compounds – 10 from the initial screen, four from the second screen, and five compounds discovered through SAR by catalogue against **10** – were all repurchased and the compounds were purified by preparative reverse-phase HPLC (**Table S3**). Two of these compounds were unstable or exhibited poor aqueous solubility, and were excluded, resulting in 17 *repurified hits* suitable for further analysis.

The repurified compounds were then re-tested in dose response, and 13 of the 17 small molecules retained inhibitory activity (**Figure 3B**). Of the four that did not reconfirm, three were compounds selected through “SAR by catalog” against **10** and were not tested in either of the initial screens. Potencies of the hits varied between 57 µM and 850 µM, with a majority of the compounds possessing triple digit µM IC_50_ values (**Figure 3D**). The FRET-based activity assay used to screen these compounds relies on the activity of two proteins to achieve a signal: A3B to deaminate the target cytosine to uracil, followed by uracil DNA glycosylase (UDG) to excise the uracil base. Addition of sodium hydroxide results in cleavage of the phosphodiester backbone and releases the FRET-quenching, producing a fluorescent signal. All 13 compounds were counter screened for UDG inhibition, as inhibition of UDG would produce the same result as A3B inhibition in this assay, and none of the 13 compounds exhibited any UDG inhibition (**Table S2**). Further investigation focused on identifying the mechanism of inhibition.

### A covalent analogue of inhibitor 10 adducts the allosteric pocket

With the computational identification of a novel allosteric binding site on A3B, considerable work went into validating the existence of this pocket. Most of the hits were predicted to bind at this allosteric site and results from competitive fluorescence polarization suggested that these molecules were not competing with DNA, supporting a binding mode distal to the active site (**Figure S4**). To probe the binding location of the allosteric molecules, we returned to the computational docking of **10** bound to A3Bctd, where we observed **10** positioned near two potentially solvent exposed cysteines, C239 and C247 (**Figure 4A**). We hypothesized that modification of **10** to possess an electrophilic warhead could validate its binding modality if covalent adduction to either cysteine was detected by mass spectrometry. Accordingly, we modified **10** to contain a chloroacetamide for cysteine reactivity, resulting in **24** (**Scheme 1**). Intact protein mass spectrometry revealed **24** adducts A3Bctd in a dose- and time-dependent manner with most of the protein being labeled at concentrations above 10X (**Figure S5**). Similar results were obtained against A3A, which is 92% identical in amino acid sequence to A3Bctd, with even higher levels of reactivity to C64 (C247 in A3B) (**Figure S6**). As expected, we observed a higher degree of labeling to both A3A and A3Bctd at the longer incubation times as covalent adduction is a time-dependent event.

**Figure 4.**
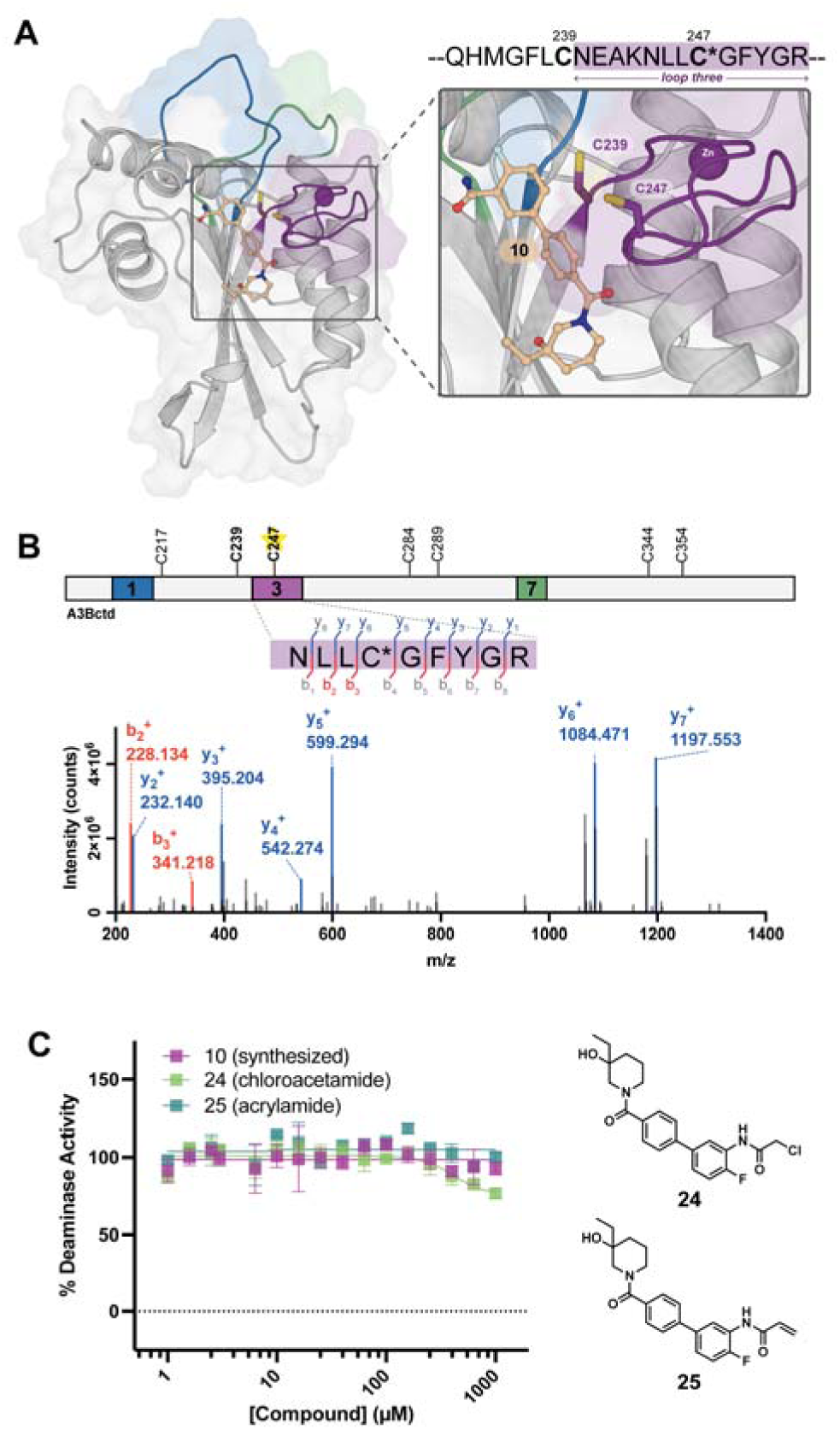
**A**) Model of **10** docked against A3Bctd (structure based on PDB ID 5TD5) with both solvent exposed cysteines labeled next to compound. Sequence of loop 3 above enlarged image of **10** docking model, with cysteines numbered and the primarily adducted cysteine (C247) indicated by the asterisk (**C\***). **B**) Collision-induced fragmentation pattern of peptides digested from A3Bctd treated with **24**. Precursor peptides encompass amino acids 244-252. The lines between the amino acids indicate the observed b- and y-ions. Sequence of the protein is depicted above with the DNA-engaging loops 1, 3, and 7, highlighted in colors matching the PDB structure with cysteines labeled. Cysteine adducted by **24** indicated by yellow star and asterisk. **C**) Activity assay of **10**, **24**, and **25** against A3Bctd. Structures of **24** and **25** depicted to right of graph. Graph is one representative replicate of two biological replicates. Error bars are the standard error of the mean (SEM) of three technical replicates.

**Scheme 1.**
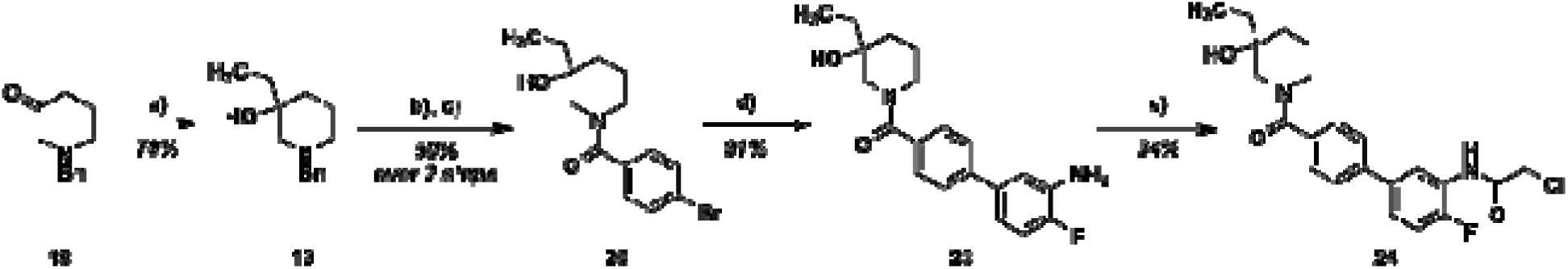
Synthesis of **24**. Reagents and Conditions: (a) EtMgBr, THF, 0 °C-rt; (b) H_2_, Pd/C, MeOH, 50 psi; (c) (4- Br)BzCl, Et_3_N, CH_2_Cl_2_; (d) 2-fluoro-5-(4,4,5,5-tetramethyl-1,3,2-dioxaborolan-2-yl)aniline, Pd(dppf)Cl_2_, K_3_PO_4_, EtOH, H_2_O, 80 °C; (e) chloroacetyl chloride, Et_3_N, CH_2_Cl_2_, °C-rt

To determine the specific site of adduction, A3Bctd was incubated with compound **24** and digested with trypsin for bottom-up proteomic analysis, where **24** was found to covalently label C247 (**Figure 4B**). This cysteine resides in the flexible region of loop 3 near the computationally predicted allosteric pocket, suggesting the compound may indeed be inhibiting through this novel allosteric mechanism. However, some additional adduction to C217 was observed (**Figure S8)**. Similar sites of adduction were detected in A3A digest experiments after treatment with compound **24** (**Figure S9** and **S10**). Using the previously described deaminase assay, we tested two covalent analogues of **10**, **24** (chloroacetamide) and **25** (acrylamide), against A3Bctd for inhibition. Covalent molecules commonly increase potency due to the inactivation of the protein by eliminating the rate of dissociation.^30^ Strikingly, **24** and **25** produced no significant inhibition of A3B. This unanticipated result prompted us to revisit the enzyme inhibitory properties of **10** using synthesized material (**Figure 4C**; synthesis in SI section 1.1). Our prior studies with **10** used compound that we purchased commercially, repurified by preparative reverse-phase HPLC, and characterized by mass spectrometry. Surprisingly, **10** did not recapitulate the inhibition data observed in the initial dose response assays; moreover, **10**, **24**, and **25** were also tested against A3A and no enzyme inhibition was observed. (**Figure S12**). We next focused on synthesizing representative chemotypes from the remaining allosteric hits to validate their allosteric inhibition properties observed from primary screens.

### Triage and resynthesis of representative chemotypes

Ten out of the thirteen small molecules identified from the primary screen were predicted to bind in the putative allosteric pocket. The allosteric molecules were grouped based on structural similarity, and the most potent hit from each chemotype was selected for resynthesis and testing (**Figure 3D**). Compounds **7**, **13**, and **16** were synthesized for follow up experiments. Notably, **7** is structurally similar to **10**, but was found to be 13-fold more potent in initial screens.

Synthesized **7**, **13**, and **16** were tested for inhibition against A3Bctd and compared to the original compound stocks (purchased from ChemBridge and repurified by preparative HPLC). Unfortunately, none of the synthesized compounds induced any significant inhibition of A3Bctd, while the original compound stocks produced IC_50_ values ranging between 60 - 178 µM (**Figure 5A**). The deaminase assay is a plate-based experiment that relies on the release of fluorescence quenching between TAMRA and AlexaFluor488 upon deamination by a cytidine deaminase. Fluorescence intensity is used to measure inhibition and thus compound stocks that interfere with fluorescence may produce a false positive result. To determine whether the purchased and repurified compound stocks were producing false positive results or genuinely inhibiting A3B, the resynthesized and purchased chemical stocks were tested in an orthogonal gel-based inhibition assay that we have previously reported.^31, 32^ Similar to the plate-based deaminase assay, the purchased and repurified stocks produced significant inhibition of A3B, while the synthesized compounds were inactive (**Figure 5B**). The stocks were also tested against endonuclease Q (a component of the gel-based deaminase assay) and no inhibition of enzyme activity was observed, indicating the positive results obtained with the commercial screening compounds were due to A3B inhibition and not due to another type of assay interference (**Figure 5C**).

**Figure 5.**
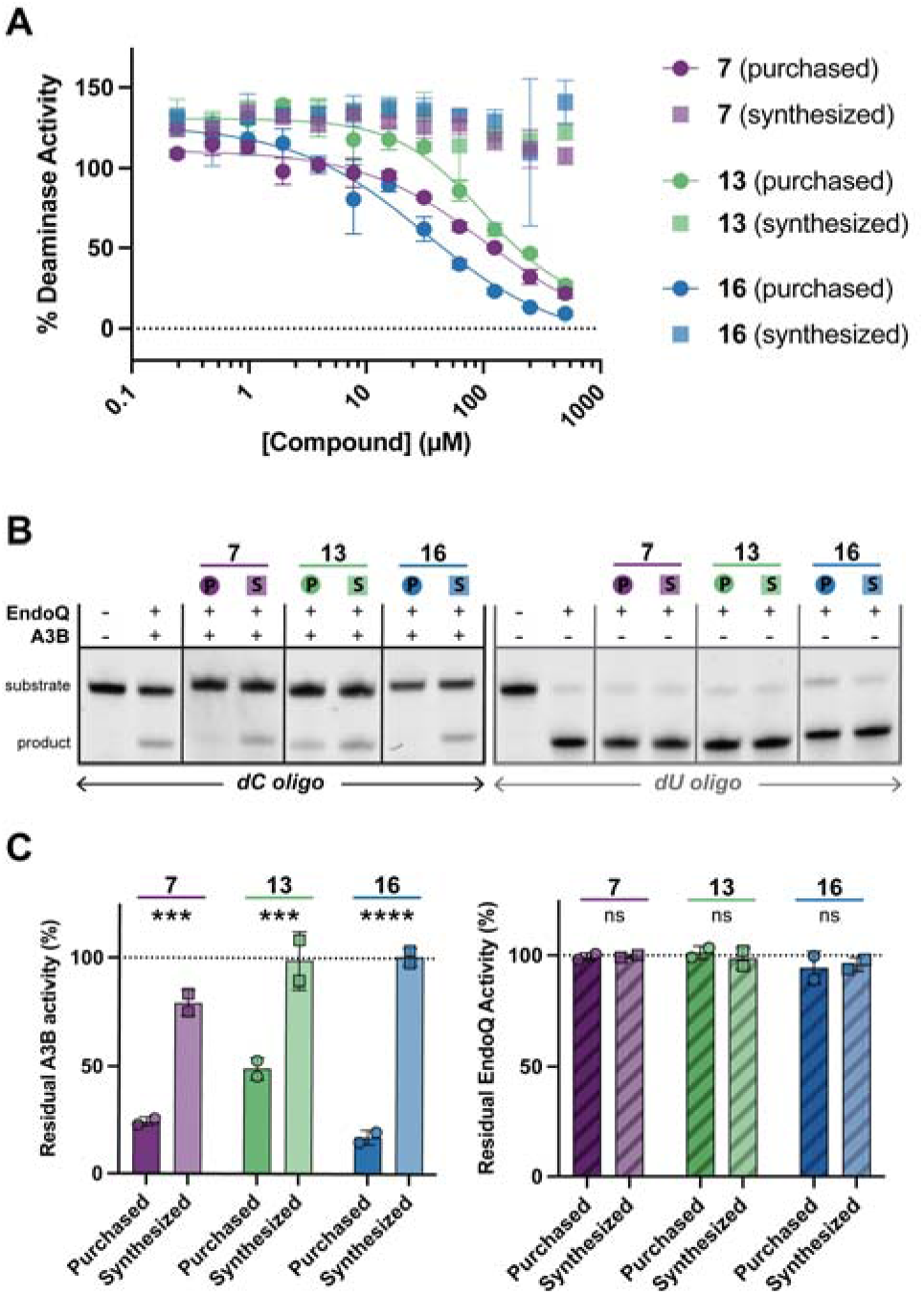
**A**) Plate-based deaminase assay of synthesized and purchased stocks of **7**, **13**, and **16** against A3Bctd. Graph is one representative replicate of two biological replicates. Error bars are the standard error of the mean (SEM) of three technical replicates. **B**) Fluorescently labeled DNA-substrates treated with A3B and Endo Q then separated by electrophoresis. Compound incubation (1 mM) samples are indicated by number above the lane, with purchased compounds indicated by the round symbol with ‘P’ and resynthesized stocks indicated by square symbol with ‘S’. Compound coloring matches previous graph (A). (left) Gel images of incubation with ‘TC’ substrate to measure A3B inhibition. (right) Gel images of incubation with pre-deaminated ‘TdU’ substrate to measure endonuclease Q inhibition. Uncropped gel images are available in supplementary information. (**C**) Bar plot of the percent residual activity of gel-images acquired in (B) and an additional biological replicate. The plot represents N = 2 biological replicates and error bars show standard deviation (SD). *** *p* < 0.001, **** *p* < 0.0001, ns = not significant.

### Metal contamination leads to inhibition of A3Bctd

The discrepancy in inhibition data of the synthesized and purchased compounds prompted the HPLC-(re)analysis of both stocks, which yielded peaks with the same retention times (**Figures S15-S23**). The synthesized versions of **7**, **13**, and **16** were further verified by NMR spectroscopy and mass spectrometry. With significant differences in biochemical potency between the stocks, and no detectable contaminants by HPLC, we hypothesized that non-UV active trace metal contaminants may be the culprit. Consequently, the purchased and synthesized stocks of **7**, **13**, and **16** were analyzed for metal contamination via inductively coupled plasma mass spectrometry (ICP-MS). Each sample was purified by preparative HPLC prior to analysis. Each stock was analyzed for cobalt, copper, zinc, palladium, platinum, and gold, as these are metals that can be used in the synthesis of hit molecules. None of the compounds possessed any significant levels of cobalt, platinum, or gold (**Table S13).** However, there were detectable amounts of copper, zinc and palladium (**Table 2**). The purchased stock of **7** exhibited higher levels of copper and zinc than its synthesized counterpart. Both the purchased and synthesized stocks of **13** possessed relatively low levels of all the metals analyzed and had higher levels in the synthesized stock, providing little insight into the source of inhibition. Conversely, **16**, the most potent compound in triage experiments, possessed 13 times more palladium contamination than the synthesized stock.

**Table 1.** Inhibition data for purchased and resynthesized compound stocks incubated at 1 mM. IC_50_ values are the mean of two biological replicates performed in technical triplicate, with propagated error as standard error of the mean (SEM). Average percent residual activity from gel-based deaminase from two biological replicates with propagated error as standard deviation (SD).

| Compound | Stock origin | IC<sub>50</sub> (plate-based) | Percent residual activity (gel-based) |
| --- | --- | --- | --- |
| 7 | Purchased | 94.1 ± 46 μM | 24 ± 2% |
|  | Synthesized | n.d. | 80 ± 6% |
| 13 | Purchased | 177 ± 44 μM | 49 ± 5% |
|  | Synthesized | n.d. | 99 ± 13% |
| 16 | Purchased | 59.7 ± 13 μM | 17 ± 3% |
|  | Synthesized | n.d. | 100 ± 4% |

**Table 2.**
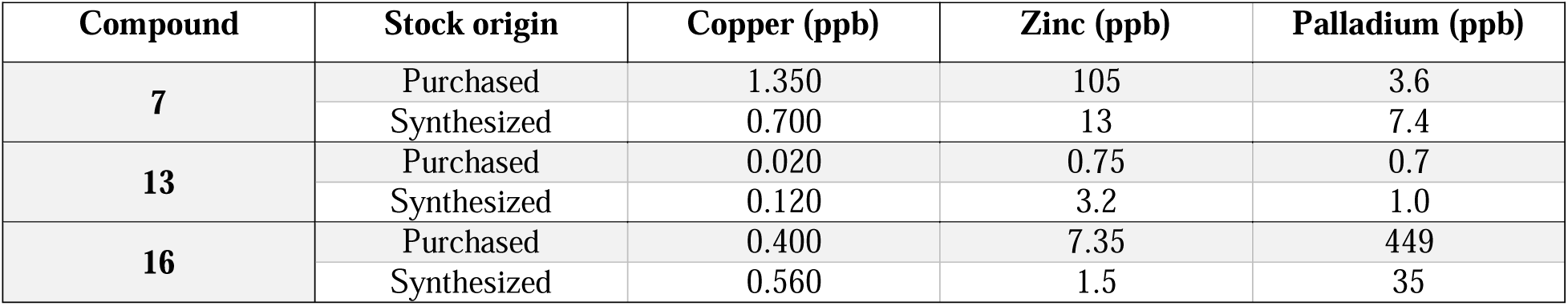
ICP-MS analysis of purchased and resynthesized stocks of **7**, **13**, and **16** scaled for 1 mM compound concentration reported in parts per billion (ppb).

| Compound | Stock origin | Copper (ppb) | Zinc (ppb) | Palladium (ppb) |
| --- | --- | --- | --- | --- |
| 7 | Purchased | 1.350 | 105 | 3.6 |
|  | Synthesized | 0.700 | 13 | 7.4 |
| 13 | Purchased | 0.020 | 0.75 | 0.7 |
|  | Synthesized | 0.120 | 3.2 | 1.0 |
| 16 | Purchased | 0.400 | 7.35 | 449 |
|  | Synthesized | 0.560 | 1.5 | 35 |

In the interest of identifying the source causing the observed A3Bctd inhibition, all three detectable metals were tested against A3Bctd in the plate-based deaminase assay. Neither CuCl_2_ nor ZnCl_2_ significantly inhibited A3Bctd, while Pd(OAc)_2_ potently inhibited the protein at 72 ppb (IC_50_ = 320 nM) (**Figure 6A**). These metals were additionally tested against A3A where similar results were obtained; however, A3A is even more sensitive to palladium (31 ppb, IC_50_ = 139 nM) (**Figure S24**). Considering that purchased **16** has a Pd(II) concentration more than 6-fold higher than the concentration needed to inhibit the protein to 50% deaminase activity, the inhibition observed in the primary and triage experiments may be due to the trace palladium contamination. Considering these samples were purified prior to ICP-MS and deaminase testing, the original concentrations in the purchased powders were likely much higher.

**Figure 6.**
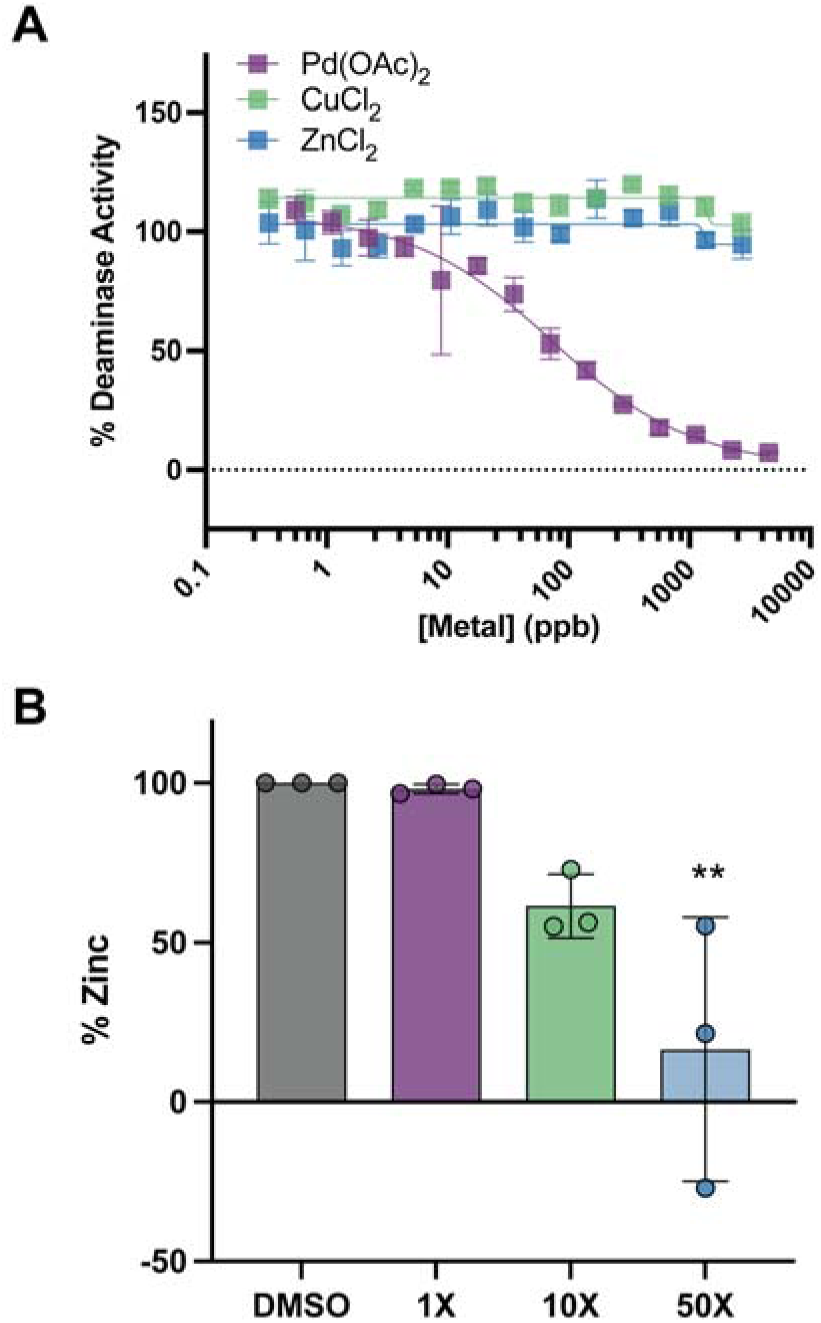
**A**) Plate-based deaminase assay of Pd(OAc)_2_, CuCl_2_, and ZnCl_2_ against A3Bctd. Graph is one representative replicate of two biological replicates. Error bars are the standard error of the mean (SEM) of three technical replicates. **B**) Bar plot of the colorimetric zinc detection assay (Abcam) against A3Bctd in the presence of increasing ratios of Pd(OAc)_2_. The plot represents N = 3 biological replicates, each tested in technical duplicate, and error bars show standard deviation (SD). ** *p* < 0.01.

We were next interested in exploring the mechanism of cytidine deaminase inhibition by Pd(II). Previous studies have discovered that platinum and palladium metal complexes may expel zinc ions coordinated by cysteines in zinc finger proteins.^33,34^ We hypothesized that palladium may be acting similarly against A3B, as A3 enzymes have a catalytic zinc coordinated by two cysteines and a histidine. To analyze this, the levels of zinc in A3Bctd were measured in the presence of increasing amount of Pd(OAc)_2_ using a colorimetric assay. Concentrations higher than 10X and 50X resulted in a decrease in the amount of zinc detected in A3Bctd (**Figure 6B**). Removal of the catalytic zinc by palladium contamination should result in enzyme inhibition as the zinc is required for catalysis (**Figure 1B**). These results support that the palladium contamination present in purchased **16** is responsible for the observed inhibition of A3Bctd. However, these results do not provide insight into the source of inhibition from the purchased stocks of **7** or **13**. Due to their high purity observed by HPLC, we hypothesize that other non-UV active contaminants may be contributing to their false-positive behavior. However, more importantly, our results indicate that neither **7** nor **13** are A3Bctd inhibitors.

## Conclusions

In this study, a CADD approach was taken to identify small molecule inhibitors of A3Bctd, with a focus on active- and allosteric-site-targeting compounds. Unfortunately, none of the synthesized molecules recapitulated the inhibition data observed in the primary activity screen. Although the molecules identified in this study cannot be utilized to explore the computationally identified allosteric pocket; a representative covalent inhibitor based on **10**, compound **24**, was found to adduct C247 in the computationally predicted allosteric pocket, which provides a rationale for exploration into other covalent molecules to target this region of A3B. This presence of an allosteric pocket is further supported by the human cysteine database that reports the only druggable cysteine in A3B is C247, which is the residue covalently modified by compound **24**.^35^ The reproducibility of this site across multiple molecular dynamics simulations supports its plausibility and motivates further investigation as allosteric inhibition may prove as an effective method for the development of selective A3B inhibitors.

This study also highlights several methodological challenges relevant to research with A3 enzymes, or metalloproteins, in general. As is the case with many computational pipelines, our docking prioritization relied on the static representations of a dynamic enzyme. More recent computational work has reported that the N- and C-terminal domains may interact to open and close the active site of A3B.^11^ Our work only examined the C-terminal domain and utilized a heavily mutated crystal structure^32^ where the wildtype residues were reverted computationally. Additionally, our study highlights the sensitivity of A3B to trace metal contamination, which may disrupt the coordination of the catalytic zinc. This reveals a unique vulnerability of zinc-dependent metalloproteins, and drug hunters pursuing these targets should always resynthesize their hit compounds and be wary of contamination not readily detected by HPLC or NMR. However, this is not a new phenomenon as metal contamination and false positive results have consistently challenged screening projects.^36–38^ Taken together, although no validated A3B inhibitors were identified in this study, we nonetheless report the first validated allosteric site on A3B that is amenable for covalent targeting, which is a focus on ongoing studies by our team.

## Supporting Information

Additional supplemental experimental data (**Figures S1-S26**, **Tables S1-S13**), experimental methods, synthetic procedures, and NMR characterization spectra are available.

## Acknowledgements

This work was supported by the National Institutes of Health, National Cancer Institute (P01-CA234228 to RSH, REA, DAH). Funding for M.K.W. was provided by the University of Minnesota Doctoral Dissertation Fellowship (UMN DDF) Program. K.M.F.J. was funded by the National Science Foundation Graduate Research Fellowship Program (NSF GRFP) and UMN DDF. C.K.M. was supported by an NIH NCI traineeship (T32-CA009523). Mass spectrometry was performed at the University of Minnesota Masonic Cancer Center, Analytical Biochemistry Core Facility, which is supported by the NIH (P30- CA77598).

## Abbreviations

APOBEC3: Apolipoprotein B mRNA Editing Catalytic Polypeptide-like Subunit 3, A3
C: Cytidine
CADD: Computer Aided Drug Discovery
dC: 2′-Deoxycytidine
dU: 2′-Deoxyuridine
EndoQ: Endonuclease Q
FRET: Förster Resonance Energy Transfer
HPLC: High Performance Liquid Chromatography
ICP-MS: Inductively Coupled Plasma Mass Spectrometry
MD: Molecular Dynamics
PAINS: Pan-Assay Interference Compounds
T: Thymidine
U: Uridine
UDG: Uracil DNA Glycosylase

