## Supplemental Information for "Computational Discovery of an Allosteric Pocket on APOBEC3B and the Impact of Metal Contamination on Inhibitor Validation"

### 1 Methods

#### 1.1 Synthesis

Unless otherwise noted, reactants and glassware were not dried before use, and reactions were stirred with a Teflon-coated stir bar. Reaction solvents acetonitrile ($CH_3CN$), dichloromethane ($CH_2Cl_2$), and tetrahydrofuran (THF) were dried by passage over a column of activated alumina using a solvent purification system (MBraun). Dimethylformamide (DMF) was dried by passage over a column of molecular sieves using a solvent purification system (MBraun). Reactions were monitored using thin layer chromatography with EMD Chemicals Silica Gel 60 $F_{254}$ glass plates (250 μm thickness) and visualized with UV irradiation at either 254 nm or 365 nm. Flash chromatography was performed with a Teledyne-Isco CombiFlash NextGen instrument, equipped with both a UV-Vis detector and an ELSD, using Redisep Rf High Performance silica gel columns (Teledyne-Isco). $^1$H NMR (500 MHz) and $^{13}$C NMR (125 MHz) were collected on a Bruker Advance NMR spectrometer at room temperature. NMR chemical shifts (δ) are recorded relative to TMS (0.05% v/v, δ = 0.0) or residual solvent signal (δ = 7.26 ppm for $CDCl_3$, δ = 2.50 ppm for DMSO-*d*$_6$, or δ = 3.31 ppm for MeOD) for $^1$H NMR and the solvent signal for $^{13}$C NMR (δ = 77.0 for $CDCl_3$, δ = 40.0 for DMSO-*d*$_6$, or δ = 49.0 ppm for MeOD). $^1$H peak broadening and additional $^{13}$C resonances are observed for compounds **23**, **24**, and **25**, which are attributed to conformational restriction of the substituted piperidine. This observation is consistent with a prior report.[1] HRMS data were collected on an Orbitrap Elite (Thermo) mass analyzer at 240,000 resolution with electrospray ionization in positive mode. Reverse-phase purification was performed on a preparative-scale Agilent 1200 series instrument. The preparative column was an Agilent Zorbax SB-C18 (21.2 x 250 mm, 7 μm pore). Mobile phase A = $H_2O$ with 0.1% TFA, mobile phase B = MeCN with 0.1% TFA, flow rate = 30 mL/min. Elution: 90:10 A:B for 0-2 min, gradient to 30:70 A:B from 2-22 min, gradient to 5:95 A:B from 22 min to 28 min, isocratic at 5:95 A:B from 28-30 min.

*1.1.1 Synthesis of 1-benzyl-3-ethylpiperidin-3-ol (**19**)*

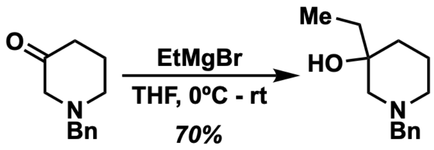


**18** **19**

To a suspension of *N*-benzyl-3-piperidone HCl salt (**18**, 4.51 g, 20.0 mmol) in anhydrous THF (40 mL) in a flame-dried flask was added EtMgBr (1.0 M in THF, 50 mL, 50 mmol) slowly at 0°C under $N_2$ and then warmed to room temperature overnight. Saturated aqueous $NH_4Cl$ solution (75 mL) was carefully added, and the reaction mixture was concentrated *in vacuo* to remove the THF. The crude material was diluted with water (50 mL) and extracted with $CH_2Cl_2$ (50 mL x 3), then the combined organic layers were dried over $Na_2SO_4$, concentrated *in vacuo* then purified by flash chromatography (0-50% EtOAc in hexanes gradient) resulting in a pale-yellow oil, **19** (2.41 g, 11.0 mmol) in 70% yield.

**$^1$H NMR (500 MHz, $CDCl_3$):** δ 7.36 – 7.21 (m, 5H), 3.62 – 3.44 (m, 2H), 3.24 (s, 1H), 2.79 (d, *J* = 11.0 Hz, 1H), 2.59 (dt, *J* = 10.9, 2.2 Hz, 1H), 2.03 – 1.85 (m, 2H), 1.77 (qt, *J* = 12.7, 4.3 Hz, 1H), 1.67 – 1.58 (m, 1H), 1.54 (ddq, *J* = 13.5, 5.6, 2.8 Hz, 1H), 1.45 (qd, *J* = 7.5, 4.8 Hz, 2H), 1.20 (td, *J* = 13.0, 4.9 Hz, 1H), 0.90 (t, *J* = 7.6 Hz, 3H).

**HRMS:** $C_{14}H_{22}NO$ Calc'd $[M+H]^+$: 220.1701, Found: 220.1686.

*1.1.2 Synthesis of 1-(4-bromobenzoyl)-3-ethylpiperidin-3-ol (**20**)*

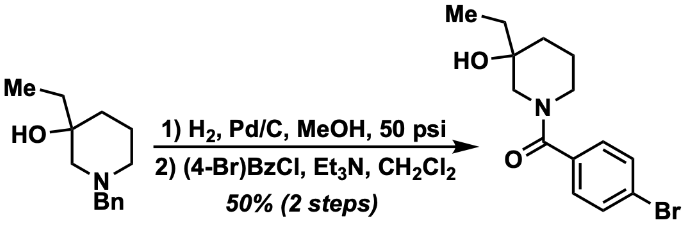


**19** **20**

To a solution of **19** (2.41 g, 11.0 mmol) in MeOH (55 mL, 0.2 M) was added Pd/C (10% w/w) then the mixture was degassed with $N_2$ and then charged $H_2$. The mixture was then shaken under 50 psi $H_2$ overnight on a Parr hydrogenator. The mixture was filtered through Celite and concentrated *in vacuo*, yielding a pale-yellow oil. The crude product (782 mg, 6.05 mmol) was used in the next step without further purification.

The crude intermediate was dissolved in anhydrous $CH_2Cl_2$ (12.1 mL, 0.5 M), then $Et_3N$ (1.27 mL, 9.08 mmol) and 4-bromobenzoyl chloride (1.46 g, 6.66 mmol) was added. The solution was stirred overnight under $N_2$ then concentrated *in vacuo*. The resulting crude material was taken up in EtOAc (25 mL), washed with saturated aqueous $NH_4Cl$ solution (20 mL), water (20 mL), and brine (20 mL). The organic layer was dried over $Na_2SO_4$, concentrated *in vacuo*, and purified by flash chromatography (0-10% MeOH in $CH_2Cl_2$) yielding the product, **20**, as a white solid (545 mg, 1.75 mmol) in 50% yield over two steps.

**$^1$H NMR (500 MHz, $CD_3OD$):** δ 7.61 (dd, *J* = 13.3, 7.9 Hz, 2H), 7.37 (dd, *J* = 25.6, 8.0 Hz, 2H), 4.08 (dd, *J* = 162.9, 13.0 Hz, 1H), 3.41 (dd, *J* = 37.7, 13.5 Hz, 1H), 3.29 – 3.02 (m, 2H), 1.84 (dd, *J* = 47.3, 12.2 Hz, 1H), 1.76 – 1.68 (m, 1H), 1.68 – 1.45 (m, 3H), 1.45 – 1.24 (m, 1H), 0.89 (dt, *J* = 93.7, 7.6 Hz, 3H).

**$^{13}$C NMR (126 MHz, $CD_3OD$):** δ 170.9, 170.5, 135.2, 134.9, 131.5, 131.2, 129.2, 128.3, 123.4, 69.7, 69.5, 56.8, 51.0, 42.5, 34.2, 34.1, 31.6, 31.6, 21.6, 20.4, 6.0, 5.7 (note: extra peaks are due to conformational restrictions).

**HRMS:** $C_{14}H_{18}BrNO_2$ Calc'd $[M+H]^+$: 312.0599, Found: 312.0585.

#### *1.1.3 Synthesis of 4'-(3-ethyl-3-hydroxypiperidine-1-carbonyl)-4-fluoro-[1,1'-biphenyl]-3-carboxamide (**10***)*

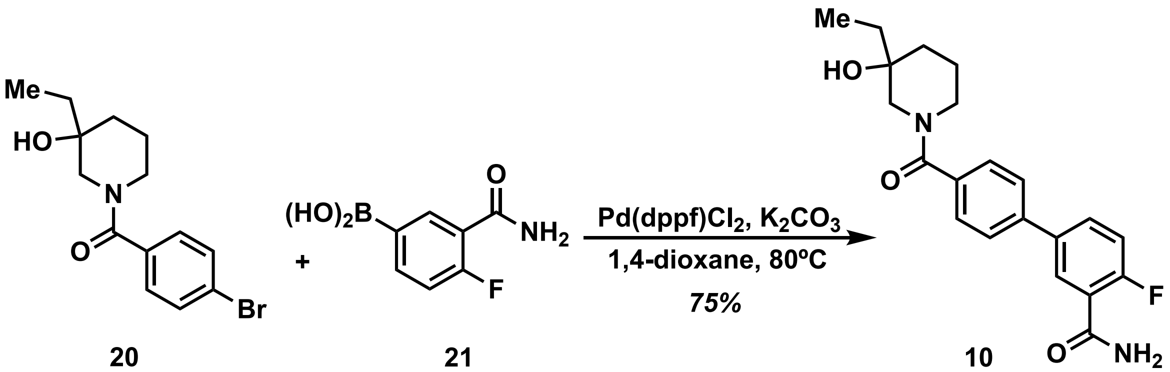


**20** **21** **10**

A solution of **20** (187 mg, 0.450 mmol), (3-carbamoyl-4-fluorophenyl)boronic acid (**21**, 90.6 mg, 0.495 mmol), and $K_2CO_3$ (5 M in $H_2O$, 270 µL) in 1,4-dioxane (20 mL, 0.02 M) was degassed by placing the flask under vacuum and purging several times with $N_2$, then $Pd(dppf)Cl_2$ (36.7 mg, 0.045 mmol) was added and the mixture heated to 60°C overnight. The dioxane was evaporated *in vacuo*, and the resulting crude material was diluted with EtOAc (20 mL) and water (20 mL), then filtered through celite. The organic layer was washed with water (15 mL) and brine (20 mL), dried over $Na_2SO_4$, concentrated *in vacuo* and then purified by flash chromatography (0-100%

EtOAc in hexanes) yielding a light brown foamy solid. The product was further purified by reverse-phase preparative HPLC (described in General) yielding the final product, **10**, (125 mg, 0.337 mmol) as a foamy white solid in 75% yield.

**$^1$H NMR (500 MHz, DMSO-*d*$_6$):** δ 7.92 - 7.90 (m, H), 7.85 – 7.82 (m, 2H), 7.73 – 7.68 (m, 3H), 7.52 – 7.46 (m, 2H), 7.39 – 7.35 (m, 1H), 4.44 – 4.39 (m, 1H), 3.97 (bs, 1H), 3.30 – 3.07 (m, 4H), 1.73 – 1.19 (m, 6H), 0.88 – 0.69 (m, 3H).

**$^{13}$C NMR (126 MHz, DMSO-*d*$_6$):** δ 165.3, 160.2, 158.3, 135.8 (d, $J_{C\text{-}F}$ = 12.2 Hz), 130.8 (d, $J_{C\text{-}F}$ = 32.4 Hz), 128.4, 127.5, 126.7, 126.4, 124.3 (d, $J_{C\text{-}F}$ = 54.1 Hz), 117.1 (d, $J_{C\text{-}F}$ = 85.3 Hz), 68.8, 56.5, 51.0, 42.1, 34.5, 31.4, 20.7, 7.0.

**$^{19}$F NMR (471 MHz, CDCl$_3$):** δ -115.30

**HRMS:** $C_{21}H_{23}FN_2O_3$ Calc'd $[M+H]^+$: 371.1771, Found: 371.1757.

*1.1.4 Synthesis of 1-{3'-amino-4'-fluoro-[1,1'-biphenyl]-4-carbonyl}-3-ethylpiperidin-3-ol (**23**)*

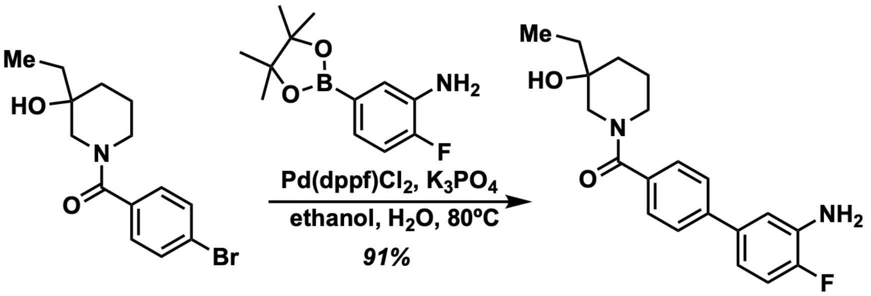


**20** **23**

A solution of 1-(4-bromobenzoyl)-3-ethylpiperidin-3-ol (**20**, 150 mg, 0.480 mmol), 2-fluoro-5-(4,4,5,5-tetramethyl-1,3,2-dioxaborolan-2-yl)aniline (114 mg, 0.480 mmol), and $K_3PO_4$ (1.0 M in $H_2O$, 1.44 mmol) in ethanol (2.5 mL) was degassed by placing under vacuum and purging several times with $N_2$. Next, Pd(dppf)$Cl_2$ (19.6 mg, 0.024 mmol) was added, and the reaction was heated to reflux for 4 hours. After this time, the reaction mixture was cooled to room temperature, diluted with water (10 mL) then extracted with $CH_2Cl_2$ (15 mL, 3x). The combined organics were washed with saturated brine (10 mL), dried over $Na_2SO_4$, concentrated *in vacuo*, and purified on a silica gel column (0-20% MeOH in $CH_2Cl_2$) yielding 1-{3'-amino-4'-fluoro-[1,1'-biphenyl]-4-carbonyl}-3-ethylpiperidin-3-ol (**23**) as an off-white foamy solid in 91% yield (150 mg, 0.438 mmol).

**$^1$H NMR (500 MHz, CDCl$_3$):** δ 7.53 - 7.46 (m, 4H), 7.04 – 7.97 (m, 2H), 6.89 – 6.86 (m, 1H), 4.33 (bs, 1H), 3.70 – 3.51 (bd, 3H), 3.10 – 3.00 (m, 2H), 1.91 – 1.83 (m, 1H), 1.76 (bs, 1H), 1.51 (bs, 4H), 0.92 (bs, 3H).

**$^{13}$C NMR (126 MHz, CDCl$_3$):** δ 171.5, 151.5 (d, $J_{C\text{-}F}$ = 240 Hz), 141.7, 136.7 (d, $J_{C\text{-}F}$ = 2.5 Hz), 134.5, 134.4 (d, $J_{C\text{-}F}$ = 13.8 Hz), 127.5, 126.7, 117.3 (d, $J_{C\text{-}F}$ = 7.5 Hz), 115.6 (d, $J_{C\text{-}F}$ = 3.7 Hz) , 115.4 (d, $J_{C\text{-}F}$ = 19.0 Hz), 70.3, 56.9, 51.8, 48.4, 42.6, 34.5, 32.9, 31.7, 22.1, 20.8, 6.9 (note: extra peaks are due to conformational restrictions).

**$^{19}$F NMR (471 MHz, CDCl$_3$):** δ -136.5.

**HRMS:** $C_{21}H_{23}FN_2O_3$ Calc'd [M+H]$^+$: 343.1822, Found: 343.1806.

*1.1.5 Synthesis of N-{4'-[(3-ethyl-3-hydroxy-1-piperidyl)carbonyl]-4-fluoro-3-biphenylyl}chloroacetamide (**24**)*

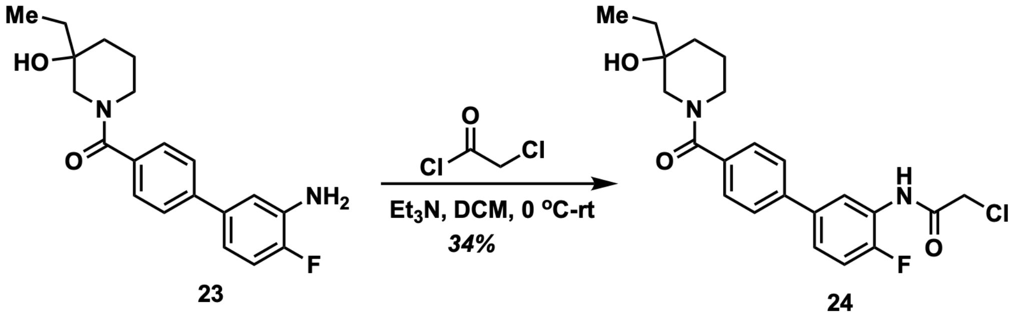


To a solution of 1-{3'-amino-4'-fluoro-[1,1'-biphenyl]-4-carbonyl}-3-ethylpiperidin-3-ol (**23**, 150 mg, 0.438 mmol) in $CH_2Cl_2$ (3 mL) was added $Et_3N$ (79 µL, 0.569 mmol). The mixture was cooled to 0 °C and to the reaction mixture was added chloroacetyl chloride (41.8 µL, 0.526 mmol) in $CH_2Cl_2$ (1 mL). The reaction mixture was stirred and allowed to warm to room temperature for 2 h. The reaction was monitored by TLC and upon consumption of starting material, the reaction mixture was concentrated *in vacuo*. The residue was redissolved in EtOAc (20 mL) and washed with water (15 mL) and brine (10 mL). The organic layer was dried over $Na_2SO_4$, concentrated in *vacuo* and then purified by flash chromatography (0-100% EtOAc in hexanes). The product was further purified by reverse-phase HPLC, as detailed in the General section, yielding N-{4'-[(3-ethyl-3-hydroxy-1-piperidyl)carbonyl]-4-fluoro-3-biphenylyl}chloroacetamide (**24**) as a white solid in 34% yield (62 mg, 0.148 mmol).

**$^1$H NMR (500 MHz, CD$_3$OD):** δ 10.2 (s, 1H), 8.22 (d, *J* = 5.5 Hz, 1H), 7.66 (d, *J* = 7.3 Hz, 2H), 7.53 – 7.47 (m, 3H), 7.41 – 7.38 (m, 1H), 4.39 (s, 2H), 4.34 (bs, 1H), 3.97 (bs, 0.5H), 3.54 (bs, 0.5H), 3.32 (bs, 1H), 3.14 - 3.05 (m, 1H), 1.75 – 1.21 (m, 6H), 0.89 (s, 1H), 0.70 (s, 2H).

**$^{13}$C NMR (126 MHz, DMSO-d$_6$):** δ 169.3, 168.9, 165.3, 153.4 (d, $J_{C\text{-}F}$ = 248 Hz), 139.6, 135.9 (d, $J_{C\text{-}F}$ = 3.7 Hz), 135.6, 128.2, 127.3, 126.5, 126.2, 125.9 (d, $J_{C\text{-}F}$ = 12.6 Hz), 124.1 (d, $J_{C\text{-}F}$ = 7.5

Hz), 122.3, 116.2 (d, $J_{C\text{-}F}$ = 20.1 Hz), 68.6, 56.4, 50.9, 47.4, 43.1, 41.9, 34.9, 32.2, 31.0, 21.8, 20.6, 7.1, 6.9 (note: extra peaks are due to conformational restrictions).

**$^{19}$F NMR (471 MHz, $CDCl_3$):** δ -132.3.

**HRMS:** $C_{22}H_{24}ClFN_2O_3$ Calc'd $[M+H]^+$: 419.1537, Found: 419.1523.

*1.1.6 Synthesis of N-(4'-(3-ethyl-3-hydroxypiperidine-1-carbonyl)-4-fluoro-[1,1'-biphenyl]-3-yl)acrylamide (**25**)*

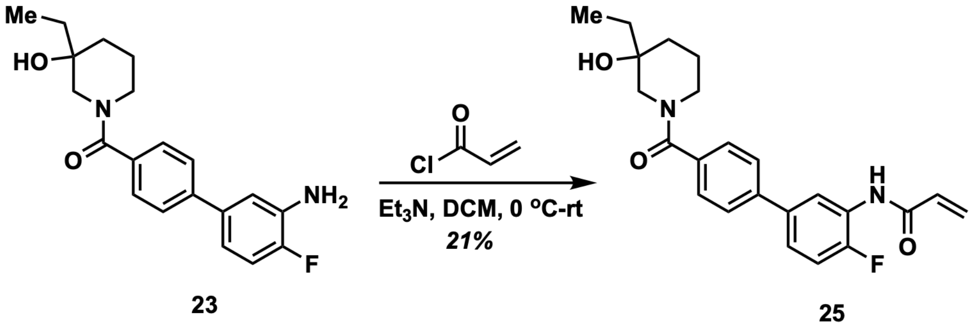


To a solution of 1-{3'-amino-4'-fluoro-[1,1'-biphenyl]-4-carbonyl}-3-ethylpiperidin-3-ol (**23**, 200 mg, 0.584 mmol) in $CH_2Cl_2$ (3 mL) was added $Et_3N$ (106 µL, 0.759 mmol). The mixture was cooled to 0°C and to the reaction mixture was added acryloyl chloride (56.6 µL, 0.701 mmol) in $CH_2Cl_2$ (1 mL). The reaction mixture was stirred and allowed to warm to room temperature for 2 h. The reaction was monitored by TLC and upon consumption of starting material, the reaction mixture was concentrated *in vacuo*. The residue was redissolved in EtOAc (20 mL) and washed with water (15 mL) and brine (10 mL). The organic layer was dried over $Na_2SO_4$, concentrated *in vacuo* and then purified by flash chromatography (0-100% EtOAc in hexanes). The product was further purified by reverse-phase C18 column chromatography (10-95% MeCN in water) yielding N-[4'-(3-ethyl-3-hydroxypiperidine-1-carbonyl)-4-fluoro-[1,1'-biphenyl]-3-yl]prop-2-enamide (**25**) as a white solid in 21% yield (48 mg, 0.121 mmol).

**$^1$H NMR (500 MHz, $CDCl_3$):** δ 8.69 (d, *J* = 5.9 Hz,1H), 7.62 (s, 1H), 7.59 (d, *J* = 8.0 Hz, 2H), 7.49 (d, *J* = 7.9 Hz, 2H), 7.29 – 7.26 (m, 1H), 7.18 – 7.14 (m, 1H), 6.48 (d, *J* = 16.8 Hz, 1H), 6.35 – 6.30 (m, 1H), 5.83 (d, *J* = 10.2 Hz, 1H), 4.33 (bs, 1H), 3.70 – 3.52 (m, 1H), 3.11 – 3.01 (m, 2H), 2.10 – 1.53 (m, 8H), 0.99 – 0.85 (m, 3H).

**$^{13}$C NMR (126 MHz, $CDCl_3$):** δ 171.6, 171.1, 163.6, 152.4 (d, $J_{C\text{-}F}$ = 245 Hz), 141.2, 135.7 (d, $J_{C\text{-}F}$ = 248 Hz), 130.7, 128.3, 127.5, 126.9, 126.3 (d, $J_{C\text{-}F}$ = 10.0 Hz), 123.0 (d, $J_{C\text{-}F}$ = 6.3 Hz), 120.9,

115.1 (d, $J_{C-F}$ = 20.1 Hz), 70.4, 60.3, 57.1, 53.3, 51.8, 48.4, 42.6, 34.5, 32.8, 31.8, 29.6, 22.2, 20.9, 14.1, 6.9 (note: extra peaks are due to conformational restrictions).

**$^{19}$F NMR (471 MHz, CDCl3):** δ -133.1.

**HRMS:** $C_{23}H_{25}FN_2O_3$ Calc'd $[M+H]^+$: 397.1928, Found: 397.1910.

*1.1.7 Synthesis of (1-((5-bromofuran-2-yl)methyl)piperidin-3-yl)methanol (**28**)*

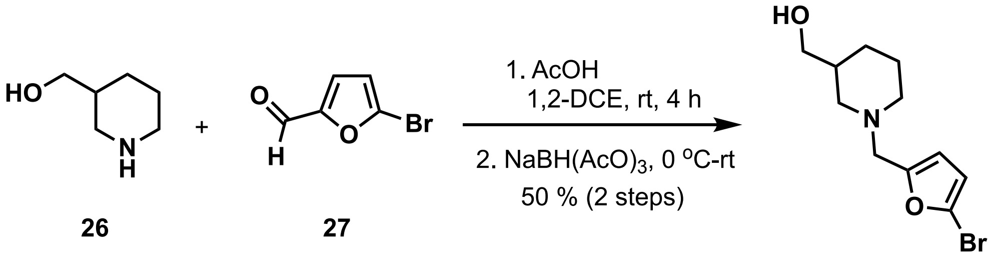


26 27 28

To a mixture of (3-piperidyl)methanol (**26**, 987 mg, 8.57 mmol) and 5-bromo-2-furaldehyde (**27**, 1.50 g, 8.57 mmol) in 1,2-dichloroethane (20 mL) was added acetic acid (pH ~4.5) and the resulting mixture was stirred at room temperature for 4 h. After this time, the reaction mixture was cooled to 0°C and was added sodium triacetoxyborohydride (2.18 g, 10.3 mmol). The reaction mixture was allowed to warm to room temperature and stirred further for 17 h. Then, the reaction mixture was cooled to 0 °C and carefully quenched with water (5 mL) and neutralized with saturated aqueous sodium bicarbonate solution (50 mL) and extracted with 5% MeOH in $CH_2Cl_2$ (100 mL, 3x). The organic layer was washed with brine (50 mL) and dried over sodium sulfate, concentrated *in vacuo* and purified on silica column (0-10% MeOH in $CH_2Cl_2$) yielding (1-((5-bromofuran-2-yl)methyl)piperidin-3-yl)methanol (**28**) as brown oil in 50% yield (1.18 g, 4.31 mmol).

**$^1$H NMR (500 MHz, $CDCl_3$):** δ 6.23 (d, $J$ = 3.5 Hz, 1H), 6.19 (d, $J$ = 3.0 Hz, H), 3.61 – 3.58 (m , 1H), 3.50 – 3.48 (m, 3H), 2.87 (d, $J$ = 10 Hz, 1H), 2.73 – 2.71 (m, 1H), 2.15 – 1.96 (m, 3H), 1.84 – 1.57 (m, 4H), 1.09 – 1.04 (m, 1H).

**$^{13}$C NMR (126 MHz, $CDCl_3$):** δ 153.7, 120.9, 111.8, 111.7, 66.8, 56.7, 55.2, 53.6, 38.1, 27.0, 24.5

**HRMS:** $C_{11}H_{17}BrNO_2$ Calc'd $[M+H]^+$: 274.0443, Found: 274.0431.

*1.1.8 Synthesis of 2-chloro-5-(5-((3-(hydroxymethyl)piperidin-1-yl)methyl)furan-2-yl)benzoic acid (**7**)*

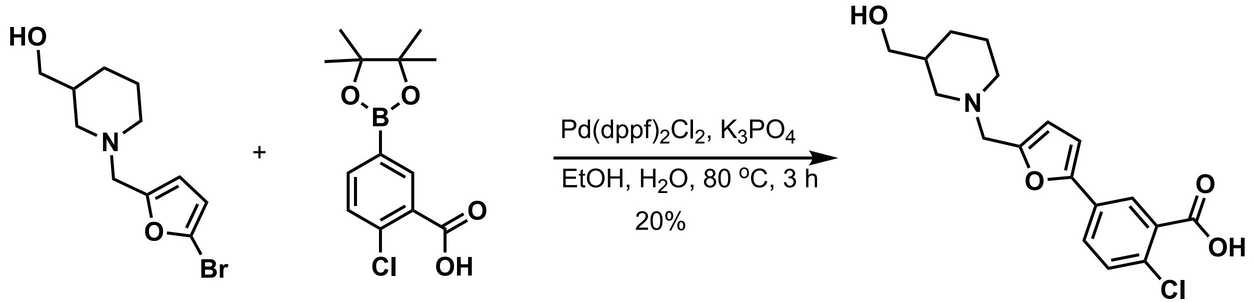


**28** **29** **7**

A solution of {1-[(5-bromo-2-furyl)methyl]-3-piperidyl}methanol (**28**, 170 mg, 0.620 mmol), 2-chloro-5-(4,4,5,5-tetramethyl-1,3,2-dioxaborolan-2-yl)benzoic acid (**29**, 175 mg, 0.620 mmol), and $K_3PO_4$ (1.0 M in $H_2O$, 1.86 mmol) in ethanol (3 mL) was degassed by evacuating and purging several times with $N_2$. Next, Pd(dppf)$Cl_2$ (25.3 mg, 0.031 mmol) was added, and the reaction was heated to reflux for 4 hours. After this time, the reaction mixture was cooled to room temperature, diluted with water (10 mL) then extracted with $CH_2Cl_2$ (20 mL x 3). The combined organics were washed with saturated brine (10 mL), dried over $Na_2SO_4$, concentrated *in vacuo*, and purified on a silica gel column (0-40% MeOH in $CH_2Cl_2$) then by reverse-phase C18 column chromatography (10-95% MeCN in water) yielding 2-chloro-5-(5-{[3-(hydroxymethyl)-1-piperidyl]methyl}-2-furyl)benzoic acid (**7**) as an off-white foamy solid in 20% yield (42.2 mg, 0.121 mmol).

**$^1$H NMR (500 MHz, CD$_3$OD):** δ 7.77 (d, *J* = 2.0 Hz, 1H), 7.58 – 7.56 (m, 1H), 7.34 (d, *J* = 8.5 Hz, 1H), 6.79 (d, *J* = 3.5 Hz, 1H), 6.62 (d, *J* = 3.5 Hz, 1H). 4.12 (s, 2H), 3.48 – 3.45 (m, 1H), 3.37 – 3.35 (m, J = 2H), 2.28 – 3.26 (m, 3H), 2.62 – 2.57 (m, 1H), 2.41 (t, *J* = 11.5 Hz, 1H), 1.94 – 1.83 (m, 2H), 1.77 – 1.69 (m, 2H), 1.17- 1.08 (m, 1H).

**$^{13}$C NMR (126 MHz, CD$_3$OD):** δ 174.7, 155.6, 147.5, 142.5, 131.2, 130.6, 130.2, 125.4, 124.7, 116.3, 108.0, 65.4, 56.7, 54.5, 54.1, 38.8, 26.6, 24.3.

**HRMS:** $C_{18}H_{21}ClNO_4$ Calc'd $[M+H]^+$: 350.1159, Found: 350.1145.

*1.1.9 Synthesis of 3-((4-hydroxyphenyl)amino)-6-methyl-1,2,4-triazin-5(4H)-one (**13**)*

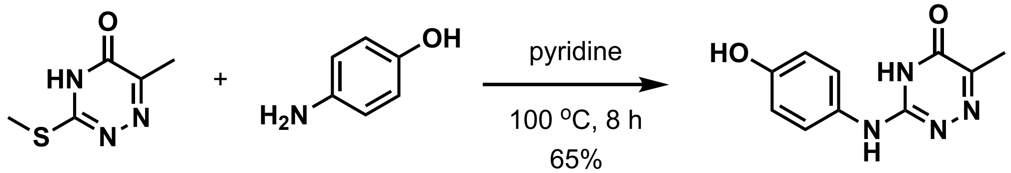


**30** **31** **13**

6-methyl-3-(methylthio)-1,2,4-triazin-5(4H)-one (**30**, 300 mg, 1.91 mmol) and 4-aminophenol (**31**, 417 mg, 1.91 mmol) were dissolved in pyridine (3 mL) and heated at 100 °C for 8 h. After this time, the reaction mixture was cooled to 0 °C and quenched with isopropyl alcohol (4 mL) which resulted in precipitation. The precipitate was filtered and washed with ice cold isopropyl alcohol 5-10 mL) afforded compound **13** in 65% yield (270 mg, 1.24 mmol).

**$^1$H NMR (500 MHz, DMSO-*d*$_6$):** δ 11.9 (bs, 1H), 9.42 (bs, 1H), 8.87 (s, 1H), 7.19 (d, *J* = 9.0 Hz, 2H), 6.74 (d, *J* = 8.5 Hz, 2H), 2.04 (s, 3H).

**$^{13}$C NMR (126 MHz, DMSO-*d*$_6$):** δ 163.4, 154.5, 154.2, 146.7, 128.4, 124.6, 115.4, 16.9.

**HRMS:** $C_{10}H_{11}N_4O_2$ Calc'd $[M+H]^+$: 219.0882, Found: 219.0871.

*1.1.10 Synthesis of 3-(2-amino-6-methylpyridin-3-yl)benzoic acid (**34**)*

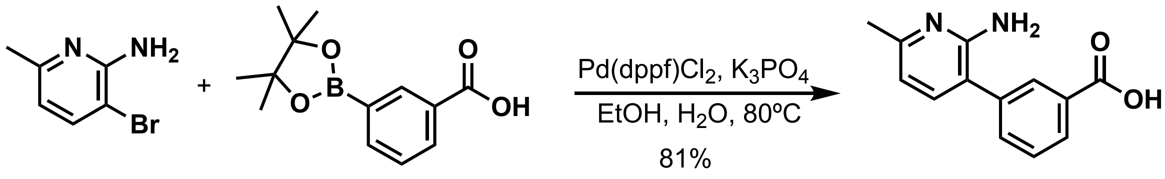


**32** **33** **34**

A solution of 3-bromo-6-methyl-2-pyridylamine (**32**, 300 mg, 1.60 mmol), m-(4,4,5,5-tetramethyl-1,3,2-dioxaborolan-2-yl)benzoic acid (**33**, 398 mg, 1.60 mmol), and $K_3PO_4$ (1.0 M in $H_2O$, 4.81 mmol) in ethanol (8 mL) was degassed by evacuating and purging several times with $N_2$. Next, $Pd(dppf)Cl_2$ (65.5 mg, 0.080 mmol) was added, and the reaction was heated to reflux for 16 hours. After this time, the reaction mixture was cooled to room temperature, diluted with water (10 mL) then extracted with $CH_2Cl_2$ (20 mL x 3). The combined organics were washed with saturated brine (10 mL), dried over $Na_2SO_4$, concentrated *in vacuo*, and purified on a silica gel column (0-20% MeOH in $CH_2Cl_2$) afforded m-(2-amino-6-methyl-3-pyridyl)benzoic acid (**34**) as an off-white solid in 81% yield (295 mg, 1.29 mmol).

**$^1$H NMR (500 MHz, DMSO-$d_6$):** δ 7.97 (s, 1H), 7.91 (d, *J* = 7.5 Hz, 1H), 7.66 (d, *J* = 7.5 Hz, 1H), 7.57 (t, *J* = 7.5 Hz, 1H), 7.27 (d, *J* = 7.5 Hz, 1H), 6.54 (d, *J* = 7.5 Hz, 1H), 5.61 (s, 2H), 2.30 (s, 3H).

**$^{13}$C NMR (126 MHz, DMSO-$d_6$):** δ 167.2, 155.6, 155.5, 138.6, 138.2, 132.8, 131.3, 129.2, 129.1, 127.9, 116.6, 112.2, 23.4.

**HRMS:** $C_{13}H_{13}N_2O_2$ Calc'd $[M+H]^+$: 229.0977, Found: 229.0966.

#### *1.1.11 Synthesis of 3-(2-amino-6-methylpyridin-3-yl)-N-(1-(1-methyl-1H-pyrazol-5-yl)propyl)benzamide (**16**)*

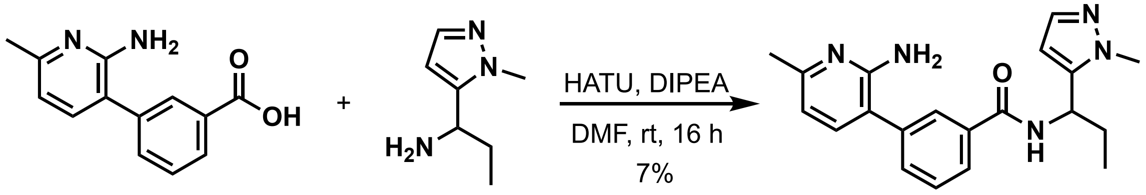


**34** **35** **16**

m-(2-Amino-6-methyl-3-pyridyl)benzoic acid (**34**, 291 mg, 1.27 mmol) and HATU (581 mg, 1.53 mmol) was dissolved in DMF (5 mL) and *N,N*-diisopropylethylamine (333 $\mu$L, 1.91 mmol) was added. The resulting mixture was stirred at room temperature for 15 min and then 1-(1-methyl-5-pyrazolyl)propylamine (**35**, 177 mg, 1.27 mmol) was added and the reaction mixture was stirred overnight. Next, the reaction mixture was diluted with water (10 mL) and extracted with ethyl acetate (50 mL x 3). The combined organic layer was washed with brine (10 mL) and dried over sodium sulfate then concentrated in *vacuo* and purified on silica column (0-20% MeOH in DCM). The product was further purified by reverse-phase C18 column chromatography (10-95% MeCN in water) afforded **16** in 7% yield as white solid (30.1 mg, 0.086 mmol).

**$^1$H NMR (500 MHz, MeOD):** δ 7.85 – 7.84 (m, 1H), 7.80 – 7.78 (m, 1H), 7.61 – 7.59 (m, 1H), 7.54 (t, *J* = 8.0 Hz, 1H), (t, *J* = 8.0 Hz, 1H),7.38 (d, *J* = 2.0 Hz, 1H), 7.33 (d, *J* = 7.5 Hz, 1H), 6.62 (d, *J* = 7.5 Hz, 1H), 6.29 (d, *J* = 2.0 Hz, 1H), 5.23 (t, *J* = 2.5 Hz, 1H), 3.89 (s, 3H), 2.36 (s, 3H), 2.00 (q, *J* = 7.5 Hz, 2H), 1.02 (t, *J* = 7.5 Hz, 3H).

**$^{13}$C NMR (126 MHz, MeOD):** δ 169.7, 157.07, 157.05, 145.6, 140.3, 139.8, 139.0, 136.2, 133.1, 130.3, 128.7, 127.6, 119.6, 114.3, 104.8, 48.0, 36.7, 28.6, 23.5, 11.2.

**HRMS:** $C_{18}H_{24}N_5O$ Calc'd $[M+H]^+$: 350.1981, Found: 350.1965.

### 1.2 Computational modeling and screening

*1.2.1 Molecular dynamics simulations identifying allosteric site and virtual screening*

Using POVME3.0 program for active-site-shape-based clustering, eight A3Bctd conformations were selected as cluster representatives of eight clusters from our prior molecular dynamics simulations of apo and nucleic acid-bound A3Bctd.[2,3] Each A3Bctd conformation was then evaluated for binding hot spots with FTMap.[4] FTprod was then used to process and compare all FTMap data.[5] Cluster0 and Cluster6 representative frames were selected for virtual screening against the active site and the putative allosteric site as they had the most-populated FTMap probes at those sites, respectively. Schrodinger's Glide was then used in standard precision mode to virtually screen ChemBridge Diversity Set of 110,000 compounds for the two sites.[6,7] A metal constraint was used for the zinc at the active site during active site virtual screen. Docking scores and ligand efficiencies were used to rank and select compounds for experimental testing.

*1.2.2 Ligand parameterization*

Parameters for **10** were generated with the Antechamber module of Amber18[8] using the Generalized Amber Force Field[9,10] with RESP HF 6-31G* charges calculated by Gaussian program.[11] The initial pose of compound **10** in A3B cluster 6 virtual screening was used as an input for induced fit docking to define the binding site and pose of the ligand. Using the induced-fit docking module of Schrodinger (version 2022u1), twenty-one different poses of A3B in complex with **10** were generated. Nine of the highest scoring poses (that had better than -319 kcal/mol) of **10** were selected and simulated for 100 ns for 3 replicates each using the Amber MD engine. After 100ns, the three most stable poses (poses 3, 6, and 8) were simulated for an additional 400 ns, for a total of 500 ns (**Figure S25**). Of these three poses, pose 6 displayed the highest stability at the allosteric site throughout all replicates and was simulated for a total of 1 μs x three replicates.

*1.2.3 Molecular dynamics simulations with **10** and analysis*

Simulations of A3Bctd and A3A with nucleotides were parameterized using the AMBER FF19SB forcefield for protein atoms, and OL15 forcefield for DNA atoms. The A3Bctd starting structure was based on the 5TD5 PDB structure, with amino acid substitutions returned to wild type amino acid sequence as described in previous work.[12,13] A3A starting structure was based on the PDB structure 5SWW, and missing loops and mutated residues were returned to wild type amino acid sequence.[13] The catalytic zinc was modeled using the Cationic Dummy Atom Model, and the zinc was modeled as being bound to an $^{-}$OH ion to represent the activated water molecule as is present in A3Bctd pre-catalysis. All systems were solvated in a TIP3P water box with a buffer of 10 Å, and ions were added to neutralize charge and obtain a concentration of 0.150 M sodium chloride solution. PROPKA was used to predict the protonation states of the sidechains.[14,15] Cysteine residues, as they were of particular interest for sidechain dynamics, were not locked into disulfide bonds. Ambertools' tLeap was used to build the final topology and coordinate files necessary for simulations.[8]

A3Bctd and A3A in complex with the **10** (and without ssDNA) were simulated for a total of 3 μs of simulation time per system. Each system underwent sequential energy minimization, followed by gradual heating and equilibration according to the following protocol. The initial energy minimization was performed on all atom excluding hydrogen with a restraint weight of 100.0 kcal/mol/$\text{Å}^2$ for 500 cycles. A second minimization of 500 cycles restrained all atoms except the protein including the ligand using a reduced restraint weight of 50.0 kcal/mol/$\text{Å}^2$. A third minimization run allowed the sidechains to minimize, restraining only backbone atoms (CA, C, N, O) with a restraint weight of 100.0 kcal/mol/$\text{Å}^2$. The fourth and final minimization run allowed an unrestrained minimization of all atoms in the system. It was conducted with 1,000,000 cycles using constant volume periodic boundary conditions using a 10Å cutoff for non-bonded interactions. In the heating step 250, positional restraints were applied to protein atoms, excluding hydrogen with a restraint weight of 2.0 kcal/mol/$\text{Å}^2$. The simulation started at 0.0 K and gradually heated to 310.0 K using a Langevin equilibration scheme (Nosé-Hoover thermostat). In the subsequent equilibration steps performed in three stages, each 250 ps, we utilized isotropic pressure control (ntb=2, ntp=1, pres0=1.0) to maintain a target pressure. In subsequent equilibration stages, the Nosé-Hoover thermostat-controlled temperature fluctuations, keeping the system at a constant temperature of 310.0 K using a relaxation parameter of 5.0. Nonbonded interactions were maintained with a cutoff of 10 Å, and positional restraints were released for increased system flexibility. A restraint was applied to the protein atoms and ligand atoms, excluding hydrogen with a restraint weight of 3.0 kcal/mol/$\text{Å}^2$. The final equilibration ran simulation ran for 125,000 timesteps with a timestep of 2 fs.

Each system (A3Bctd-**10**, A3Bctd-**10**-DNA, A3A-**10**, A3A-**10**-DNA) was simulated for 1 μs x 3 replicates of unrestrained MD simulations in an NPT ensemble at 310K with a timestep of 2 fs using the Amber22 Molecular Mechanics Engine.[8] Hydrogen bonding information and atomic distances were calculated using CPPTRAJ.[16] Contact analysis was performed using the MDTraj package Pycontact with a maximum atom-atom distance cutoff of 5.0 Å, an angle cutoff for hydrogen bonds of 120 degrees, and a distance cutoff for hydrogen bonds of 2.5 Å as recommended by PyContact.[17] The same protocol was repeated for A3Bctd and A3A in complex with the ligand in addition to ssDNA for a total of 12 μs of simulation time. Simulations were performed on a mix of computing resources that include our own GPU clusters as well as the NCSA Delta supercomputer resource.

#### 1.3 Protein expression and purification

The A3Bctd protein utilized in these studies was expressed in *Sf9* insect cells and purified by affinity chromatography by GenScript Biotech Corporation. The protein construct contains the sequence of A3B amino acid (aa) 193-382, followed by a non-cleavable $MycHis_6$ tag, based on Uniprot accession: Q9UH17-1. Upon delivery of the protein, >95% purity was established with a 4-12% polyacrylamide gel stained with Coomassie blue, and the protein kinetic activity was confirmed. A3A was expressed in HEK293T mammalian cells, according to a previous published protocol.[13] A3A was based on Uniprot accession: P31941-1 encompassing the full sequence (amino acids 1-199).

Full-length *Pyrococcus furiosus* EndoQ was expressed in *Escherichia coli* strain BL21(DE3) with a C-terminal non-cleavable His-tag (LEHHHHHH), using the pET-24a expression vector and codon-optimized coding sequence.[21] The protein was expressed in LB medium supplemented with 50 μM $ZnCl_2$ by induction with 0.5 mM isopropyl β-D-1-thiogalactopyranoside for around 16 hours at 18 °C. Protein was purified from the soluble fraction of *E. coli* lysate using nickel-affinity and Superdex 75 size-exclusion chromatography. The purified protein in 20 mM Tris (pH 7.4), 0.2 M NaCl, 5 mM β-mercaptoethanol was concentrated by ultrafiltration, flash-frozen in liquid nitrogen, and stored at −80 °C. Protein concentration was determined based on UV absorption and theoretical extinction coefficient calculated from the amino acid sequence.

#### 1.4 Deaminase assay

Assays were performed as previously described with a few modifications.[18] Compounds were suspended in DMSO at 20 mM stock concentrations. Compounds were diluted to 1.5 mM in A3 assay buffer (50 mM Tris, pH 7.4, 100 mM NaCl, 1 mM EDTA, 10% glycerol, 0.5% Triton X-100), and 10 μL was added to each respective well of a Nunc 384-well black flat-bottom plate. Recombinant A3Bctd was diluted to 10 ng/μL in A3 assay buffer, and A3A was diluted to 1 ng/μL, and 10 μL of diluted protein was added to plate wells. Compound and protein were incubated at 37°C. After 30 minutes, 10 μL of oligo substrate (5′-FAM-AAATAT**C**CCAAAGAGAGA-TAMRA-3′, purchased from IDT), diluted to 0.2 μM in 1x TE buffer, was added to all wells, then incubated for

30 minutes at 37°C. Next, 3 µL of 4 M NaOH was added to hydrolyze the ssDNA substrate, which was incubated for 30 minutes at 37°C, then neutralized by dilution with 2 M Tris, pH 7.9 (40 µL). The plates were cooled to 4°C before reading fluorescence on a Synergy H1 microplate reader (BioTek), with an excitation wavelength of 490 nm and an emission wavelength of 520 nm. The ChemBridge ID and associated compound numbering are described in **Table S1**. Activity was normalized to the DMSO control, and background was subtracted through monitoring the fluorescence of wells containing only the oligonucleotide. Data were processed in Microsoft Excel and analyzed using Prism v9.1.2 (GraphPad). $IC_{50}$ values were determined by a non-linear fit using the [Inhibitor] vs. response – Variable slope (four parameter) model. Data points and error bars are the mean and standard deviation of N = 3 technical replicates.

#### 1.5 Dose response testing of HPLC purified repurchased stocks

Compounds determined to be hits in the initial screening process were repurchased from Hit2Lead. Compounds were purified by preparative HPLC, then reanalyzed by analytical HPLC for purity. Compounds were purified using an Agilent Technologies Zorbax SB-C18 column (21.2 × 250 mm size, 7 µm pore), with a method flowing solvent A: $H_2O$, and solvent B: $CH_3CN$, with a gradient starting at 90:10 A:B from 0 to 2 minutes, followed by a linear gradient to 30:70 A:B from 2 to 22 minutes, followed by a linear gradient to 5:95 A:B from 22 to 28 minutes, and an isocratic 5:95 A:B from 28-35 minutes). Of the 21 compounds repurchased, 17 compounds were able to be purified to >95% by analytical HPLC analysis at 215 and 254 nm. Mass of the purified compounds were analyzed by mass spectrometry (LC-MS) (**Table S4**). Purified compounds were screened as described in section **1.4**, with minor experimental changes: Recombinant A3Bctd was diluted to 20 ng/µL instead of 10 ng/µL, the oligo was 5′-labeled with AlexaFluor488 instead of FAM, and the oligo was diluted to 0.3 µM instead of 0.2 µM. The full reaction (compound, protein, and oligo) was incubated for 45 minutes, instead of 30 minutes.

#### 1.6 UDG control inhibition assay

The UDG control assay was performed similar to the deaminase assay as described in **1.4**. Compounds were diluted to 3 mM in A3 assay buffer, and 10 µL of each compound was added to the plate. To prepare the UDG (5 U/mL, New England BioLabs), 1 µL of stock UDG was diluted to 2 mL in A3 assay buffer, and 10 µL of diluted UDG was added to the appropriate wells. Compounds and UDG were incubated for 30 minutes at 37°C. Meanwhile, the oligo substrate (5′-FAM-AAATAT**U**CCAAAGAGAGA-TAMRA-3′, purchased from IDT) was diluted to 0.2 µM in 1X TE buffer, then 10 µL was added to each well and the reaction was incubated for 45 minutes. The reaction was quenched with the addition of 3 µL of 4 M NaOH, incubated for 30 minutes at 37°C, then cooled to 4°C before reading on a Synergy H1 microplate reader (BioTek). Data were analyzed using Prism v9.1.2 (GraphPad), with a non-linear fit using the [Inhibitor] vs. response – Variable slope (four parameter) model. Data points and error bars are the mean and standard deviation of N = 3 technical replicates.

#### 1.7 Gel-based deaminase assay

The DNA deamination assay was conducted as previously described with minor changes.[19,20] A3Bctd (15 nM) was incubated in activity buffer (50 mM Tris-HCl, pH 8.0, 1 mM $MgCl_2$, 0.01% Tween-20) with 1 mM of compounds (or metals) for 30 minutes at 37°C. Following 30 minutes of incubation, a solution of the 3′-fluorescein(6-FAM)-labeled 21-mer DNA oligonucleotide (0.2 µM) (5′-ATTATGT**C**GTTGGATTTATTT-6-FAM-3′) in activity buffer was added to the solution for another 30 minutes at 37°C (final volume 10 µL). After incubation (37 °C, 1 hour) pfuEndoQ (produced in *E. coli* as previously reported[19]) was added to the mixture and incubated (1.0 µM in activity buffer, 60 °C, 10 minutes, final volume 11 µL). The reactions were stopped by the addition of 80% formamide and heating (95 °C, 10 minutes). 5 µL of each sample were separated on a 15% TBE-Urea gel (300 V, 30 minutes, 1x TBE buffer) which was scanned using fluorescent detection using a Typhoon FLA 7000 biomolecular imager (GE Healthcare Life Sciences) to visualize the DNA. To quantify the percent of deamination, ImageJ was used to perform densitometry. The control substrate (5′-ATTATGT**dU**GTTGGATTTATTT-6-FAM-3′) was used to control for EndoQ activity. DNA was purchased from IDT.

#### 1.8 Fluorescence polarization assay

Competitive fluorescence polarization assay was adapted from previous direct binding fluorescence polarization assays. A3Bctd was diluted to 2.7 µM and tracer (5′-AlexaFluor546-ATTATGTCGTTGGATTTATTT-3′) was diluted to 15 nM in a 50:50 mix of protein buffer (25 mM Tris HCl pH 6.0, 75 mM NaCl, 0.5 mM EDTA) and DNA buffer (10 mM Tris HCl pH 6.0, 0.5 mM EDTA). In a black low volume Corning 384-well plate, 20 µL of this premixed solution was added to appropriate wells. Compounds were dispensed with an HP D300 digital dispenser with the DMSO normalized to 3%. Plates were shaken on an orbital rotating block at 1500 rpm for 1 minute to mix, then centrifuged at 1000 rpm for 1 minute. The reaction was incubated for 20 minutes at room temperature, then read on a Tecan Spark, with a monochromator-based excitation wavelength of 530 $\pm$ 5 nm and a filter-based emission wavelength of 620(10) nm. Anisotropy values were background corrected based on the average of N ≥ 8 technical replicates of wells containing tracer only. Data were analyzed using Prism v9.1.2 (GraphPad), with a non-linear fit using the specific binding with Hill slope model. Data points and error bars are the mean and standard deviation of N = 3 technical replicates.

#### 1.9 Intact protein mass spectrometry

Protein samples were desalted into assay buffer (10mM Tris, pH 8.0), and concentration was determined via nanodrop. Protein concentration was adjusted to 26.3 µM using assay buffer, and 47.5 µL of this solution was transferred to a 0.6 mL microcentrifuge tubes. Stock solutions of compounds **25** (chloroacetamide) and **26** (acrylamide) were diluted in DMSO at the following concentrations (10 mM, 5 mM, 2.5 mM, 0.5 mM). 2.5 µL of the DMSO stocks were added to the 47.5 µL solution of protein (final volume = 50 µL, final protein concentration = 25 µM). The final concentrations of the compounds were 500 µM (10 mM stock), 250 µM (5 mM stock), 125 µM (2.5 mM stock), and 25 µM (0.5 mM stock). These solutions were then left to incubate for one or three hours at 37 °C. After incubation, these solutions were then desalted using 0.5 mL 7 kDa Zeba desalting spin columns (ThermoScientific). Protein samples were analyzed with an UltiMate™ 3000 liquid chromatography system (ThermoFisher) in-line with an Orbitrap Elite (ThermoFisher) hybrid mass spectrometer. The LC eluent system was water with 0.5% formic acid (Eluent A) and acetonitrile with 0.5% formic acid (Eluent B). The LC column was a Zorbax 300 SB-C3 Rapid Resolution HD column, 2.1x100 mm, 1.8 µm particle size (Agilent). The LC gradient (400 µL/min flow rate, 10 µL injection) total time was 4 minutes, with the following steps: 0 min, 20% B; 0.75 min, 100% B; 1.5 min, 20% B; 3 min, 100% B; 3.5 min, 100% B; 4.0 min, 20% B. The first ramp up and down was used to desalt the sample, and the column flow was sent to waste for the first 1.5 min; the second ramp up resulted in elution of the protein. The Orbitrap settings were as follows: FTMS analyzer in positive mode, 30,000 resolution, m/z scan range of 600-2000. Resulting spectra were deconvoluted with the Protein Deconvolution software (ThermoFisher) with the following settings: m/z range = 800-2000, S/N threshold = 3, relative abundance threshold = 5. Percent adduction was calculated by dividing the intensity of the adduct species by the sum of the intensities, multiplied by 100. '% Adducted' = [(intensity of mass adduct species)/(intensity of unadducted species + intensity of mass adduct species)] x 100. N = 2 biological replicates.

#### 1.10 Bottom-up protein digestion with 25

Incubation and desalting of A3A and A3Bctd was conducted as described in **1.8**, where A3A was incubated with 250 µM **24** and A3Bctd with 500 µM **24** and then desalted with 0.5 mL 7kDa Zeba desalting spin columns. The solutions were then dried via SpeedVac overnight. Roughly 5 µg of each protein sample were separately reconstituted in 20 µL of denaturing buffer (8 M Urea, 50 mM DTT, 100 mM $NH_4HCO_3$) and incubated for one hour at 37 °C to denature the proteins. 20 µL of 50 mM iodoacetamide (IAA) was added to each of the samples and incubated in the dark at room temperature for 30 minutes, to alkylate exposed cysteines. 160 µL of dilution buffer (50 mM $NH_4HCO_3$, 1 mM $CaCl_2$) and 20 µL of acetonitrile was added to each sample to reduce the concentration of Urea. Glu-C endoproteinase (ThermoScientific) was reconstituted in MilliQ water at a concentration of 0.01 µg/µL, and 10 µL were added to each sample for a final amount of 0.1 µg (1:50, enzyme:protein). This solution was then left to incubate overnight at 37 °C. The next day 10 µL of 0.01 µg/µL of Trypsin Gold (Promega), for a final amount of 0.1 µg (1:50, enzyme:protein) was added to samples and left to incubate for four hours at 37 °C. 20 µL of glacial acetic was then added to quench the reaction, and then the samples were concentrated to 80 µL using a SpeedVac. The solutions were then adjusted to 0.5% TFA by adding 20 µL of a 25% TFA in water solution, prior to desalting by manufacture protocol using 100 µL Pierce C18 Tips (ThermoScientific). Samples were then evaporated to dryness overnight using SpeedVac. Samples were reconstituted in 2% acetonitrile, 98% water containing 0.1% formic acid, at 0.2 µg/µL. 3µL of sample was loaded onto a home-packed analytical C18 reverse phase column with a 10 µm emission tip (75 µm × 200 mm [New Objective, Woburn, MA], with Luna C18 5 µm particles [Phenomenex, Torrance, CA]) via UltiMate™ 3000 liquid chromatography system (ThermoScientific). Peptides were eluted with buffer A (0.1% formic acid in water) and buffer B (0.1% formic acid in acetonitrile) with the following gradient profile: 0-5.5 min for loading, 2% B, flow rate 1 µL/min; 6-12 min, 2%-10% B, 0.3 µL/min; 12-52 min, 10%-25% B, 0.3 µL/min; 52-57 min, 25%-40% B, 0.3 µL/min; 57-58 min, 40%-85% B, 0.3 µL/min; 58-62 min 85% B, 0.3 µL/min; 62-63 min, 85%-2% B, 0.3 µL/min; 63-65 min, 2% B, 1.0 µL/min. Mass spectrometry was obtained on an Orbitrap Fusion Tribrid Mass Spectrometer (ThermoFisher) in positive nanospray ionization mode. The mass spectrometer conditions were: spray voltage 2.2 kV, ion transfer tube temperature 300 °C. MS survey scans were performed with a cycle time of 3 s. After each survey scan, the 10 to 20 most abundant precursor ions with $z > 1$ were selected for fragmentation using higher energy collisional dissociation (HCD). $MS^1$ resolution was at 120,000 with a scan range of 375-1500. $MS^2$ resolution was fixed at 15,000. N = 3 biological replicates of the digestion and mass spectrometry analysis were performed. The resulting data was analyzed using Proteome Discoverer 3.0 (ThermoScientific) for chromatogram processing and fragment spectra isolation.

Peptides were analyzed by referencing the A3Bctd and A3A protein sequences from a FASTA file created in-house, with cysteine carbamidomethylation and the compound **24** molecular weight set as dynamic modifications.

#### 1.11 Inductively Coupled Plasma-Mass Spectrometry

ICP-MS was performed by Triclinic Labs (Lafayette, IN) for the detection of Zn, Au, Cu, Pd, Pt, and Co. The submitted solutions were diluted 1000x with aqueous 2% nitric acid / 0.5% hydrochloric acid solution, and the diluent was also used for background subtraction. The samples were subsequently analyzed by ICP-MS. The samples were analyzed using a Thermo iCAP RQ ICP-MS. The analysis was performed using a helium collision gas to reduce polyatomic interferences by kinetic energy discrimination (KED). A 5-point calibration curve (0.01 µg/L – 100 µg/L) for the elements of interest was established using multielement standard solutions. The results above are reported by Qtegra software, which calculates concentrations by subtracting unknown values from the blank and uses the calibration curve to calculate the analyte content. Values were corrected for the prepared solution's concentration and reported as ppb (or µg of analyte per L) of the as-received sample. Instrument operational conditions are outlined in Table below.

| | |
|---|---|
| **Plasma Power (W)** | 1550 |
| **Plasma Gas Flow (L/min)** | 14 |
| **Nebulizer Gas Flow (L/min)** | 0.99 |
| **Auxiliary Gas Flow (L/min)** | 0.8 |
| **CCT Flow (L/min)** | 4.6 |
| **Sample Uptake Delay (sec)** | 60 |
| **Rinse Time (sec)** | 90 |
| **Read time (sec)** | 2.0 |
| **Replicates** | 3 |

#### 1.12 Zinc Detection via Colorimetric Assay Kit (Abcam)

The relative amount of zinc present in A3B upon different treatment was measured using the commercial colorimetric zinc detection assay supplied by Abcam (ab102507) and performed following the manufacturer’s instructions, with minor changes. Prior to incubation with the various concentrations of palladium (II) acetate, A3Bctd was desalted into buffer (20 mM Tris, pH 8.0, 50 mM NaCl) using 0.5 mL 7kDa Zeba desalting spin columns. A3Bctd was then diluted to 63 µM in buffer and 22.8 µL of A3Bctd was added to 1.2 µL of DMSO or palladium (II) acetate (final volume = 24 µL, final A3Bctd concentration = 60 µM, final palladium concentration = 60 µM, 600 µM, and 3 mM). After incubation for 30 minutes at 37 °C, the samples were desalted using the mini (75 µL) 7 kDa Zeba desalting spin columns to remove excess metal or dislodged zinc. After desalting, an equal volume of 7% TCA was added to the desalted protein samples and allowed to incubate at room temperature for 10 minutes. Following incubation, the samples were centrifuged at 15,000 rcf for 5 minutes. The supernatant was then transferred to a clean tube and vortexed prior to plating 12.5 µL in the 384-well plate (Thermo Scientific, 242757). The zinc standards (0, 1, 2, 3, 4, 5 nmol/well zinc) were prepared following the manufacturers protocol, and 12.5 µL were added to the plate. One part 'Zinc Reagent 2' was added to four parts of 'Zinc Reagent one' and 50 µL of this solution was added to every well. The plate was allowed to incubate for 10 minutes at room temperature before the absorbance at 560 nm (bandwidth 3.5 nm) was read on a Tecan Spark plate reader. The zinc concentrations were then calculated using the equation obtained from the standard curve. Each experiment was conducted in technical duplicate, and the bar plots in the main text represent the average of N = 3 biological duplicates with error represented as standard deviation (SD).

### 2.0 Supplementary data

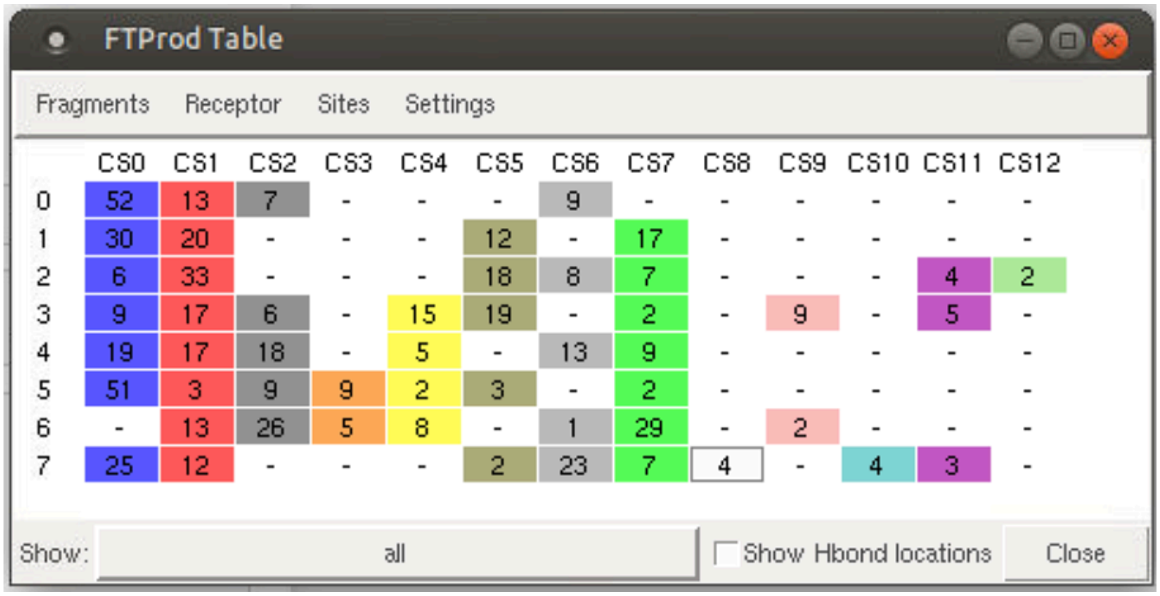


| | CS0 | CS1 | CS2 | CS3 | CS4 | CS5 | CS6 | CS7 | CS8 | CS9 | CS10 | CS11 | CS12 |
|---|---|---|---|---|---|---|---|---|---|---|---|---|---|
| 0 | 52 | 13 | 7 | - | - | - | 9 | - | - | - | - | - | - |
| 1 | 30 | 20 | - | - | - | 12 | - | 17 | - | - | - | - | - |
| 2 | 6 | 33 | - | - | - | 18 | 8 | 7 | - | - | - | 4 | 2 |
| 3 | 9 | 17 | 6 | - | 15 | 19 | - | 2 | - | 9 | - | 5 | - |
| 4 | 19 | 17 | 18 | - | 5 | - | 13 | 9 | - | - | - | - | - |
| 5 | 51 | 3 | 9 | 9 | 2 | 3 | - | 2 | - | - | - | - | - |
| 6 | - | 13 | 26 | 5 | 8 | - | 1 | 29 | - | 2 | - | - | - |
| 7 | 25 | 12 | - | - | - | 2 | 23 | 7 | 4 | - | 4 | 3 | - |

**Figure S1**. FTProd results showing the distribution of binding hot spots for eight cluster representative A3Bctd structures (from 0 to 7). Cluster0 and Cluster6 have the largest and second largest FTMap probe populations for the active site (CS0) and the putative allosteric site (CS7), respectively. Number of FTMap probes for each site is tabulated in the below table. CS stands for consensus site.

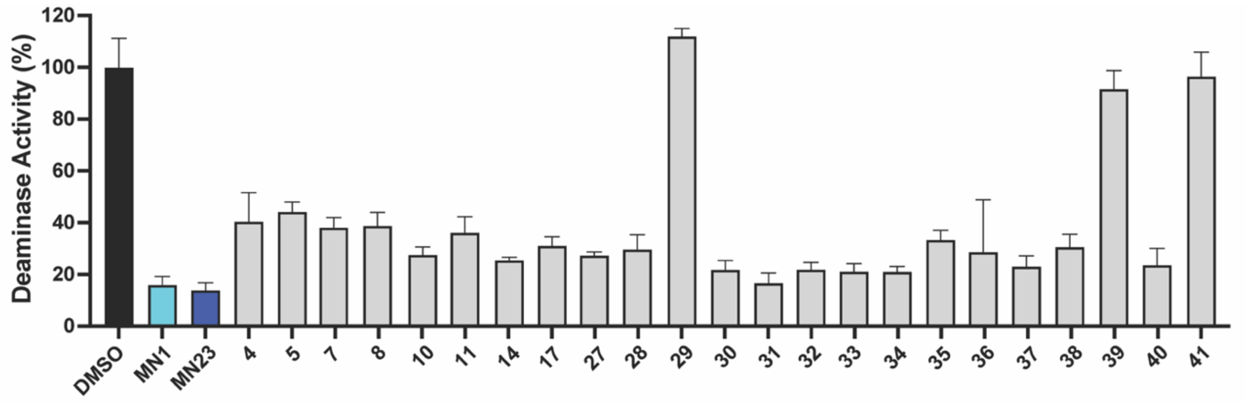


**Figure S2**. Single point re-testing of compounds selected as hits from the initial screening. Compounds tested at 500 μM concentration in the fluorescence-based activity assay against A3Bctd. Bars are the mean of N ≥ 3 technical replicates, errors bars are the standard deviation.

| Compound | ChemBridge ID | Replicate 1 | Replicate 2 |
|---|---|---|---|
| 4 | 33616296 | 16.4 | 24.2 |
| 5 | 69830916 | 7.4 | 15.0 |
| 7 | 54214567 | 23.8 | N/A[⊥] |
| 8 | 21647897 | 17.7 | 12.2 |
| 10 | 22128731 | -8.1 | 1.1 |
| 11 | 9123731 | 23.7 | 22.8 |
| 14 | 81965134 | 19.8 | 7.1 |
| 17 | 79828650 | 18.1 | 27.5 |
| 27 | 64062448 | 19.8 | 21.9 |
| 28 | 7741584 | 12.4 | 7.3 |
| 29 | 33650212 | 14.6 | 24.7 |
| 30 | 27048541 | -4.6 | 0.3 |
| 31 | 93915334 | -3.6 | -3.2 |
| 32 | 15818196 | -4.3 | 2.4 |
| 33 | 82460954 | -5.9 | -1.2 |
| 34 | 44060191 | -4.7 | -3.1 |
| 35 | 49253890 | 18.5 | 10.3 |
| 36 | 5541604 | 3.5 | 14.1 |
| 37 | 7954304 | 5.4 | 5.8 |
| 38 | 7740811 | -0.4 | 1.6 |
| 39 | 7961510 | 3.0 | 2.8 |
| 40 | 9152831 | 16.4 | 29.5 |
| 41 | 9287220 | 6.6 | 7.4 |

**Table S1**. Compounds selected as hits from primary screening and their individual biological replicate activity values (% residual deaminase activity). [⊥]In the second replicate of the primary screen, the column containing this compound had a missed liquid addition – therefore all the values for the compounds in that row were excluded from analysis. Out of an abundance of caution, the compound was selected for further confirmation testing.

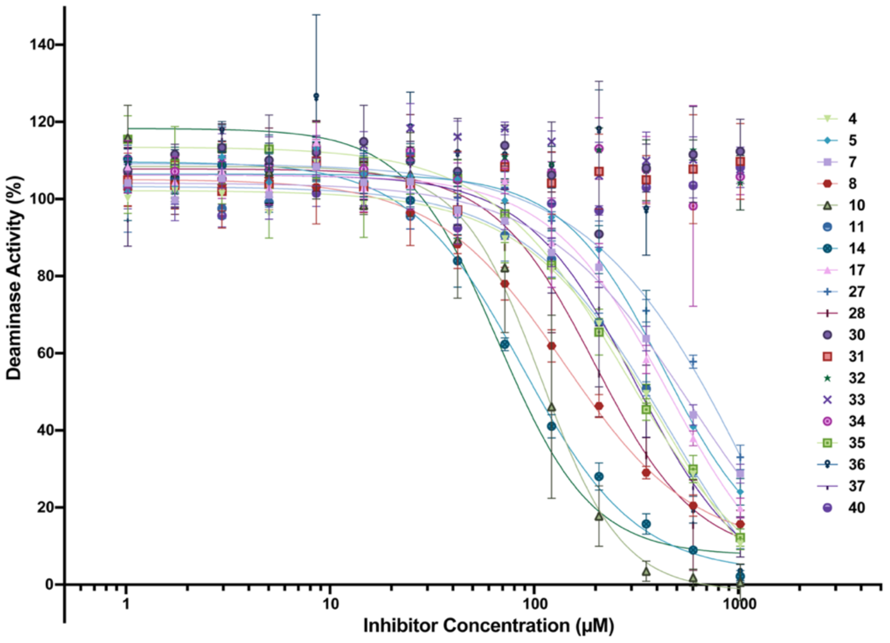


**Figure S3**. Dose response deaminase assay testing of compounds against A3Bctd that confirmed in the single point response in Figure S2.

| Compound | UDG Activity (%) | Compound | UDG Activity (%) |
|---|---|---|---|
| 5 | 108 ± 4 | 12 | 108 ± 3 |
| 6 | 101 ± 5 | 13 | 108 ± 4 |
| 7 | N/A** | 14 | 110 ± 5 |
| 8 | 99.5 ± 4.8 | 15 | 97.8 ± 3.2 |
| 9 | 92.4 ± 3.3 | 16 | 103 ± 3 |
| 10 | 97.8 ± 3.8 | 17 | 105 ± 3 |
| 11 | 96.0 ± 3.5 | | |

**Table S2.** Summary of results of control testing for primary hit inhibition of UDG in coupled activity assay. UDG activity is the average ± SEM of two biological replicates. **There was not enough material to test compound **7**, but when the compound was tested in the activity assay against A3A, it did not inhibit, indicating the compound is likely not an inhibitor of UDG.

| Compound # | 215 nm purity (%) | 254 nm purity (%) | Compound # | 215 nm purity (%) | 254 nm purity (%) |
|---|---|---|---|---|---|
| 1 | 95.2 | 97.2 | 11 | 99.7 | 99.2 |
| 2 | 99.5 | 99.9 | 12 | 98.6 | 96.8 |
| 3 | 99.6 | 99.8 | 13 | 98.1 | 98.3 |
| 4 | 97.8 | 99.1 | 14 | 99.6 | 99.9 |
| 5 | 95.7 | 95.1 | 15 | 97.4 | 98.4 |
| 6 | 99.6 | 99.9 | 16 | 97.9 | 98.5 |
| 7 | 95.7 | 96.0 | 17 | 95.8 | 95.5 |
| 8 | 98.7 | 98.7 | 27 | N/A** | N/A** |
| 9 | 95.8 | 98.6 | 28 | N/A§ | N/A§ |
| 10 | 95.1 | 98.2 | | | |

**Table S3**. Purities of repurchased compounds analyzed by analytical HPLC after preparative HPLC purification. ****27** hydrolyzed quickly in water. After every purification, as soon as the sample was dissolved in water to analyze by HPLC, the sample was already degraded to <95% purity, see HPLC trace below. §**28** precipitates in water when diluted to be analyzed by HPLC.

| Compound | Calc $[M+H]^+$ | Found $[M+H]^+$ | Compound | Calc $[M+H]^+$ | Found $[M+H]^+$ |
|---|---|---|---|---|---|
| 1 | 346.2 | 346.4 | 10 | 371.2 | 371.3 |
| 2 | 287.2 | 287.2 | 11 | 205.2 | 205.2 |
| 3 | 352.2 | 352.3 | 12 | 378.4 | 378.3 |
| 4 | 379.2 | 379.3 | 13 | 219.2 | 219.2 |
| 5 | 338.4 | 338.3 | 14 | 389.2 | 389.3 |
| 6 | 352.5 | 352.3 | 15 | 298.2 | 298.3 |
| 7 | 350.8 | 350.2 | 16 | 350.4 | 350.3 |
| 8 | 352.2 | 352.3 | 17 | 316.4 | 316.3 |
| 9 | 259.2 | 259.2 | | | |

**Table S4.** LC-MS analysis of purified compound stocks tested in dose response.

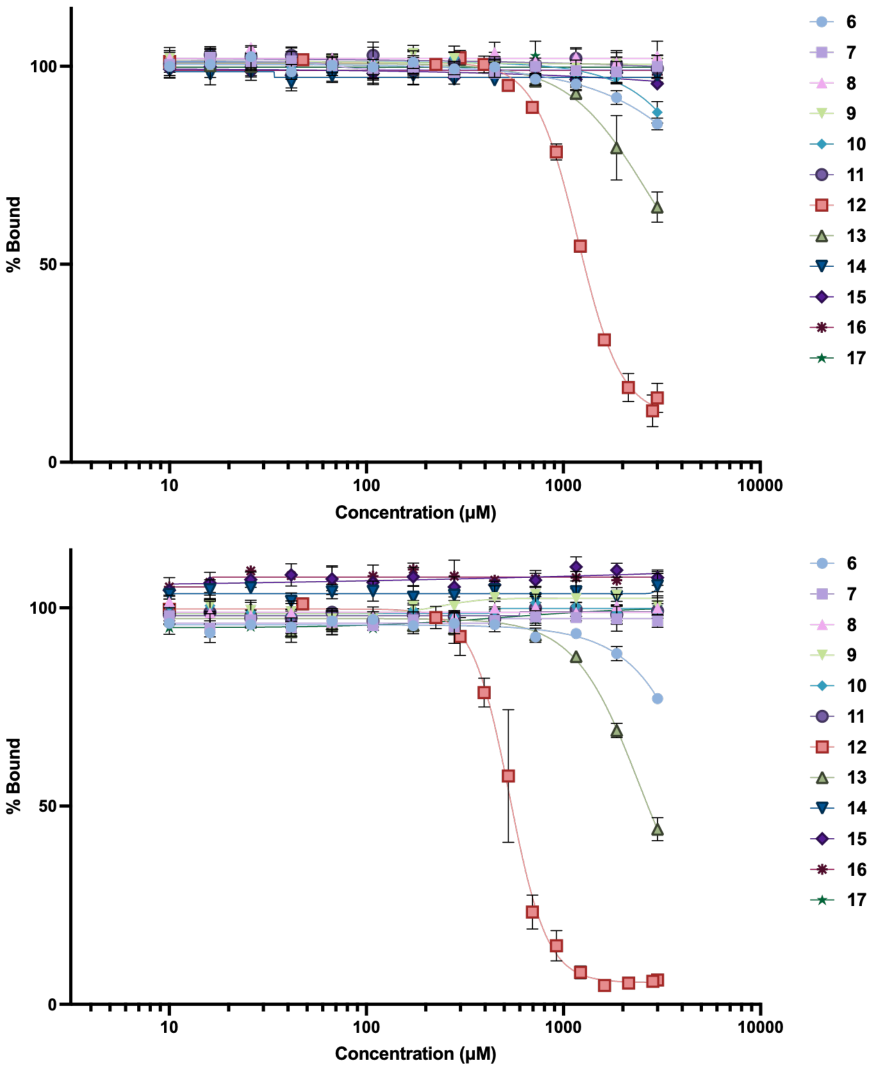


**Figure S4**. Both biological replicates of the fluorescence polarization assay against A3Bctd to test the ability of the compounds to displace DNA. Each point is mean ± standard error of the mean of N = 3 technical replicates. The average $IC_{50}$ of **12** is measured to be 820 µM.

| Compound number | ChemBridge ID number | Structure | Activity $IC_{50}$ (μM) | FP $IC_{50}$ (μM) | Predicted Binding Site |
|---|---|---|---|---|---|
| **1** | 26574649 | 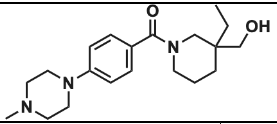 | >5000 | N/A | Allosteric |
| **2** | 31266820 | 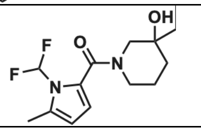 | >5000 | N/A | Allosteric |
| **3** | 38879824 | 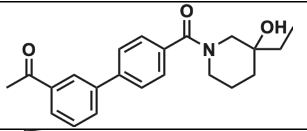 | >5000 | N/A | Allosteric |
| **4** | 33616296 | 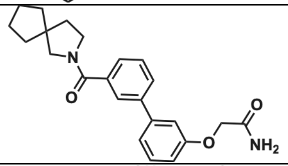 | >5000 | N/A | Allosteric |
| **5** | 69830916 | 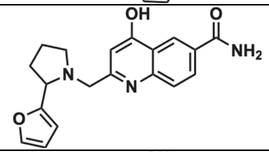 | $318 \pm 18$ | >3000 | Allosteric |
| **6** | 37925944 | 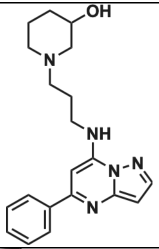 | $127 \pm 11$ | >3000 | Active site |
| **7** | 54214567 | 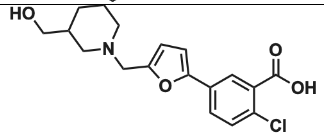 | $57.9 \pm 4.9$ | >3000 | Allosteric |
| **8** | 21647897 | 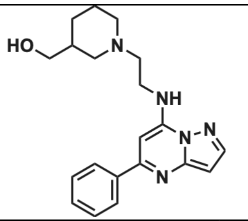 | $103 \pm 8$ | >3000 | Active site |
| **9** | 33116028 | 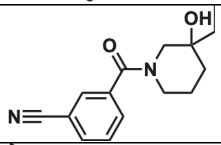 | $548 \pm 47$ | >3000 | Allosteric |
| **10** | 22128731 | 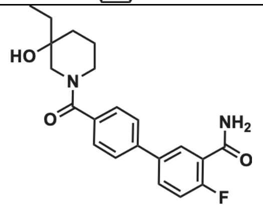 | $772 \pm 57$ | >3000 | Allosteric |

| | | | | | |
|---|---|---|---|---|---|
| **11** | 9123731 | 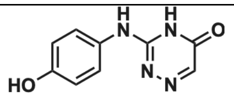 | 849 ± 52 | >3000 | Allosteric |
| **12** | 81493496 | 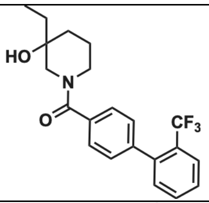 | 520 ± 26 | 861 ± 22 | Allosteric |
| **13** | 7985324 | 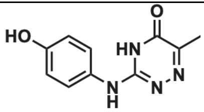 | 123 ± 9 | >3000 | Allosteric |
| **14** | 81965134 | 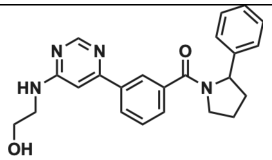 | 606 ± 132 | >3000 | Active site |
| **15** | 96694062 | 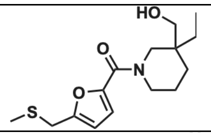 | 501 ± 33 | >3000 | Allosteric |
| **16** | 77959601 | 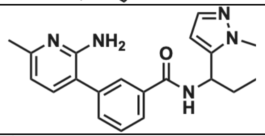 | 744 ± 54 | >3000 | Allosteric |
| **17** | 79828650 | 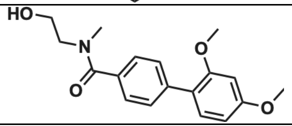 | 659 ±52 | >3000 | Active site |

**Table S5.** Structures, paper numbering, ChemBridge ID and associated $IC_{50}$'s of the 17 repurified compounds, along with the putative binding site based on virtual screening. $IC_{50}$ is the average ± SEM of two biological replicates.

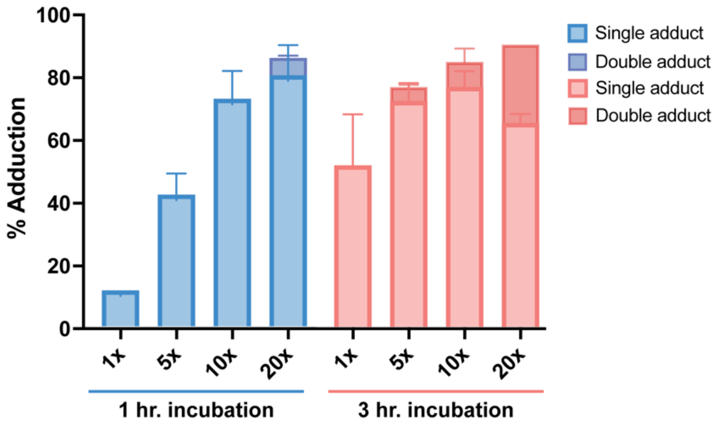


**Figure S5**. Percent adduction of **24** to A3Bctd using intact protein mass spectrometry, N = 2, error bars represent standard deviation.

| MW of species (Da) | | Incubation = 1 hr 25 μM (1X) | 125 μM (5X) | 250 μM (10X) | 500 μM (20X) |
|---|---|---|---|---|---|
| **25338 no adduct** | Replicate 1 | 88% | 52% | 20% | 7% |
| | Replicate 2 | 89% | 64% | 35% | 22% |
| | Average | 88.5% | 58% | 27.5% | 14.5% |
| **25721 (+383)** | Replicate 1 | 12% | 48% | 80% | 88% |
| | Replicate 2 | 11% | 36% | 65% | 72% |
| | Average | 11.5% | 42% | 72.5% | 80% |
| **26104 (+766)** | Replicate 1 | 0 | 0 | 0 | 5% |
| | Replicate 2 | 0 | 0 | 0 | 6% |
| | Average | 0 | 0 | 0 | 5.5% |

| MW of species (Da) | | Incubation = 3 hr 25 μM (1X) | 125 μM (5X) | 250 μM (10X) | 500 μM (20X) |
|---|---|---|---|---|---|
| **25338 no adduct** | Replicate 1 | 37% | 19% | 9% | 0 |
| | Replicate 2 | 60% | 28% | 22% | 0 |
| | Average | 48.5% | 23.5% | 15.5% | 0 |
| **25721 (+383)** | Replicate 1 | 63% | 76% | 80% | 67% |
| | Replicate 2 | 40% | 68% | 73% | 63% |
| | Average | 51.5% | 72% | 76.5% | 65% |
| **26104 (+766)** | Replicate 1 | 0 | 5% | 11% | 25% |
| | Replicate 2 | 0 | 4% | 5% | 25% |
| | Average | 0 | 4.5% | 8% | 25% |
| **26487 (+1149)** | Replicate 1 | 0 | 0 | 0 | 8% |
| | Replicate 2 | 0 | 0 | 0 | 12% |
| | Average | 0 | 0 | 0 | 10% |

**Table S6.** Average percent adduction of A3Bctd treated with **24** at variable concentration and time, from intact protein mass spectrometry deconvolution.

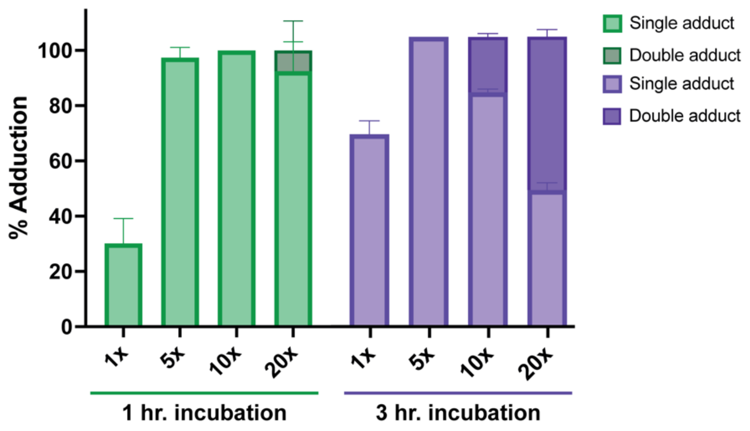


**Figure S6**. Percent adduction of **24** to A3A using intact protein mass spectrometry, N = 2, error bars represent standard deviation

| MW of species (Da) | | Incubation = 1 hr | | | |
|---|---|---|---|---|---|
| | | 25 µM (1X) | 125 µM (5X) | 250 µM (10X) | 500 µM (20X) |
| **25927 no adduct** | Replicate 1 | 76% | 0 | 0 | 0 |
| | Replicate 2 | 64% | 5% | 0 | 0 |
| | Average | 70% | 2.5% | 0 | 0 |
| **26309 (+382)** | Replicate 1 | 24% | 100% | 100% | 85% |
| | Replicate 2 | 36% | 95% | 100% | 100% |
| | Average | 30% | 97.5% | 100% | 92% |
| **26691 (+764)** | Replicate 1 | 0 | 0 | 0 | 15% |
| | Replicate 2 | 0 | 0 | 0 | 0 |
| | Average | 0 | 0 | 0 | 7.5% |

| MW of species (Da) | | Incubation = 3 hr | | | |
|---|---|---|---|---|---|
| | | 25 µM (1X) | 125 µM (5X) | 250 µM (10X) | 500 µM (20X) |
| **25927 no adduct** | Replicate 1 | 30% | 0 | 0 | 0 |
| | Replicate 2 | 37% | 0 | 0 | 0 |
| | Average | 33.5% | 0 | 0 | 0 |
| **26309 (+382)** | Replicate 1 | 70% | 100% | 80% | 49% |
| | Replicate 2 | 63% | 100% | 82% | 45% |
| | Average | 66.5% | 100% | 81% | 47% |
| **26691 (+764)** | Replicate 1 | 0 | 0 | 20% | 51% |
| | Replicate 2 | 0 | 0 | 18% | 55% |
| | Average | 0 | 0 | 19% | 53% |

**Table S7.** Average percent adduction of A3A treated with **24** at variable concentration and time, from intact protein mass spectrometry deconvolution.

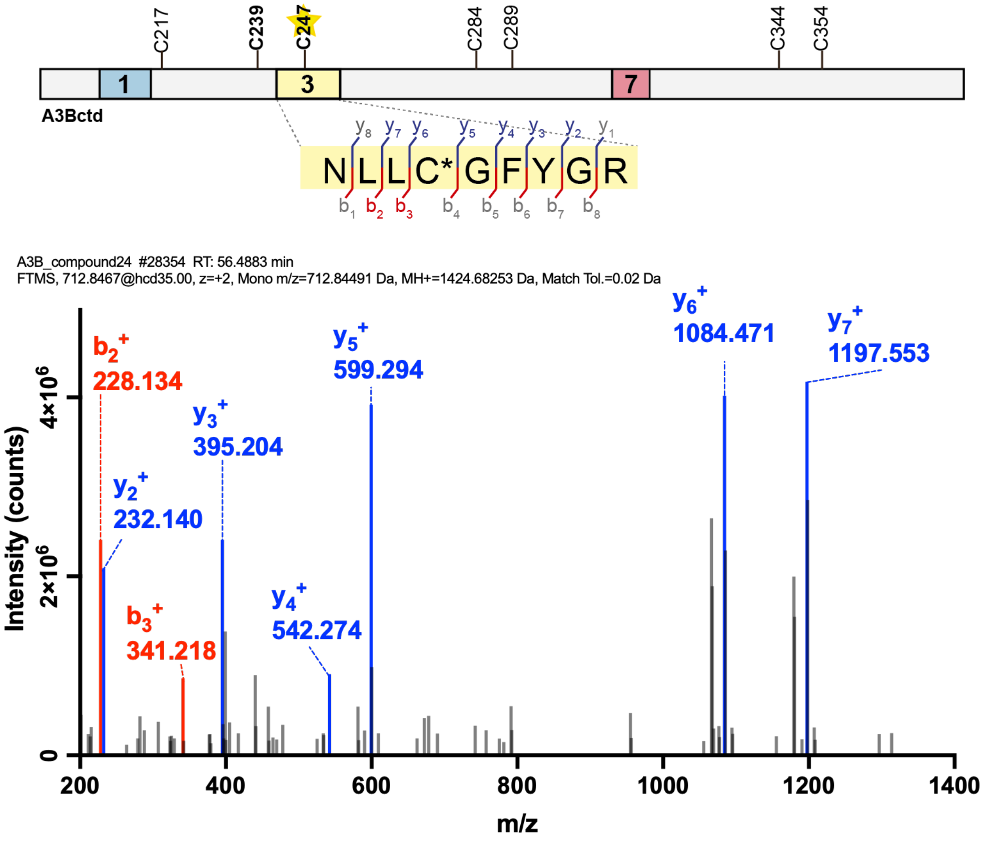


**Figure S7.** Collision-induced fragmentation pattern of major adducted peptide from A3Bctd (25 μM) treated with 500 μM (20x) **24**, incubated for one hour. 12 peptide-spectrum matches (PSM) identified.

| $B^+$ | calculated | detected | sequence | $y^+$ | calculated | observed |
|---|---|---|---|---|---|---|
| 1 | 115.05020 | nd | N | 9 | - | - |
| 2 | 228.13427 | 228.13443 | L | 8 | 1310.64133 | nd |
| 3 | 341.21833 | 341.21680 | L | 7 | 1197.55727 | 1197.55847 |
| 4 | 826.398 | nd | C+adduct | 6 | 1084.47321 | 1084.47046 |
| 5 | 883.419 | nd | G | 5 | 599.29362 | 599.29279 |
| 6 | 1030.488 | nd | F | 4 | 542.27216 | 542.27228 |
| 7 | 1193.551 | nd | Y | 3 | 395.20374 | 395.20349 |
| 8 | 1250.573 | nd | G | 2 | 232.14042 | 232.14072 |
| 9 | - | - | R | 1 | 175.11895 | nd |

**Table S8.** Table of calculated and identified ions from **Figure S7.**

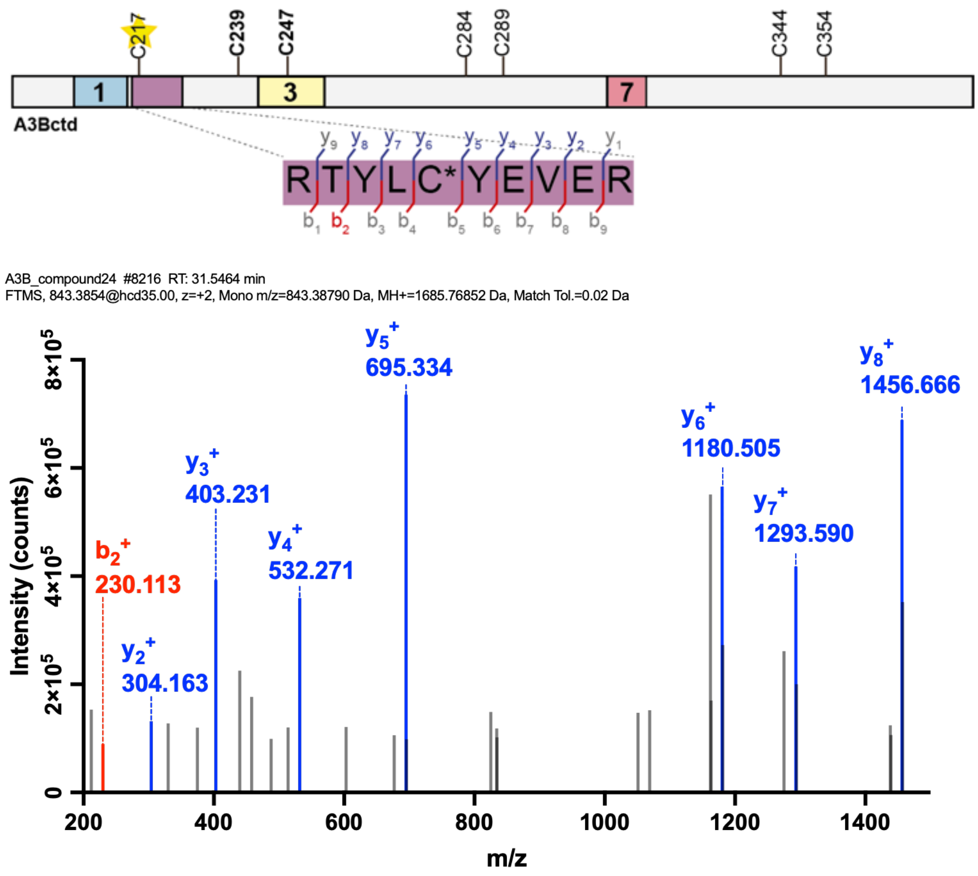


**Figure S8.** Collision-induced fragmentation pattern of minor adducted peptide from A3Bctd (25 µM) treated with 500 µM (20x) **24**, incubated for one hour. 6 peptide-spectrum matches (PSM) identified.

| $B^+$ | calculated | detected | sequence | $y^+$ | calculated | observed |
|---|---|---|---|---|---|---|
| **1** | 129.06585 | nd | **Q** | **10** | - | - |
| **2** | 230.11353 | 230.11304 | **T** | **9** | 1557.71054 | nd |
| **3** | 393.17686 | nd | **Y** | **8** | 1456.66286 | 1456.66565 |
| **4** | 506.26092 | nd | **L** | **7** | 1293.59953 | 1293.59033 |
| **5** | 991.44051 | nd | **C+adduct** | **6** | 1180.51547 | 1180.50537 |
| **6** | 1154.50384 | nd | **Y** | **5** | 695.33588 | 695.33826 |
| **7** | 1283.54643 | nd | **E** | **4** | 532.27255 | 532.27112 |
| **8** | 1382.61484 | nd | **V** | **3** | 403.22996 | 403.23105 |
| **9** | 1511.65744 | nd | **E** | **2** | 304.16155 | 304.16333 |
| **10** | - | - | **R** | **1** | 175.11895 | nd |

**Table S9.** Table of calculated and identified ions from **Figure S8.**

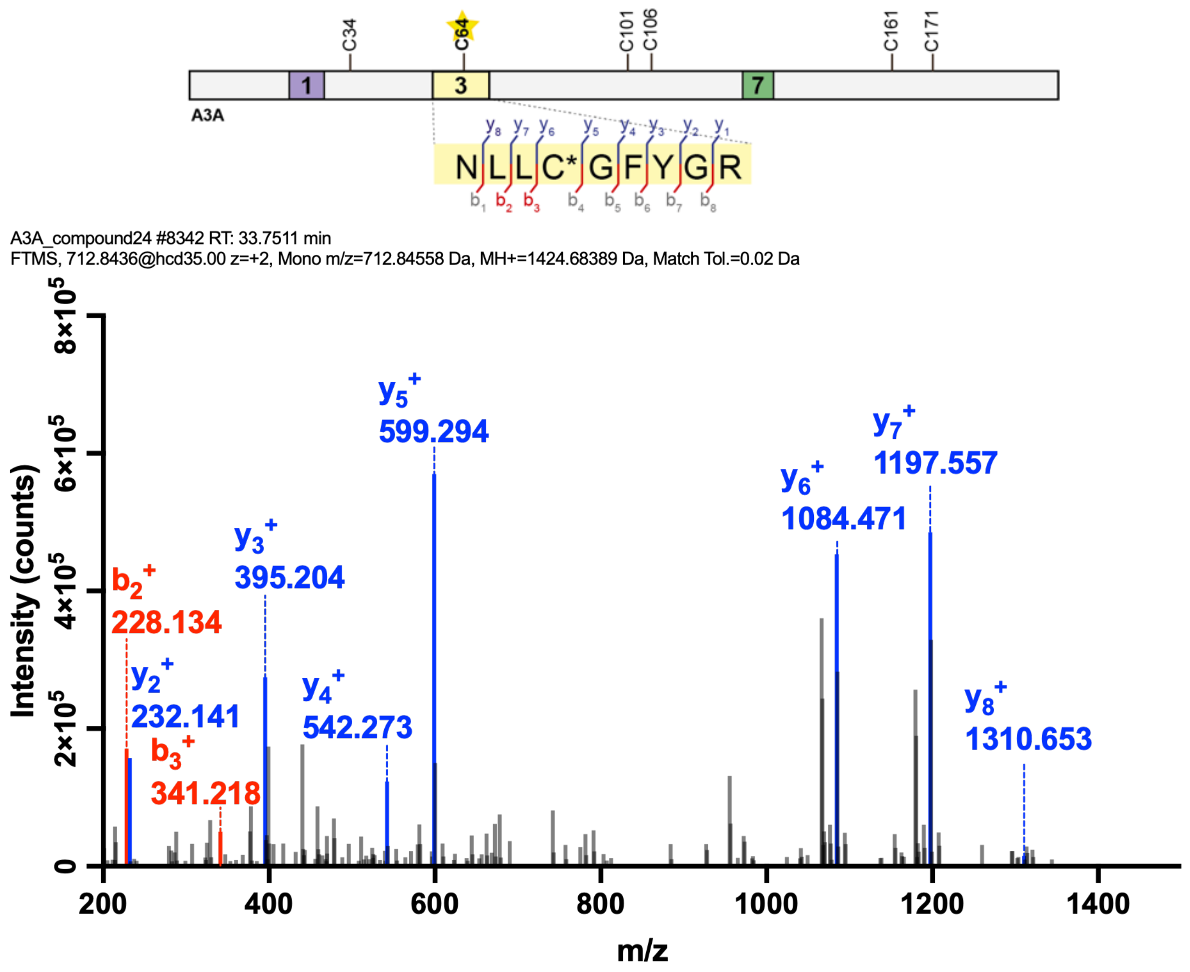


**Figure S9.** Collision-induced fragmentation pattern of major adducted peptide from A3A (25 µM) treated with 250 µM (10x) **24**, incubated for one hour. 32 peptide-spectrum matches (PSM) identified.

| $B^+$ | calculated | detected | sequence | $y^+$ | calculated | observed |
|---|---|---|---|---|---|---|
| 1 | 115.05020 | nd | N | 9 | - | - |
| 2 | 228.13427 | 228.13437 | L | 8 | 1310.64133 | 1310.63367 |
| 3 | 341.21833 | 341.21887 | L | 7 | 1197.55727 | 1197.55481 |
| 4 | 826.39792 | nd | C+adduct | 6 | 1084.47321 | 1084.47058 |
| 5 | 883.41938 | nd | G | 5 | 599.29362 | 599.29413 |
| 6 | 1030.48779 | nd | F | 4 | 542.27216 | 542.27344 |
| 7 | 1193.55112 | nd | Y | 3 | 395.20374 | 395.20432 |
| 8 | 1250.57259 | nd | G | 2 | 232.14042 | 232.14059 |
| 9 | - | - | R | 1 | 175.11895 | nd |

**Table S10.** Table of calculated and identified ions from **Figure S9.**

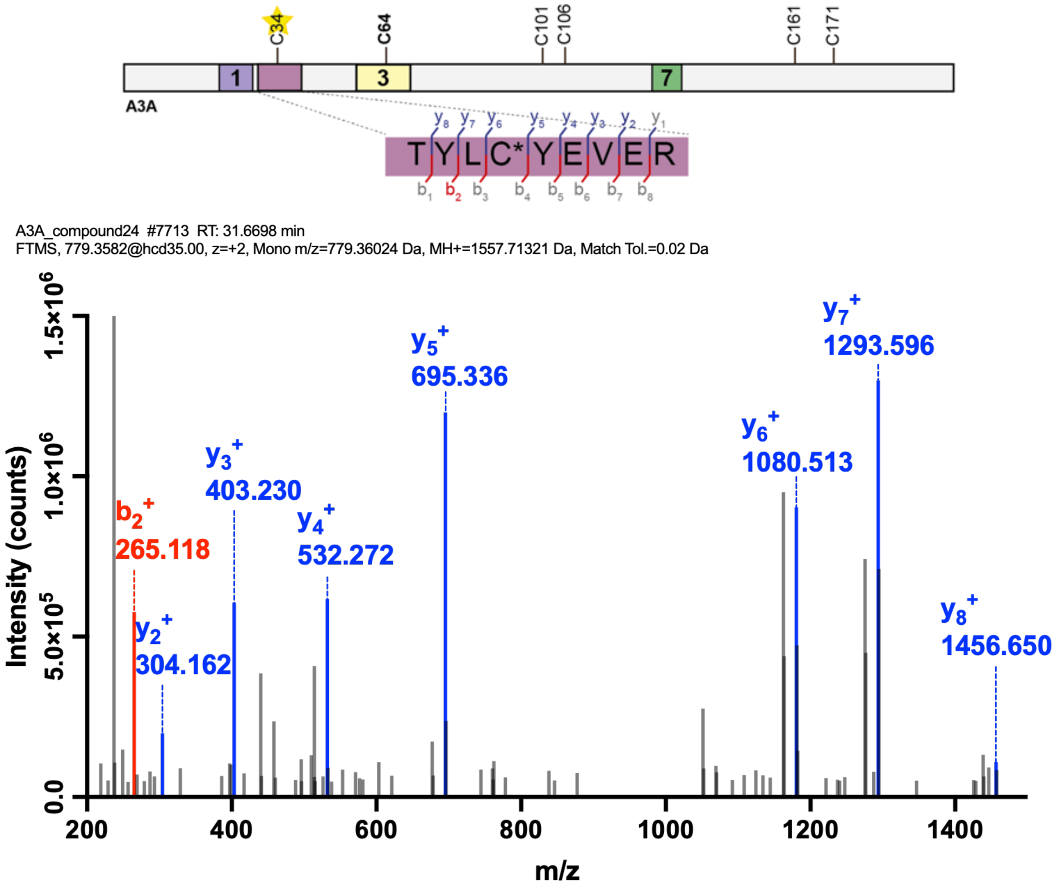


**Figure S10.** Collision-induced fragmentation pattern of minor adducted peptide from A3A (25 μM) treated with 250 μM (10x) **24**, incubated for one hour. 7 peptide-spectrum matches (PSM) identified.

| $B^+$ | calculated | detected | sequence | $y^+$ | calculated | observed |
|---|---|---|---|---|---|---|
| **1** | 102.05496 | nd | **T** | **9** | - | - |
| **2** | 265.11828 | 265.11844 | **Y** | **8** | 1456.66286 | 1456.65881 |
| **3** | 378.20235 | nd | **L** | **7** | 1293.59953 | 1293.59631 |
| **4** | 863.38193 | nd | **C+adduct** | **6** | 1180.51547 | 1180.51355 |
| **5** | 1026.44526 | nd | **Y** | **5** | 695.33588 | 695.33630 |
| **6** | 1155.48785 | nd | **E** | **4** | 532.27255 | 532.27203 |
| **7** | 1254.55627 | nd | **V** | **3** | 403.22996 | 403.23026 |
| **8** | 1383.59886 | nd | **E** | **2** | 304.16155 | 304.16165 |
| **9** | - | - | **R** | **1** | 175.11895 | nd |

**Table S11.** Table of calculated and identified ions from **Figure S10.**

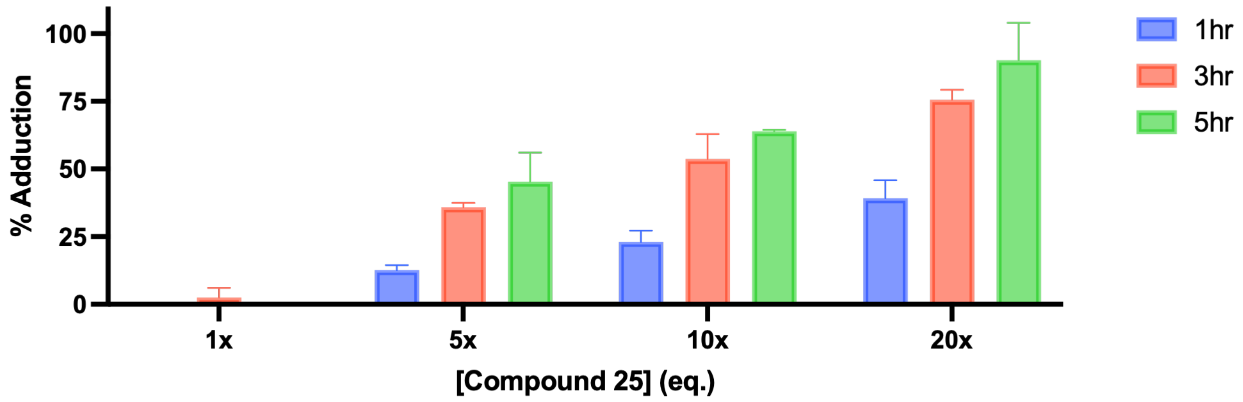


**Figure S11.** Bar graph of percent adduction of the acrylamide, **25**, to A3Bctd (25 μM) at 1x (25 μM), 5x (125 μM), 10x (250 μM), and 20x (500 μM). Percent adduction determined by intensity of mass species from deconvoluted spectra. N = 2 biological replicates, error bars represent standard deviation.

| MW of species (Da) | | Incubation = 1 hr | | | |
|---|---|---|---|---|---|
| | | 25 μM (1X) | 125 μM (5X) | 250 μM (10X) | 500 μM (20X) |
| 25338 no adduct | Replicate 1 | 100% | 89% | 80% | 65% |
| | Replicate 2 | 100% | 86% | 74% | 56% |
| | Average | 100% | 87.5% | 77% | 60.5% |
| 25735 (+397) | Replicate 1 | 0 | 11% | 20% | 35% |
| | Replicate 2 | 0 | 14% | 26% | 44% |
| | Average | 0 | 12.5% | 23% | 39.5% |

| MW of species (Da) | | Incubation = 3 hr | | | |
|---|---|---|---|---|---|
| | | 25 μM (1X) | 125 μM (5X) | 250 μM (10X) | 500 μM (20X) |
| 5338 no adduct | Replicate 1 | 100% | 63% | 53% | 27% |
| | Replicate 2 | 95% | 65% | 40% | 22% |
| | Average | 97.5% | 64% | 46.5% | 24.5% |
| 25735 (+397) | Replicate 1 | 0 | 37% | 47% | 73% |
| | Replicate 2 | 5% | 35% | 60% | 78% |
| | Average | 2.5% | 36% | 53.5% | 75.5% |

| MW of species (Da) | | Incubation = 5 hr | | | |
|---|---|---|---|---|---|
| | | 25 μM (1X) | 125 μM (5X) | 250 μM (10X) | 500 μM (20X) |
| 25339 no adduct | Replicate 1 | 100% | 47% | 36% | 0 |
| | Replicate 2 | 100% | 62% | 36% | 20% |
| | Average | 100% | 54.5% | 36% | 10% |
| 25735 (+397) | Replicate 1 | 0 | 53% | 64% | 100% |
| | Replicate 2 | 0 | 38% | 64% | 80% |
| | Average | 0 | 45.5% | 64% | 90% |

**Table S12.** Average percent adduction of A3Bctd treated with **25** at variable concentration and time, from intact protein mass spectrometry deconvolution.

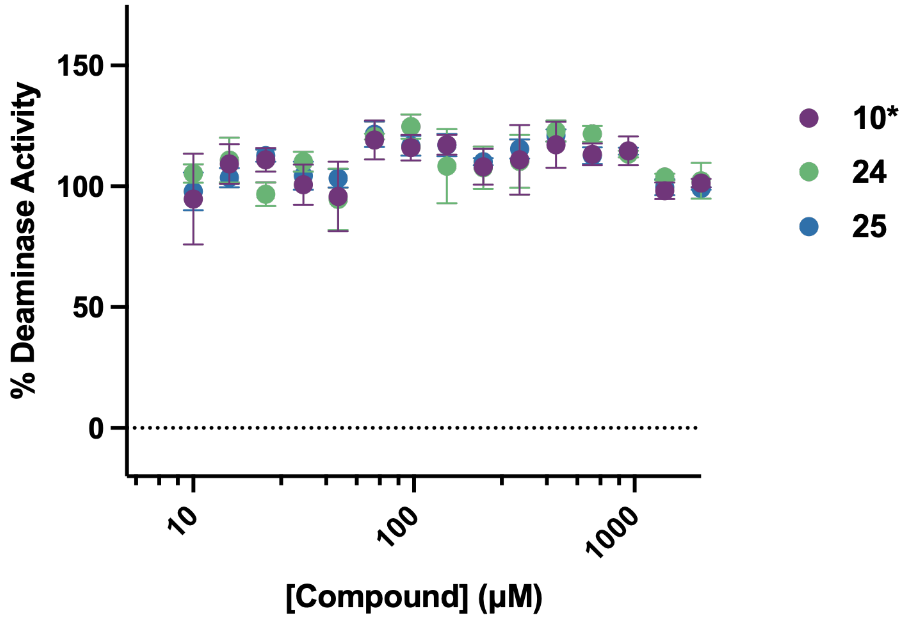


**Figure S12.** Plate-based deaminase assay of resynthesized compound **10** (**10***), and cysteine probes (**24** and **25**) against A3A. Error bars are standard error of the mean (SEM) of three technical replicates.

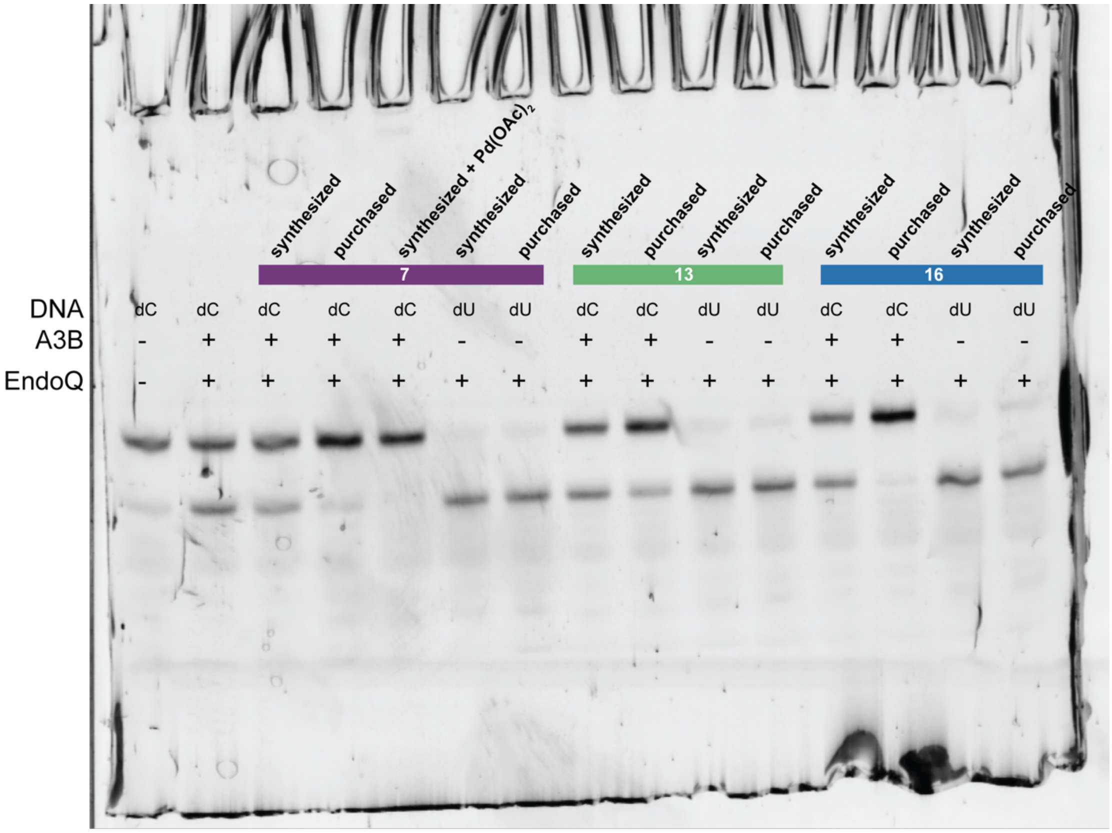


**Figure S13**. First replicate of uncropped and labeled deaminase gel of purchased, synthesized, and synthesized stocks supplemented at 0.1% $Pd(OAc)_2$ tested against A3Bctd.

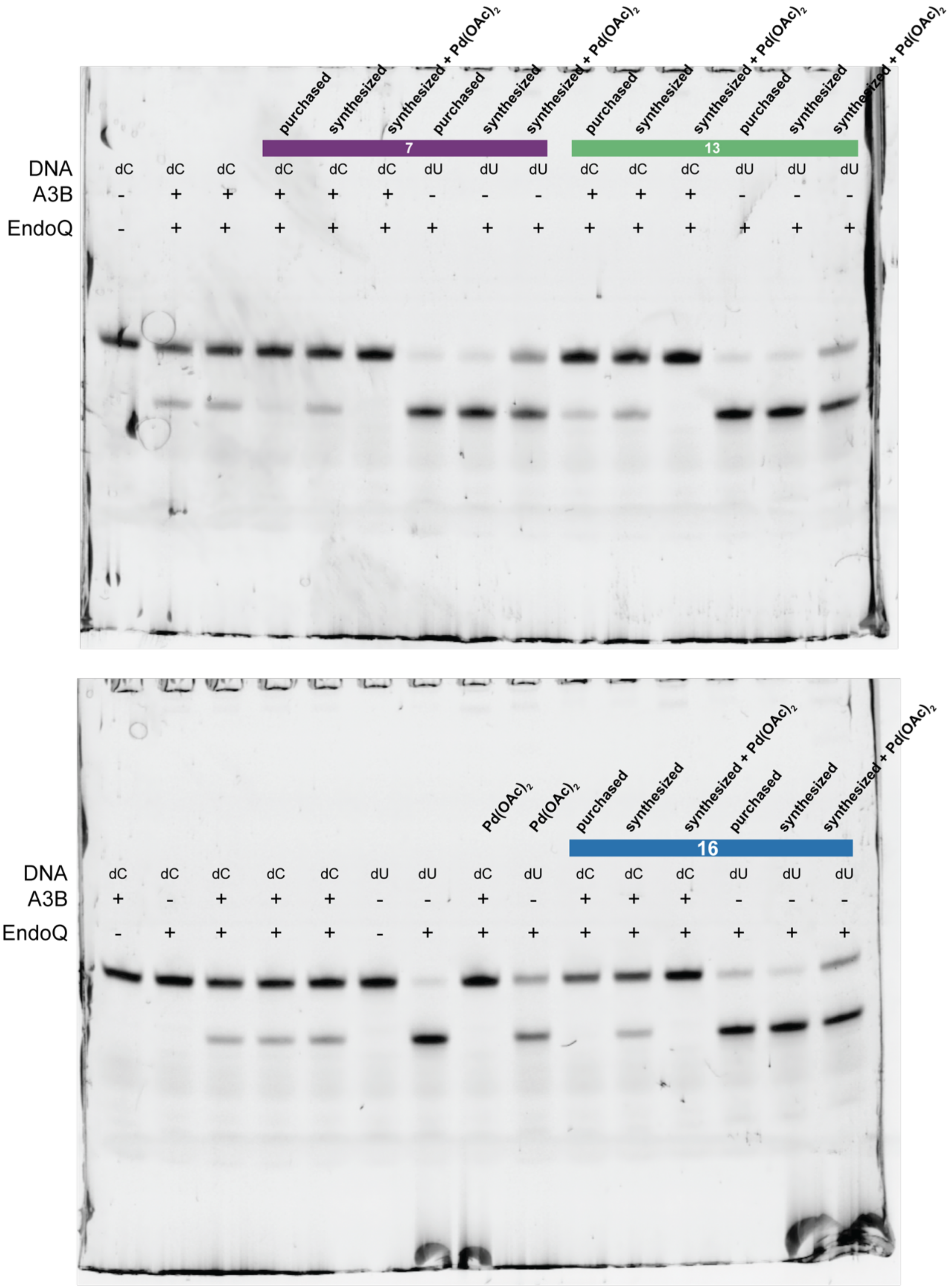


**Figure S14**. Second replicate of uncropped and labeled deaminase gel of purchased, synthesized, and synthesized stocks supplemented at 0.1% $Pd(OAc)_2$ tested against A3Bctd.

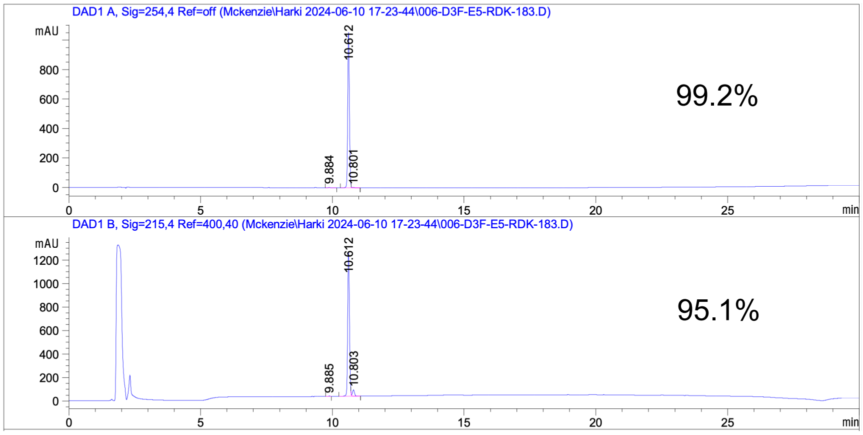


**Figure S15**. HPLC analysis of resynthesized **7.**

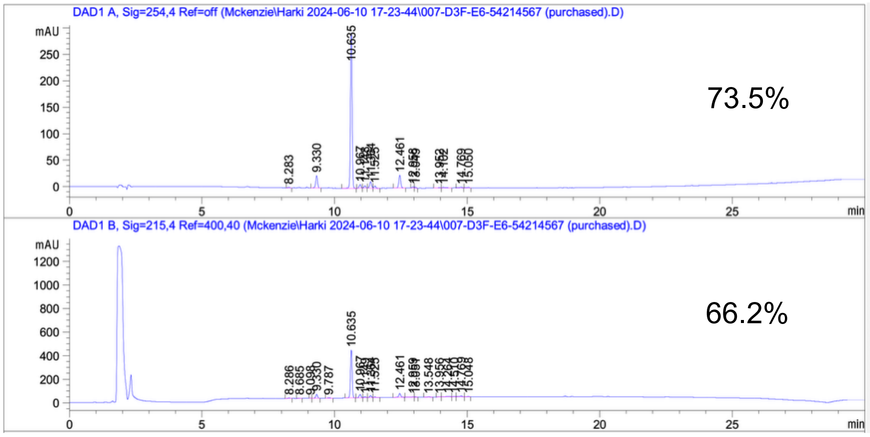


**Figure S16**. HPLC analysis of purchased **7.**

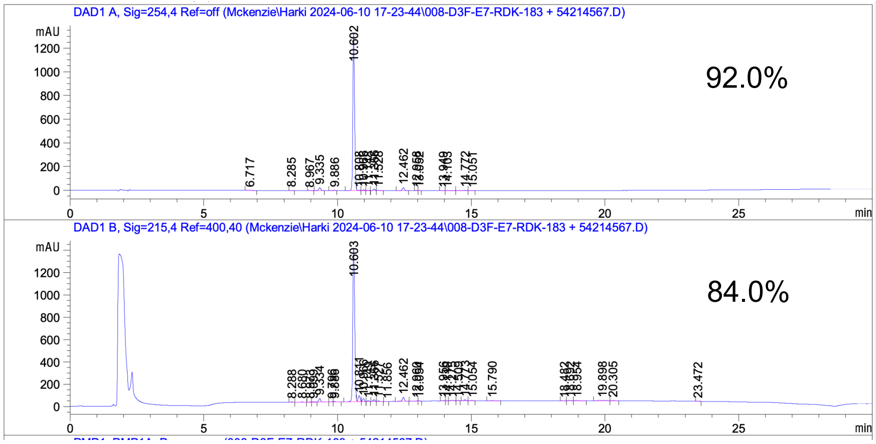


**Figure S17**. HPLC analysis of co-injection of purchased and resynthesized **7.**

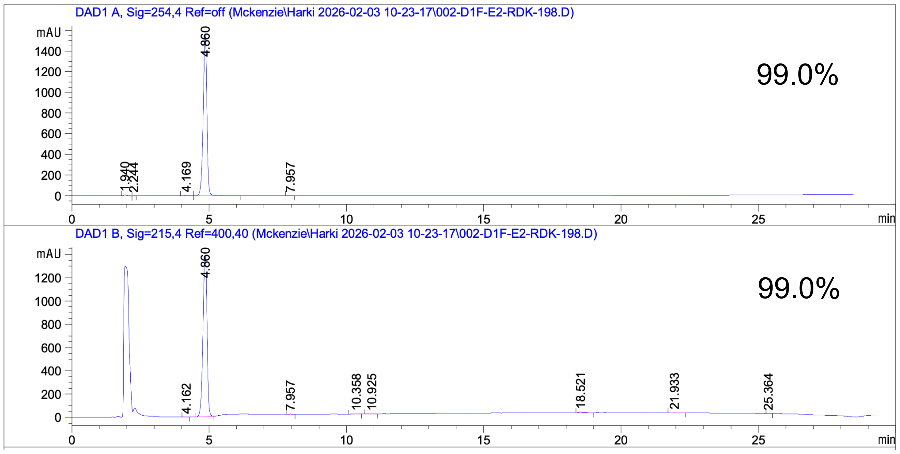


**Figure S18.** HPLC analysis of synthesized **13**.

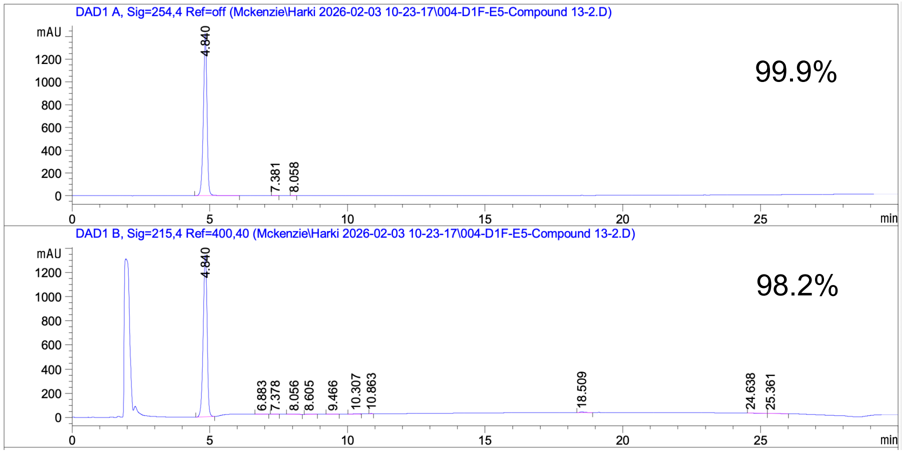


**Figure S19**. HPLC analysis of purchased **13**.

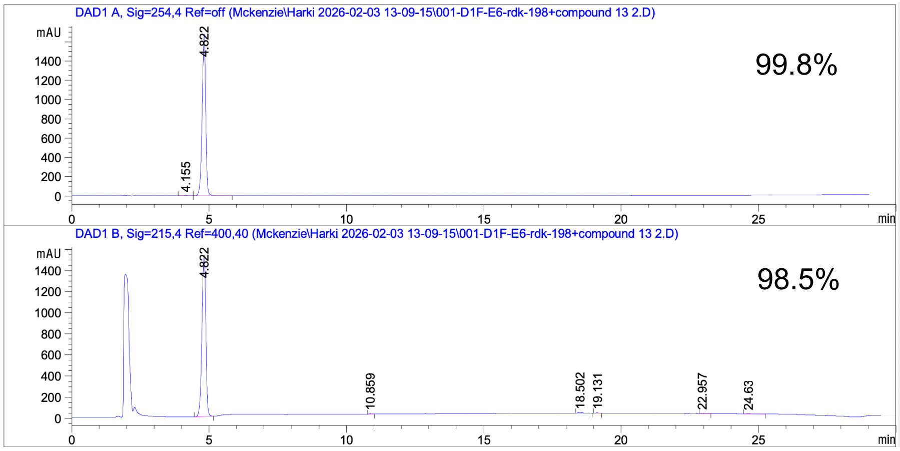


**Figure S20**. HPLC analysis of co-injection of purchased and resynthesized **13.**

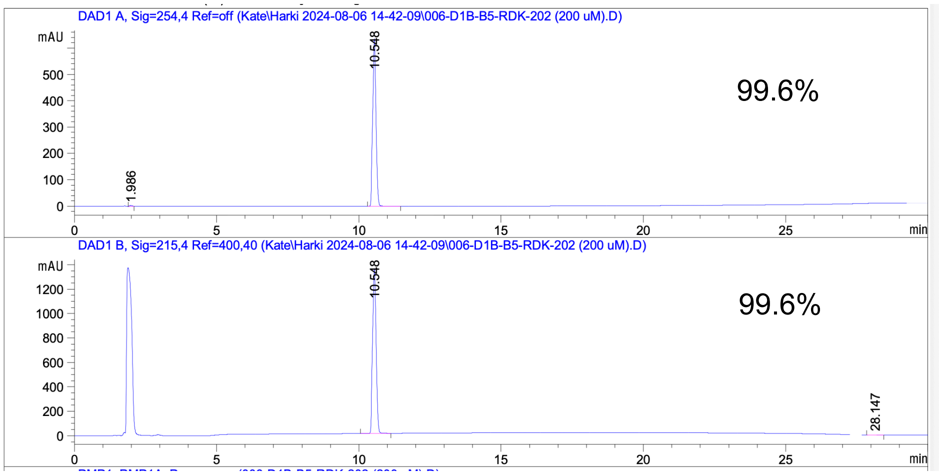


**Figure S21**. HPLC analysis of synthesized **16.**

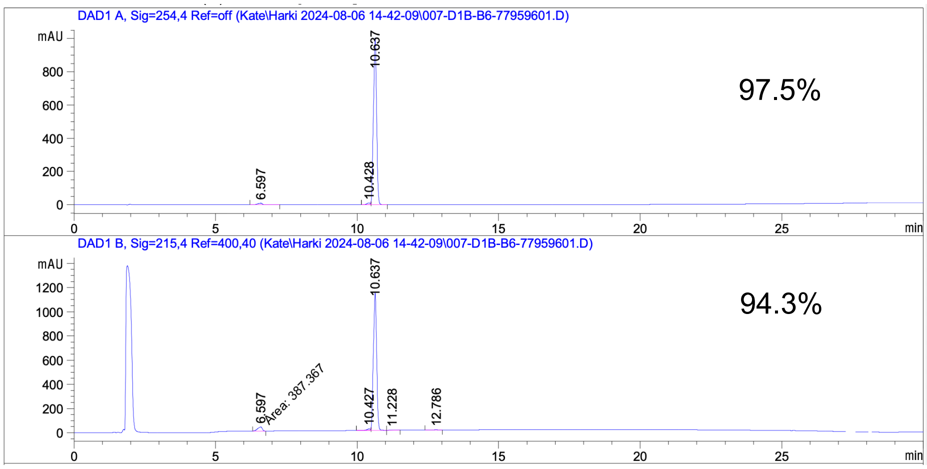


**Figure S22**. HPLC analysis of purchased **16.**

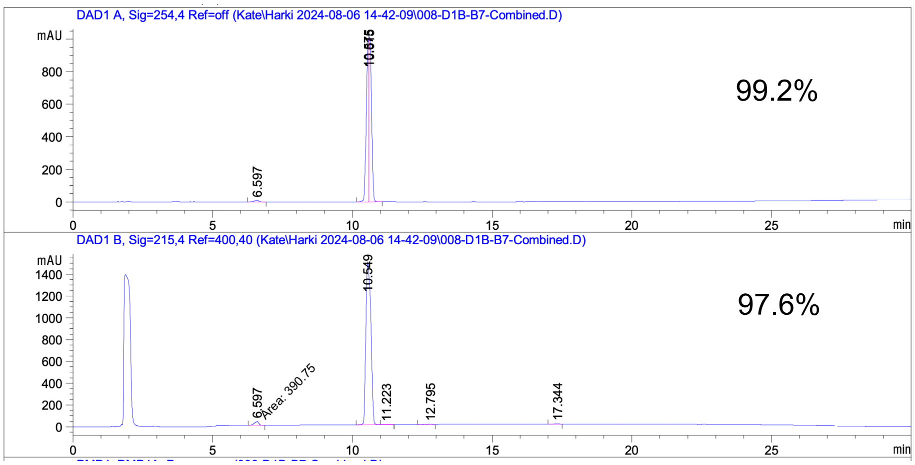


**Figure S23**. HPLC analysis of co-injection of synthesized and purchased **16.**

| Compound | Stock concentration | Co (ppb) | Cu (ppb) | Zn (ppb) | Pd (ppb) | Pt (ppb) | Au (ppb) |
|---|---|---|---|---|---|---|---|
| 7 (purchased) | 40 mM | 3.000 | 54.000 | 4220 | 144 | <BEC | <BEC |
| 7 (synthesized) | 40 mM | 1.000 | 28.000 | 515 | 295 | <BEC | <BEC |
| 13 (purchased) | 100 mM | 1.000 | 2.000 | 74 | 72 | <BEC | <BEC |
| 13 (synthesized) | 50 mM | 0.000 | 6.000 | 159 | 105 | <BEC | <BEC |
| 16 (purchased) | 20 mM | 0.000 | 8.000 | 147 | 8980 | <BEC | <BEC |
| 16 (synthesized) | 20 mM | 1.000 | 28.000 | 74 | 1729 | <BEC | <BEC |

**Table S13.** Results of the trace metal analysis by ICP-MS. Results are in parts-per-billion (ppb), which is equivalent to µg of analyte per L of the as-received liquid sample. Results reported as less than background equivalent concentration (<BEC) are less than 0.004 and 0.058 ppb for Pt and Au, respectively.

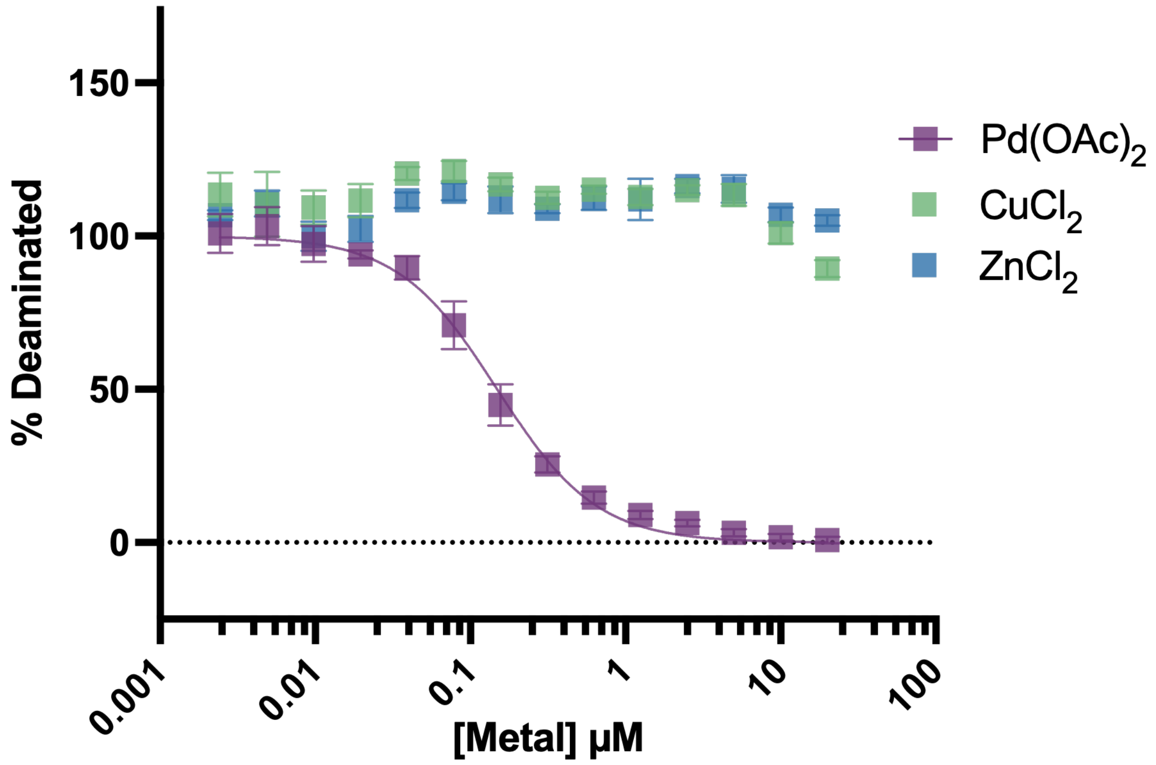


**Figure S24**. Plate-based deaminase assay of $Pd(OAc)_2$, $CuCl_2$, and $ZnCl_2$ against A3A. Graph is one representative replicate of two biological replicates. Error bars are the standard error of the mean (SEM) of three technical replicates. $IC_{50}$ value for $Pd(OAc)_2$ is 139 nM or 31 ppb.

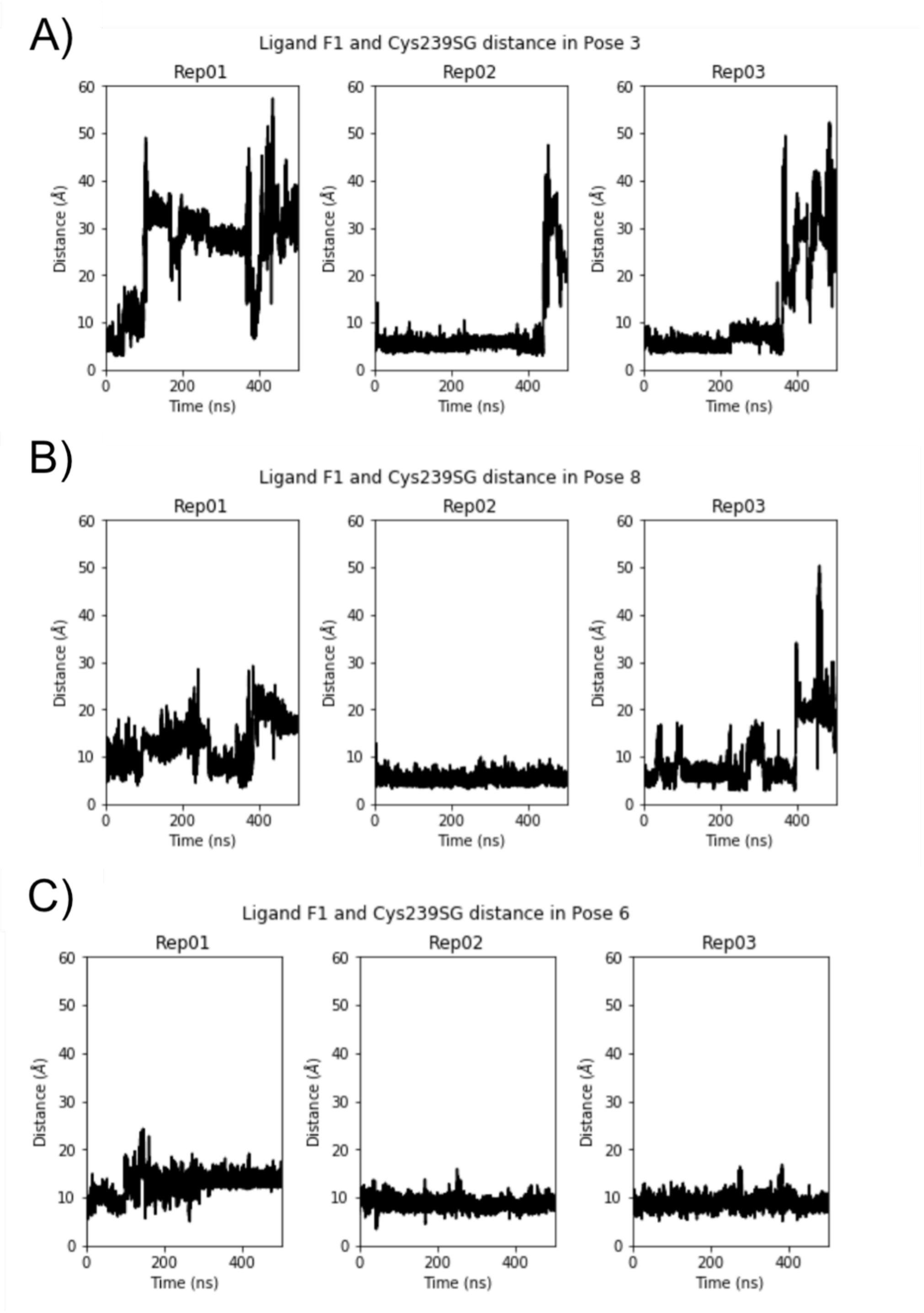


**Figure S25**. Initial binding pose stabilities between A3B-ctd and allosteric ligand **10**. Distance between ligand 4-fluoride atom and C239 SG atom of A3Bctd through 100ns of A) pose3, B) pose 8, and C) pose 6. These initial simulations and distance metrics were used evaluate the stability of the ligand in the allosteric pocket, which determined the final binding pose (pose6) to be used in further simulations.

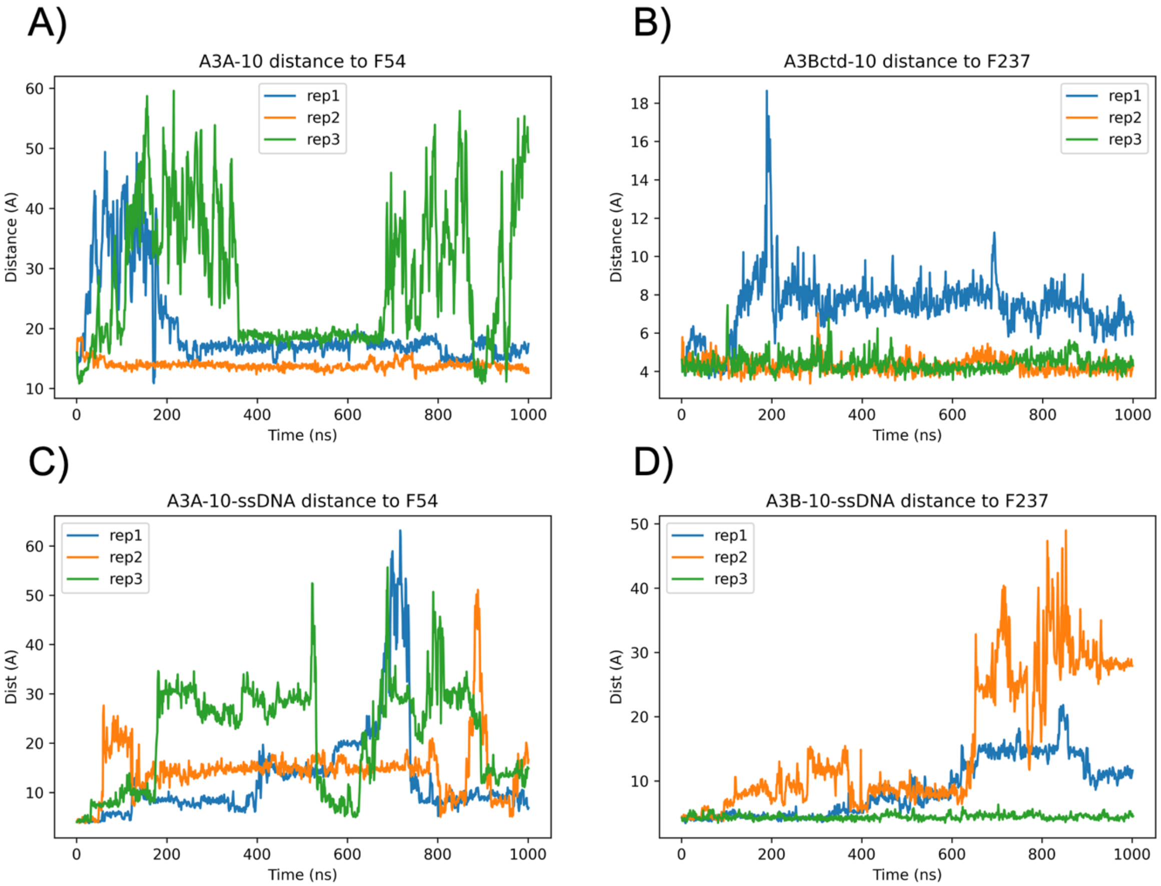


**Figure S26.** Distances between **10** and allosteric pocket COM (F237/F54 in A3Bctd and A3A respectively). Panels A & B show simulations of protein and **10** when ssDNA is not present. Panels C & D show simulations of protein, **10**, and ssDNA.

#### 3.0 Characterization Spectra

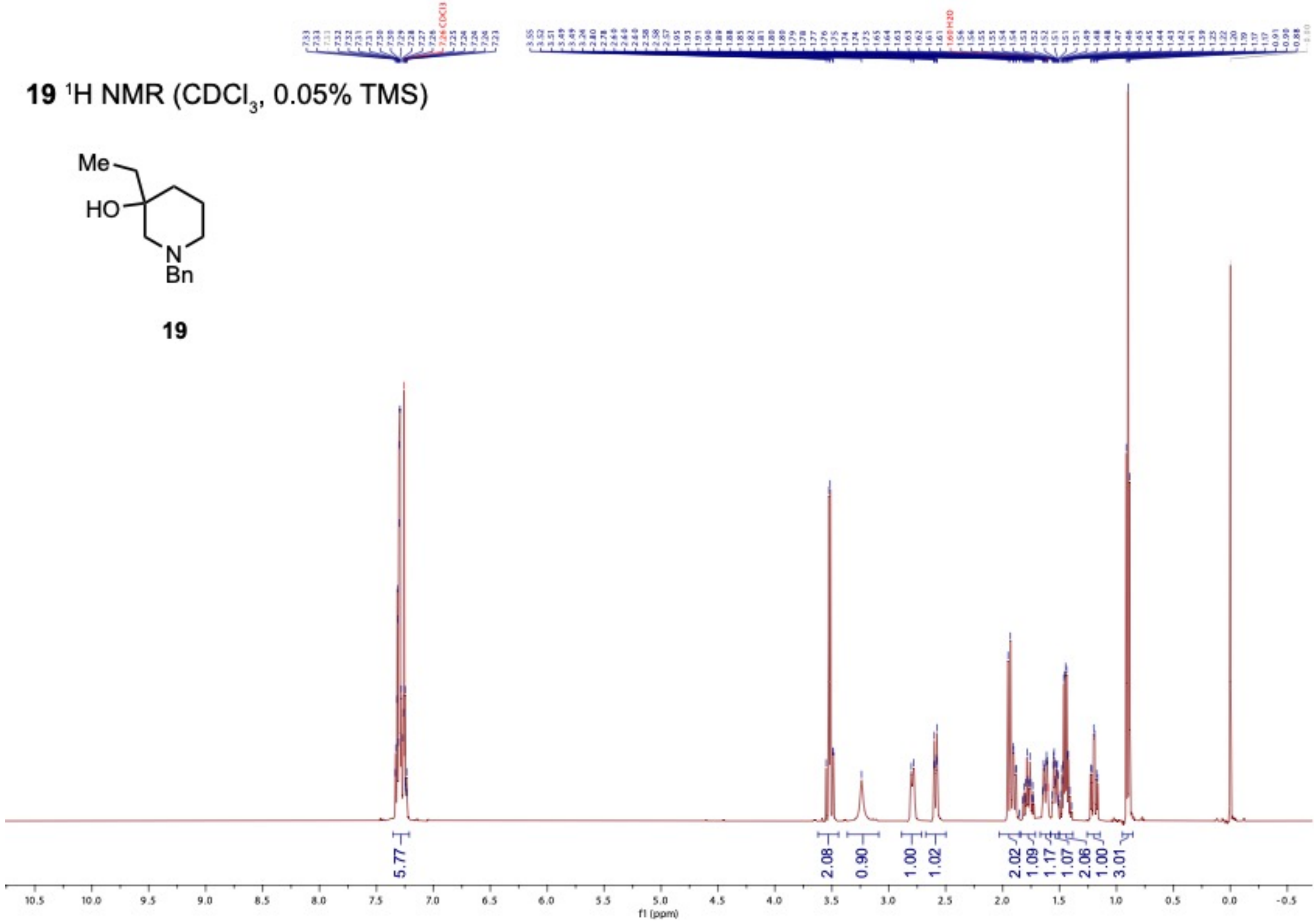

**20** $^{1}H$ NMR ($CD_3OD$, 0.05% TMS)

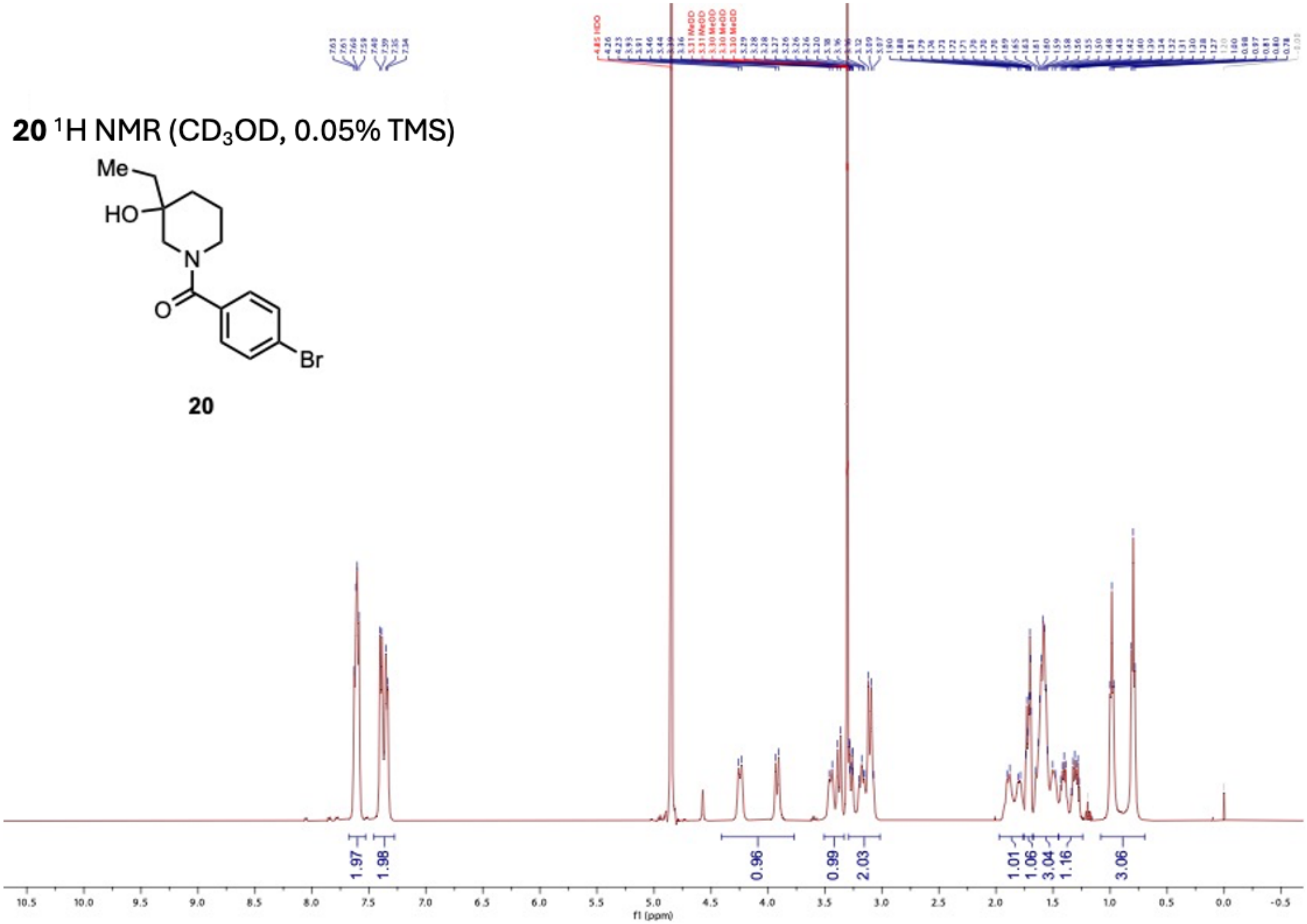


**20**

**20** $^{13}C$ NMR ($CD_3OD$)

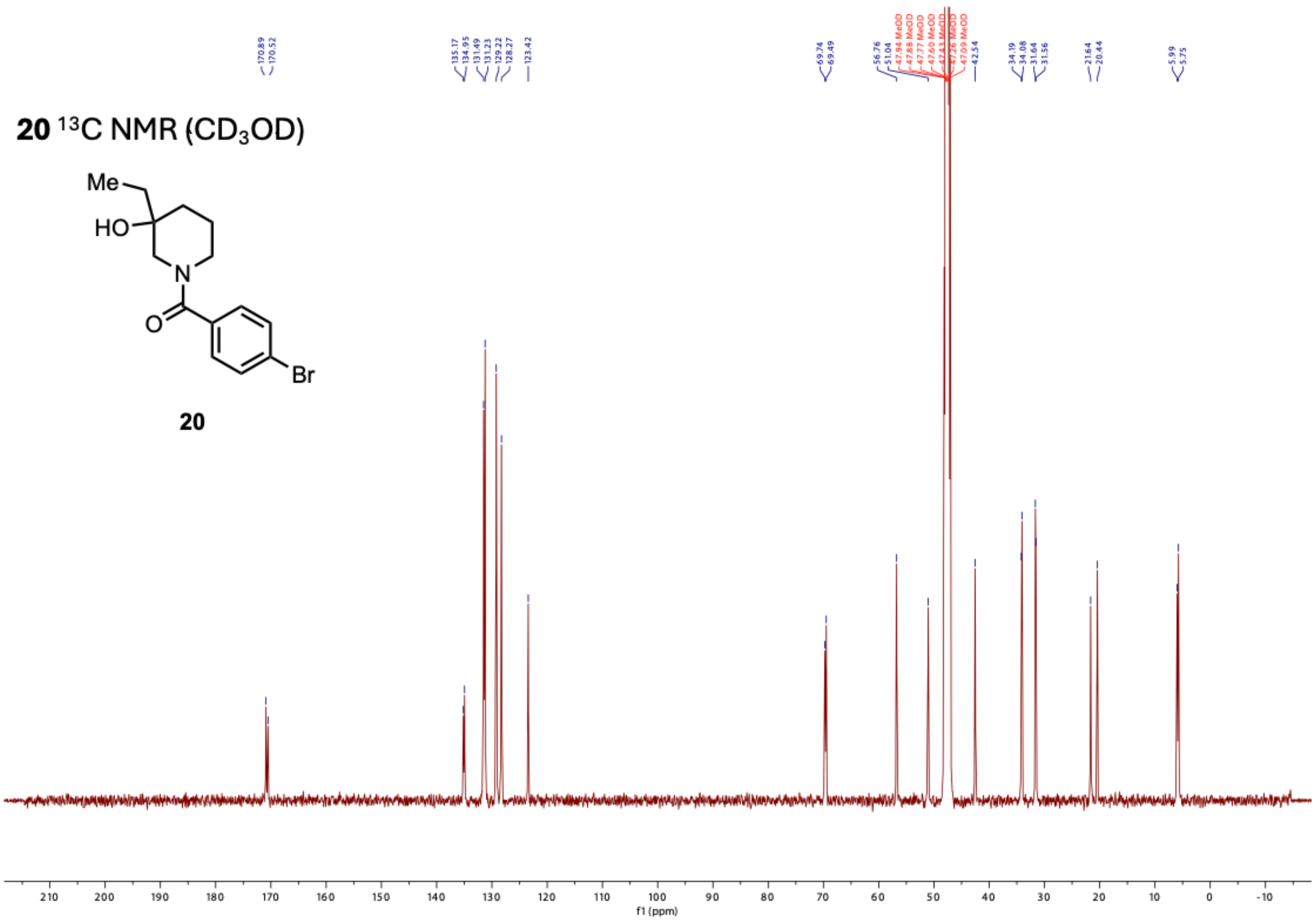


**20**

**10** $^{1}$H NMR (DMSO-$d_6$)

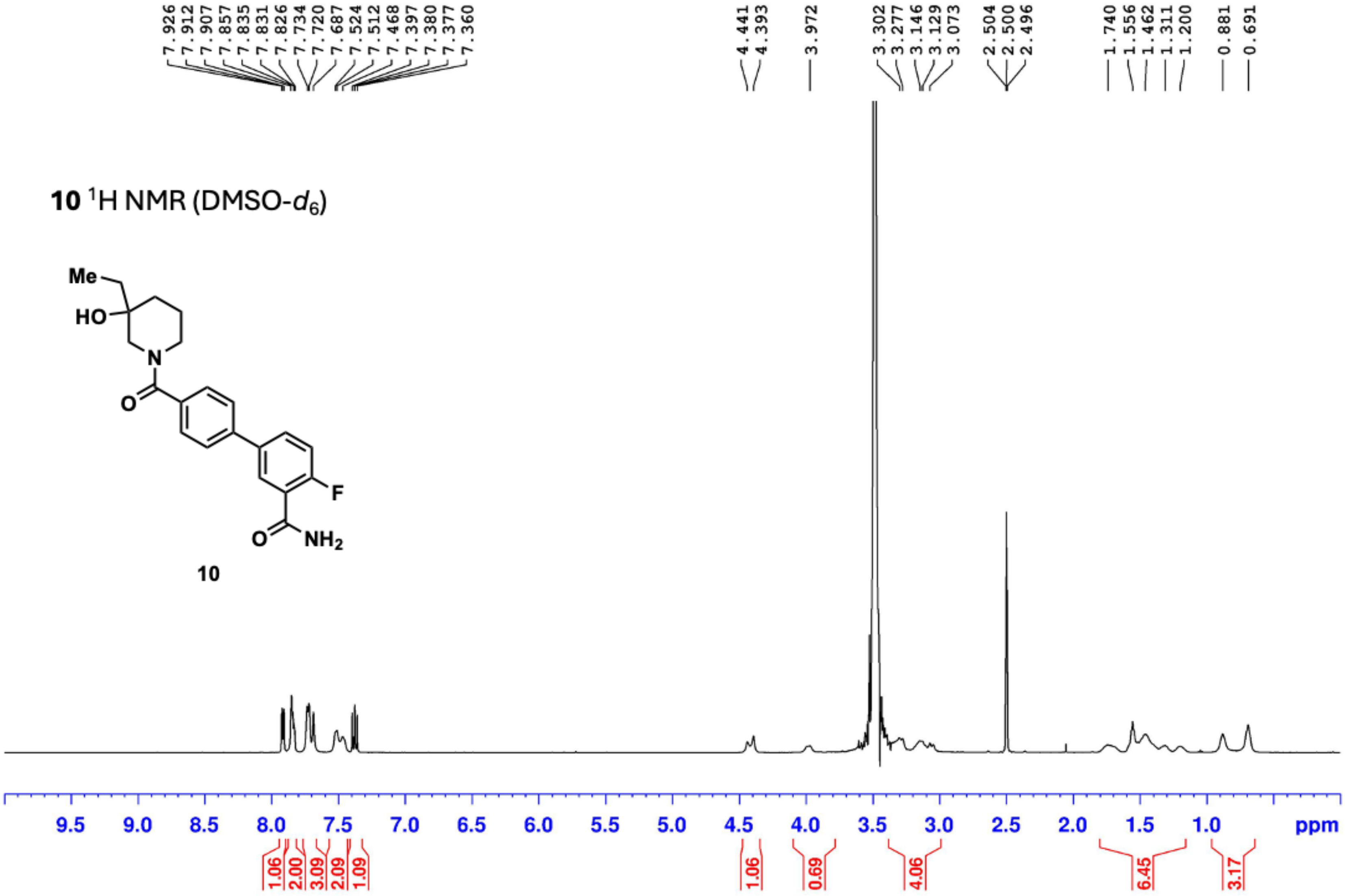


**10**

**10** $^{13}$C NMR (DMSO-$d_6$)

**10**

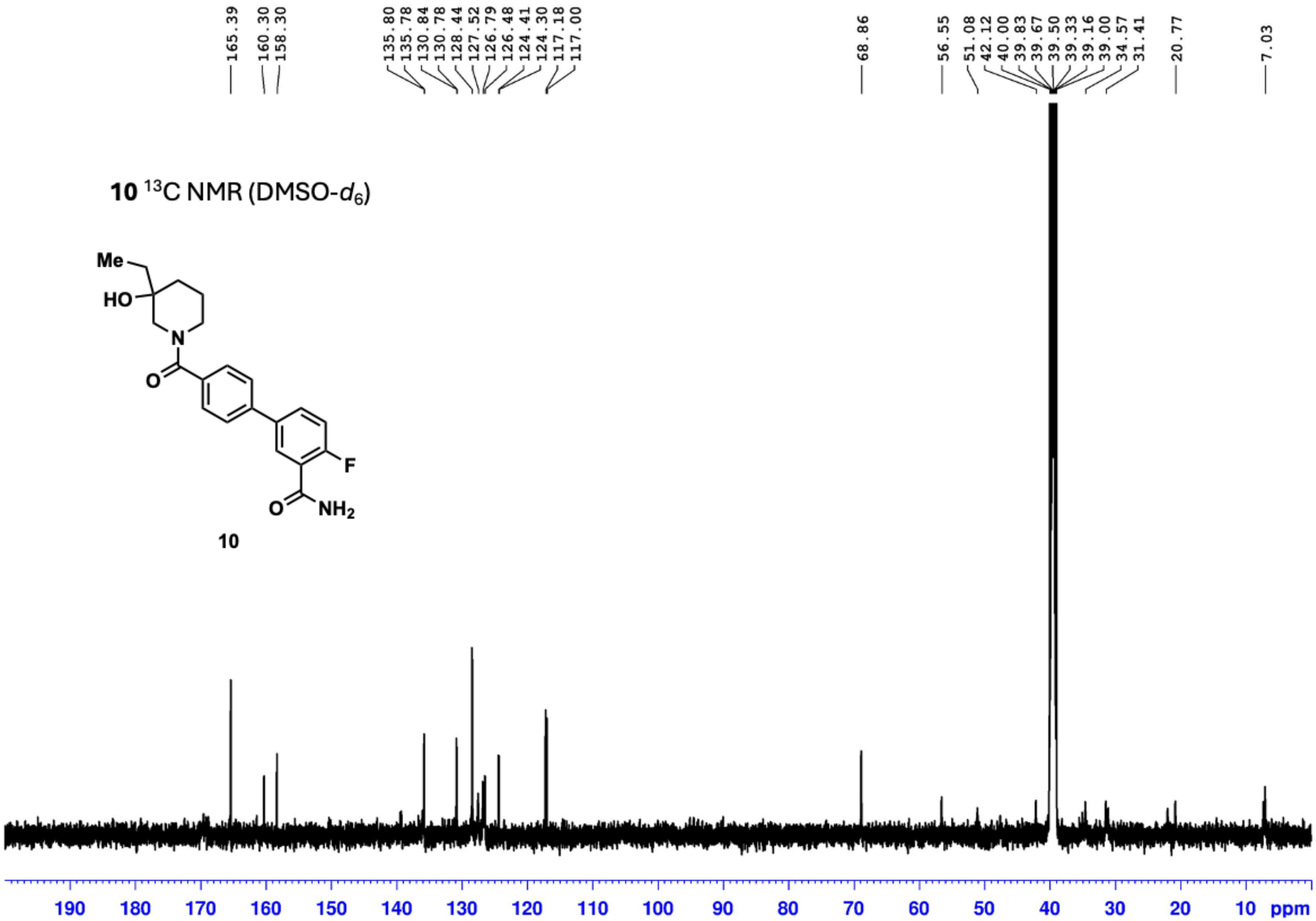

**23** $^{1}$H NMR ($CDCl_3$)

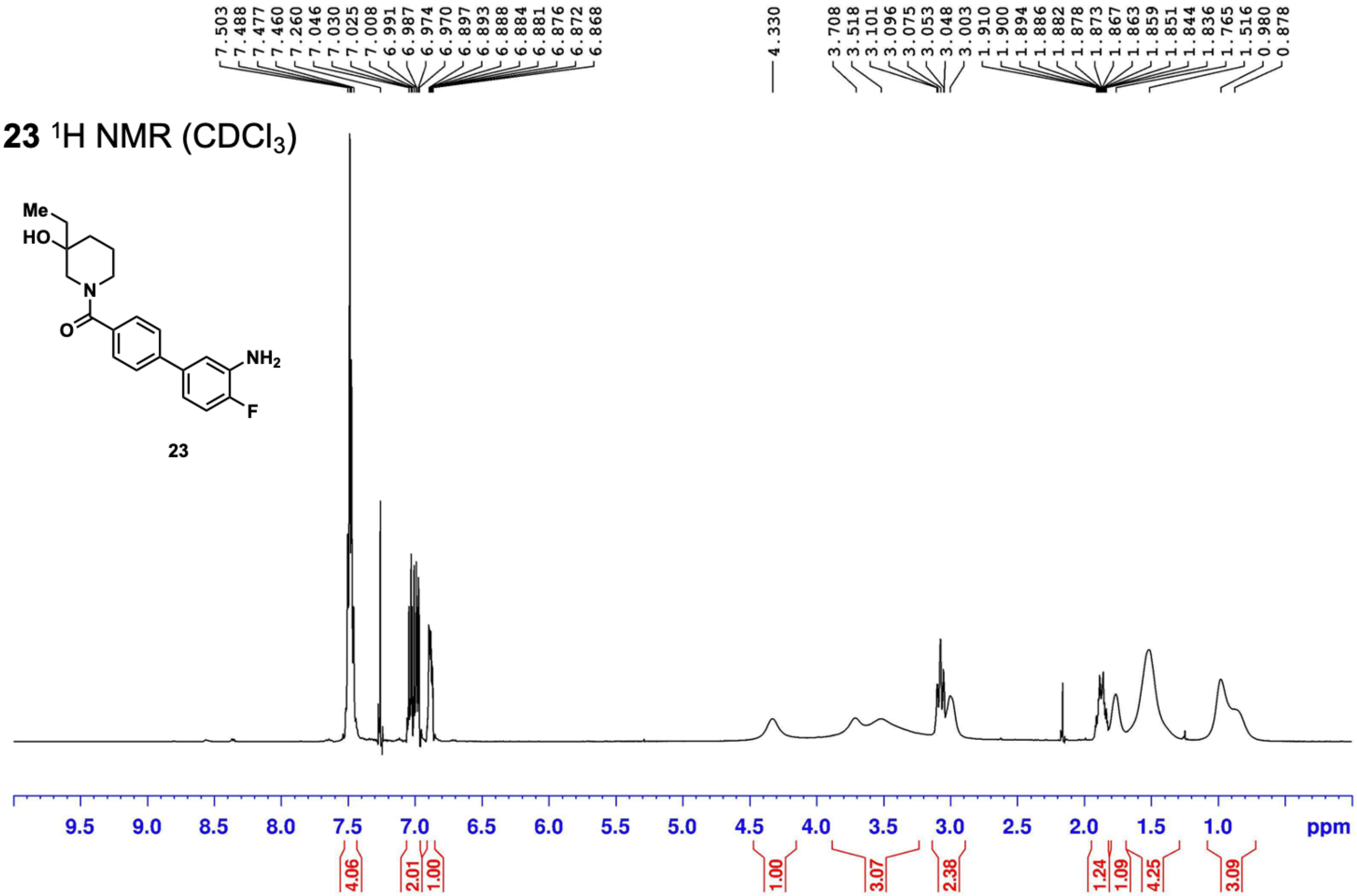

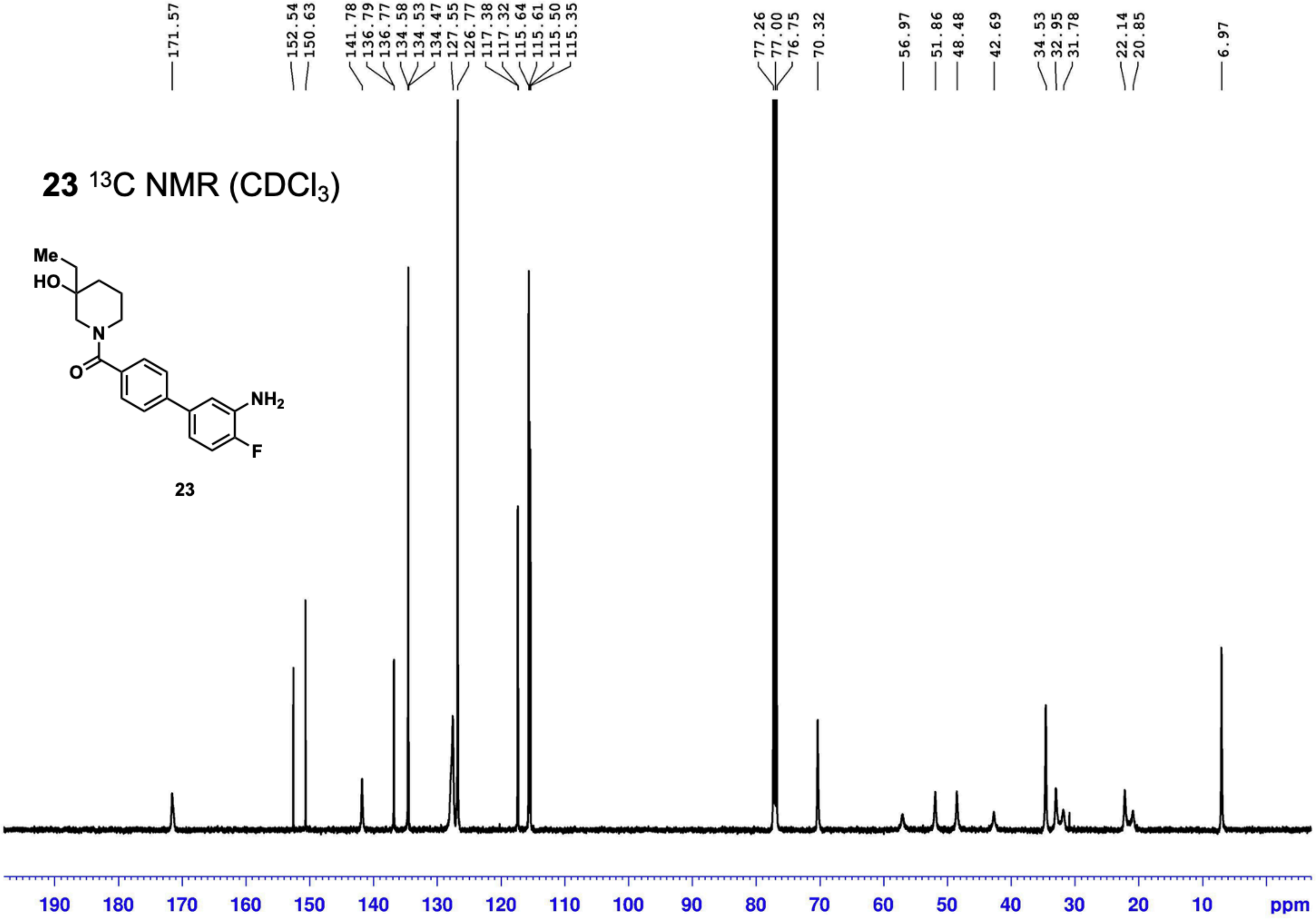
23 $^{13}C$ NMR ($CDCl_3$)
Me
HO
N
O
NH2
F
23
171.57
152.54
150.63
141.78
136.79
136.77
134.58
134.53
134.47
127.55
126.77
117.38
117.32
115.64
115.61
115.50
115.35
77.26
77.00
76.75
70.32
56.97
51.86
48.48
42.69
34.53
32.95
31.78
22.14
20.85
6.97
190 180 170 160 150 140 130 120 110 100 90 80 70 60 50 40 30 20 10 ppm

**23** $^{19}$F NMR ($CDCl_3$)

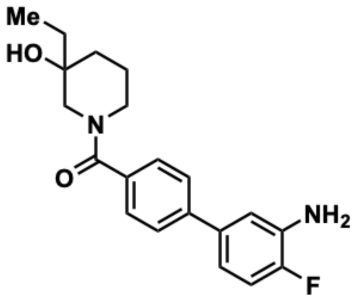


**23**

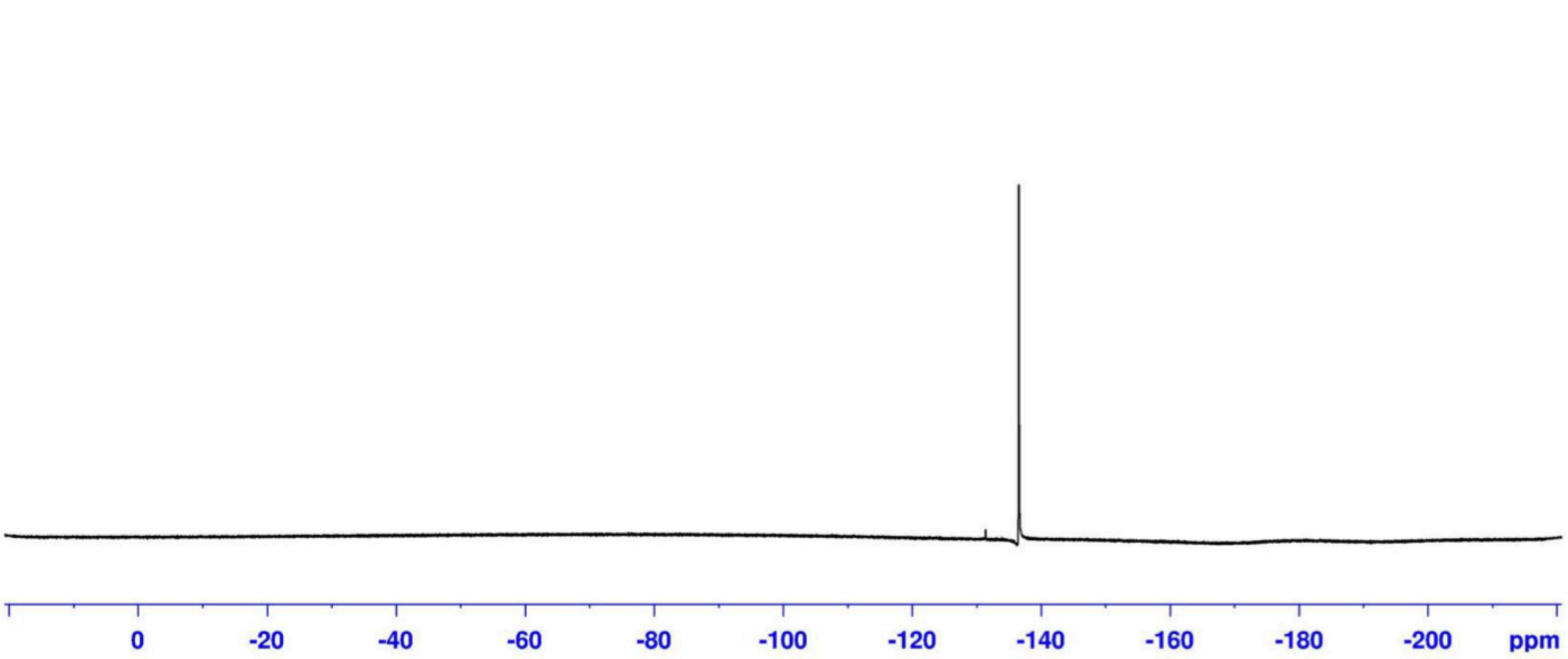

**24** [1]H NMR ($CD_3OD$)

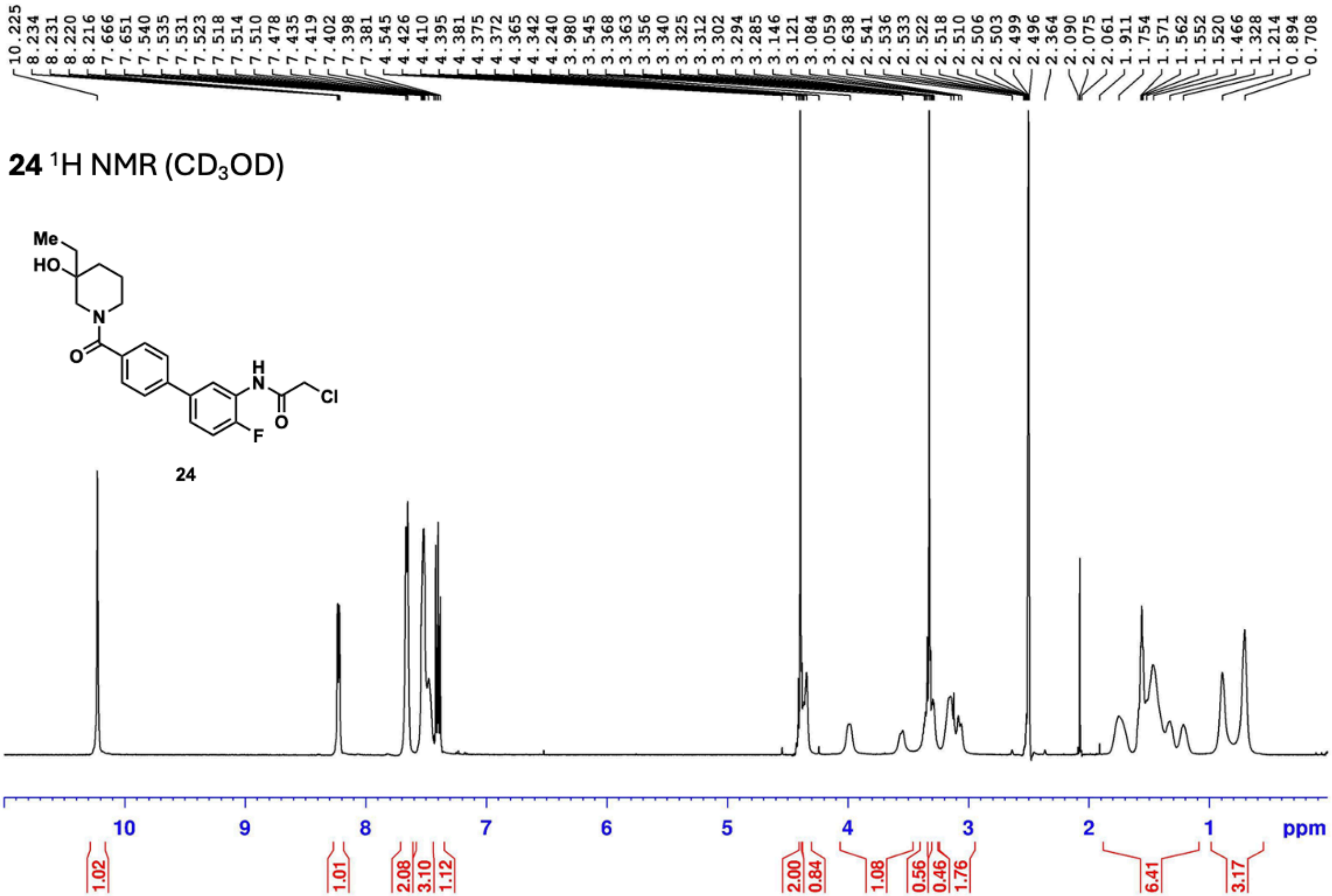

**24** $^{13}$C NMR (DMSO-$d_6$)

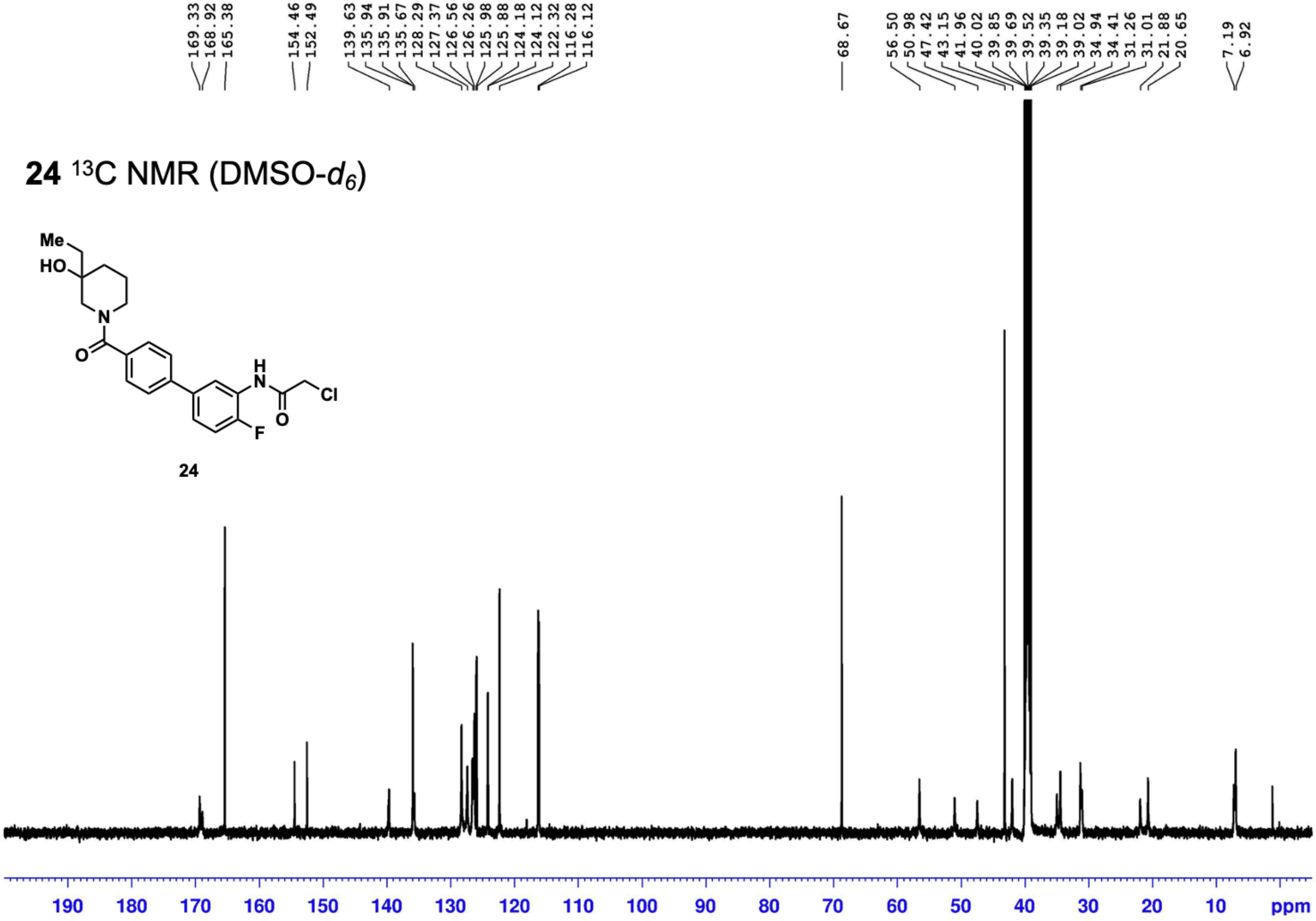

24 19F NMR (CDCl3)

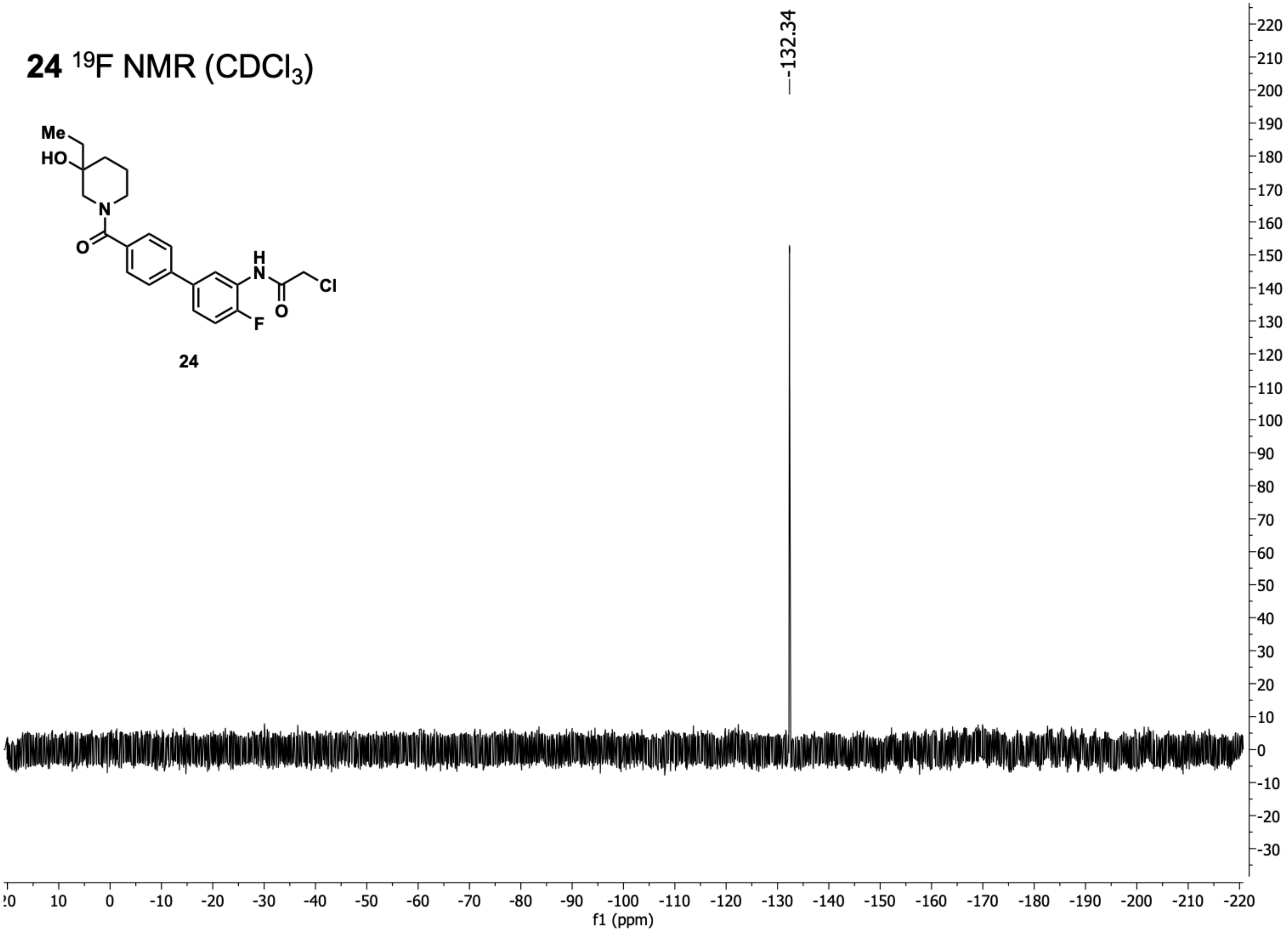
Me
HO
N
O
H
N
Cl
O
F
24
-132.34
f1 (ppm)

**25** $^{1}$H NMR ($CDCl_3$)

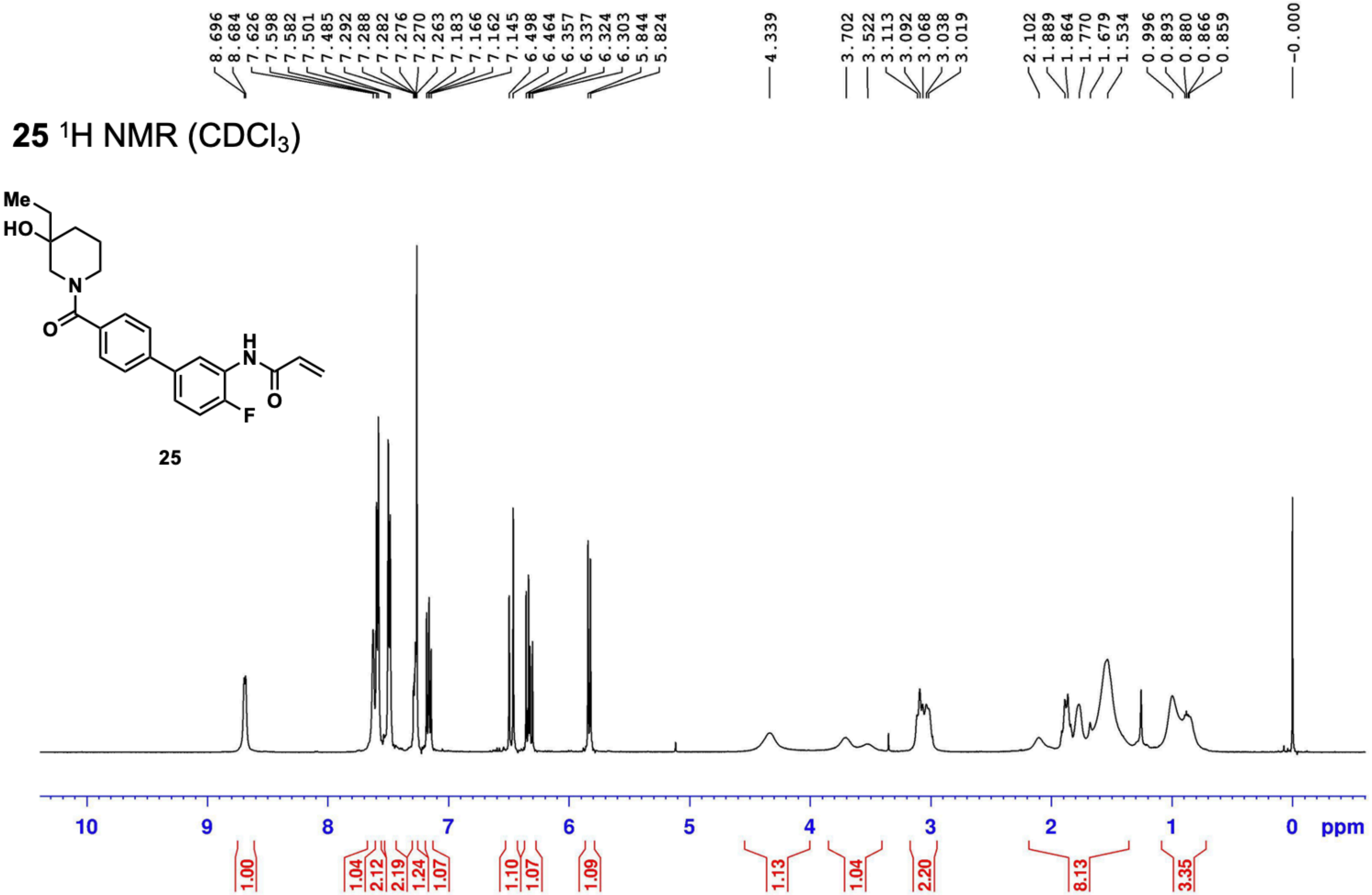

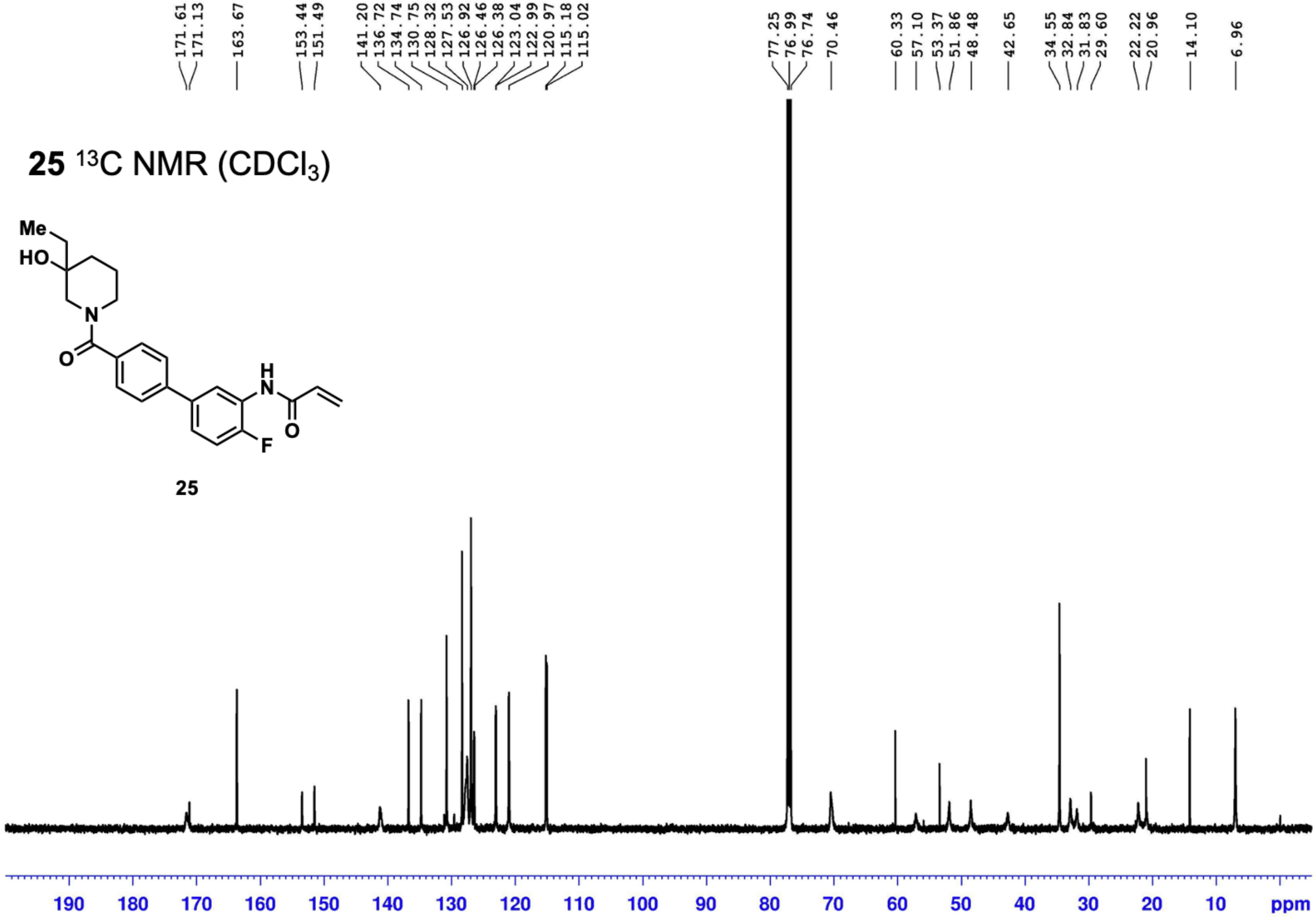
25 $^{13}$C NMR ($CDCl_3$)
171.61
171.13
163.67
153.44
151.49
141.20
136.72
134.74
130.75
128.32
127.53
126.92
126.46
126.38
123.04
122.99
120.97
115.18
115.02
77.25
76.99
76.74
70.46
60.33
57.10
53.37
51.86
48.48
42.65
34.55
32.84
31.83
29.60
22.22
20.96
14.10
6.96
Me
HO
N
O
H
N
O
F
25
190 180 170 160 150 140 130 120 110 100 90 80 70 60 50 40 30 20 10 ppm

**25** $^{19}F$ NMR ($CDCl_3$)

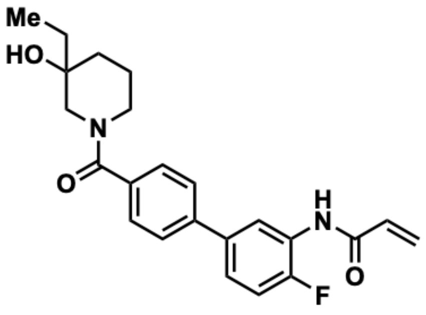

**25**

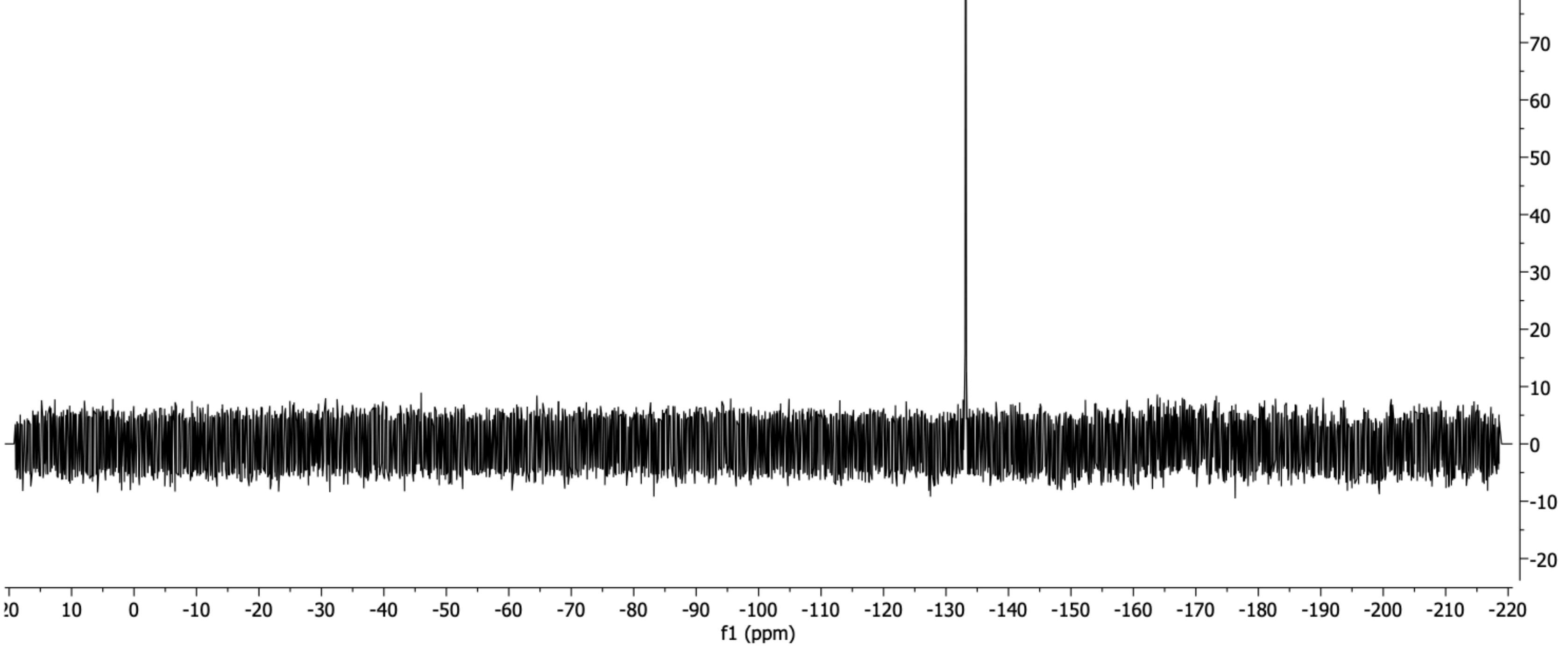

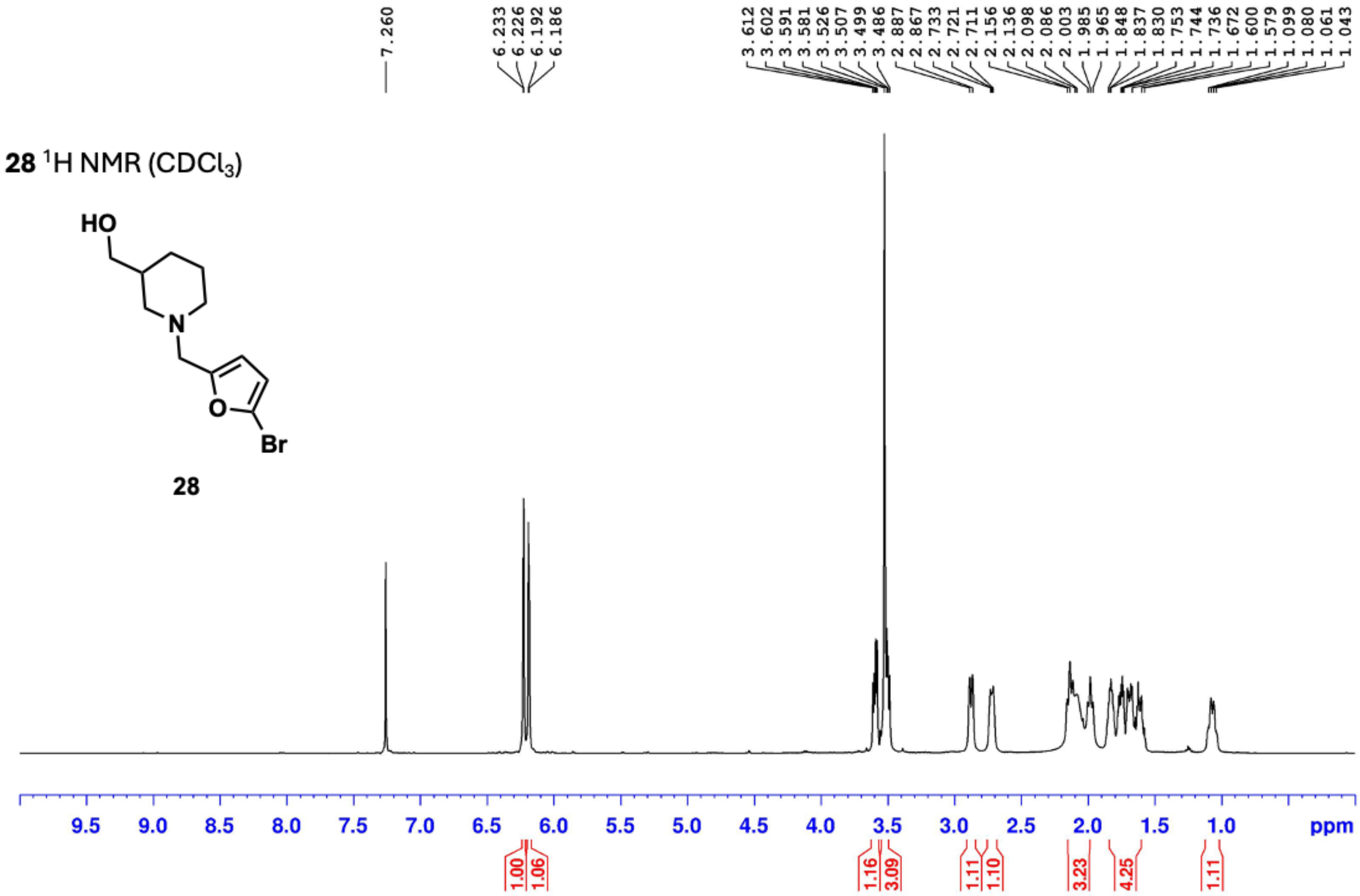
28 ¹H NMR (CDCl3)
HO
N
O
Br
28
7.260
6.233
6.226
6.192
6.186
3.612
3.602
3.591
3.581
3.526
3.507
3.499
3.486
2.887
2.867
2.733
2.721
2.711
2.156
2.136
2.098
2.086
2.003
1.985
1.965
1.848
1.837
1.830
1.753
1.744
1.736
1.672
1.600
1.579
1.099
1.080
1.061
1.043
9.5
9.0
8.5
8.0
7.5
7.0
6.5
6.0
5.5
5.0
4.5
4.0
3.5
3.0
2.5
2.0
1.5
1.0
ppm
1.00
1.06
1.16
3.09
1.11
1.10
3.23
4.25
1.11

**28** $^{1}$H NMR ($CDCl_3$)

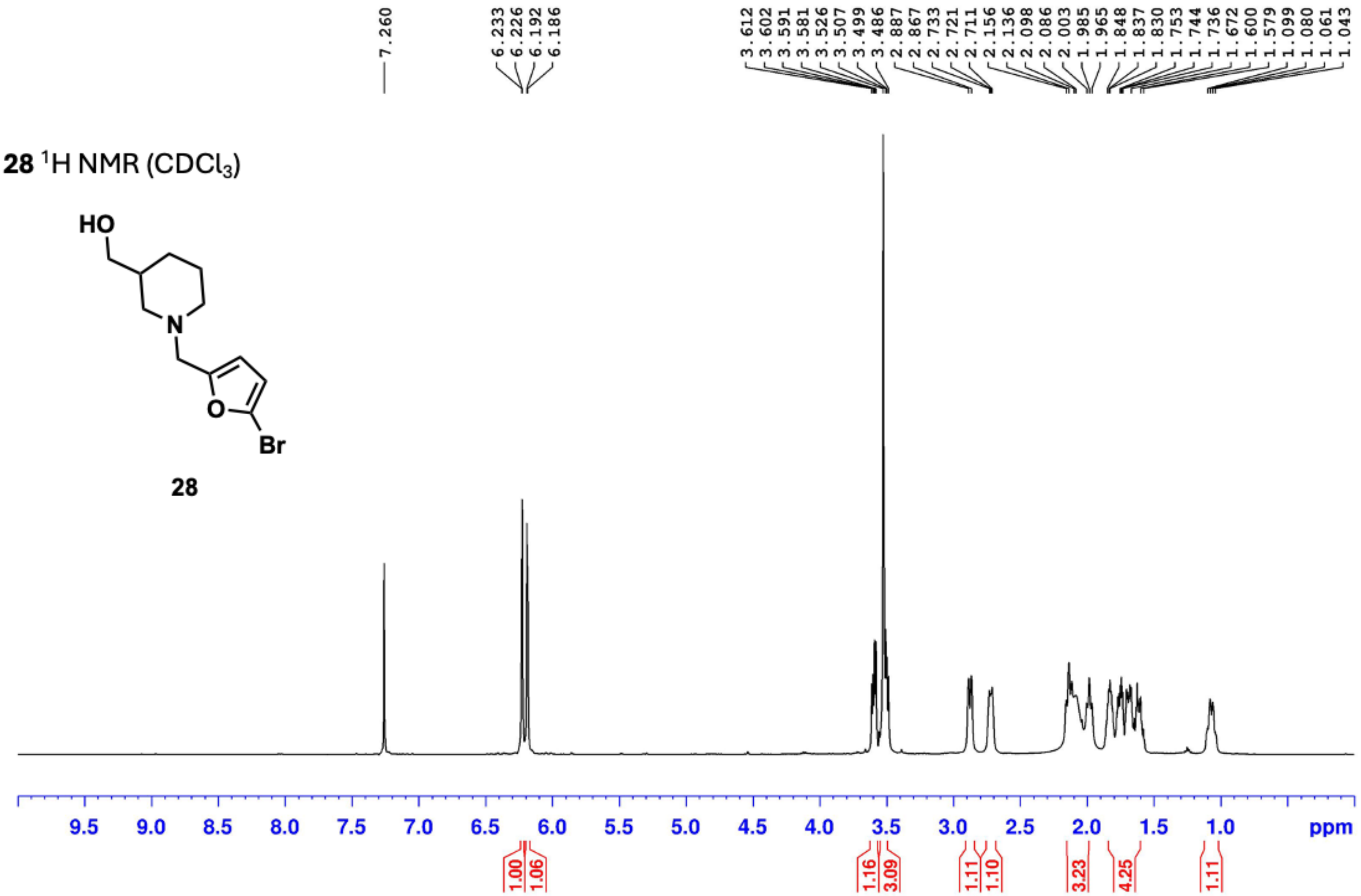

**7** $^1$H NMR ($CD_3OD$)

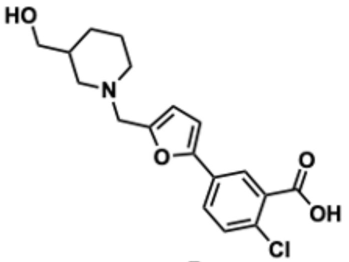


**7**

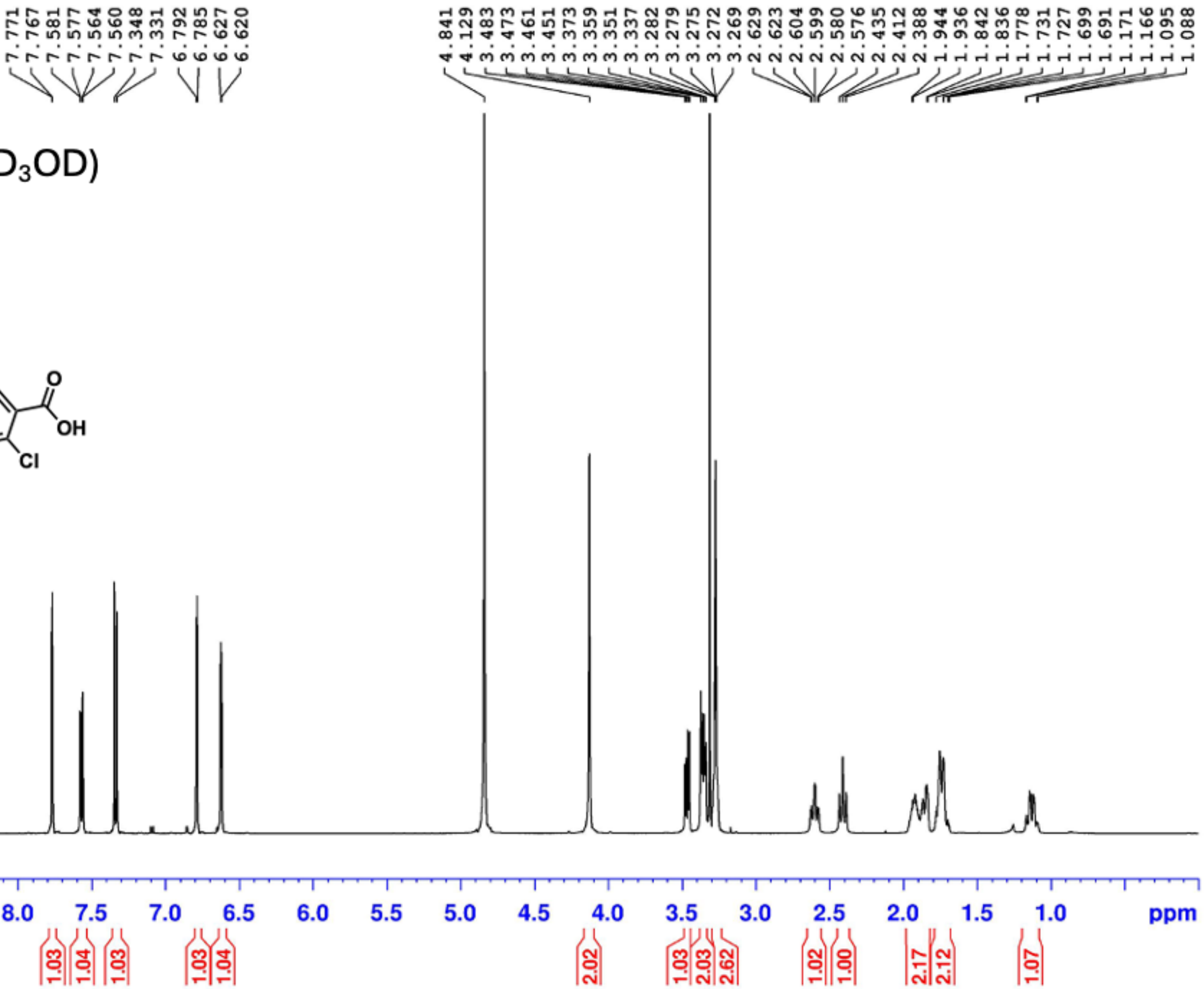

**7** $^{13}C$ NMR ($CD_3OD$)

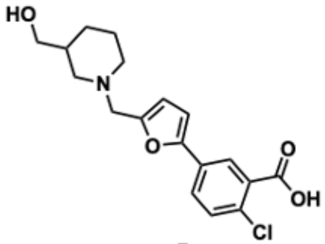


**7**

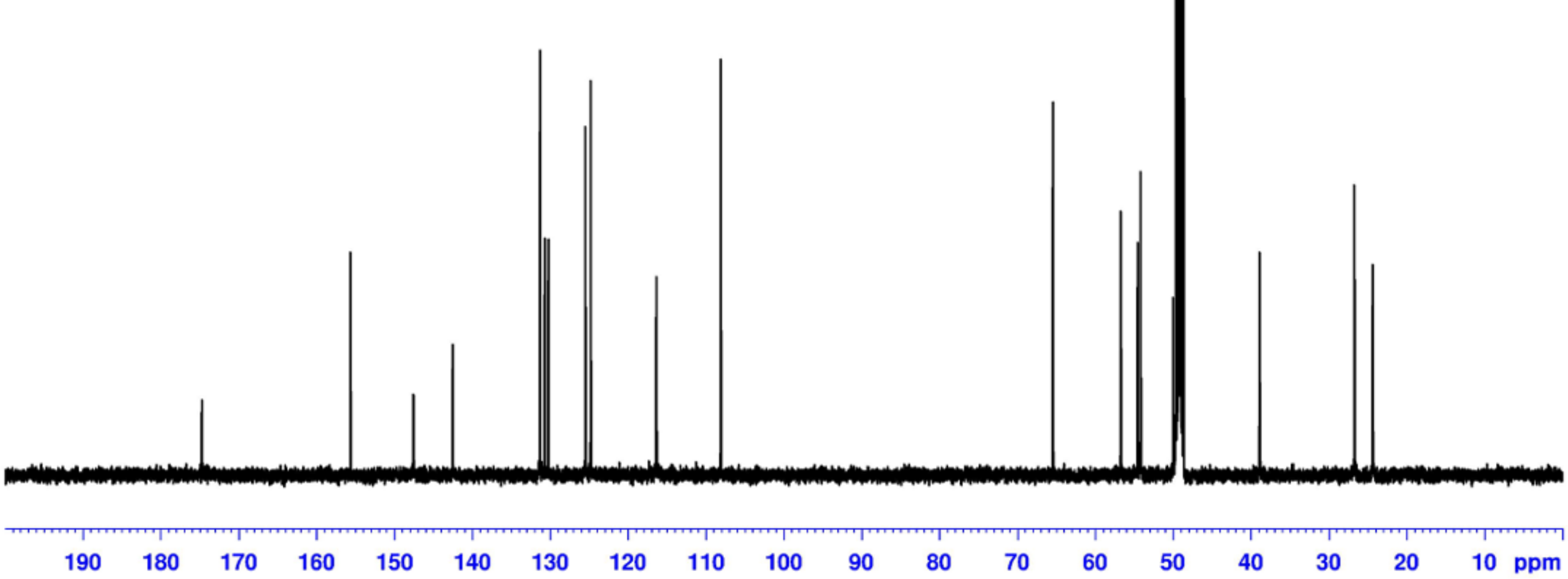

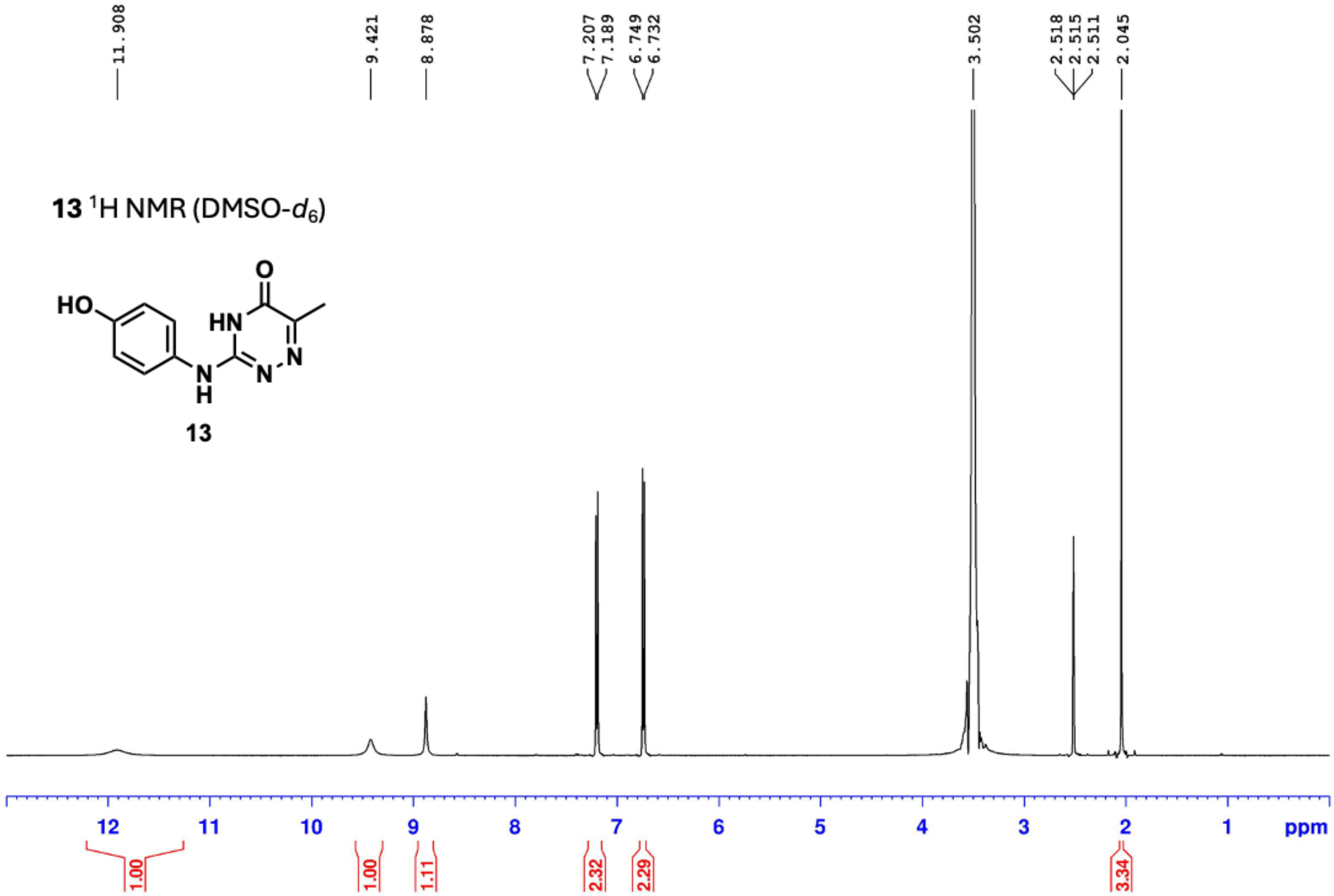
**13** $^{1}$H NMR (DMSO-$d_6$)
HO
HN
O
N
N
N
H
13
11.908
9.421
8.878
7.207
7.189
6.749
6.732
3.502
2.518
2.515
2.511
2.045
12
11
10
9
8
7
6
5
4
3
2
1
ppm
1.00
1.00
1.11
2.32
2.29
3.34

**13** $^{13}C$ NMR (DMSO-$d_6$)

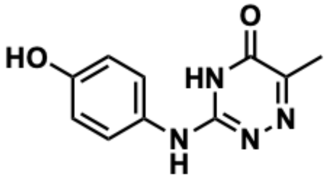


**13**

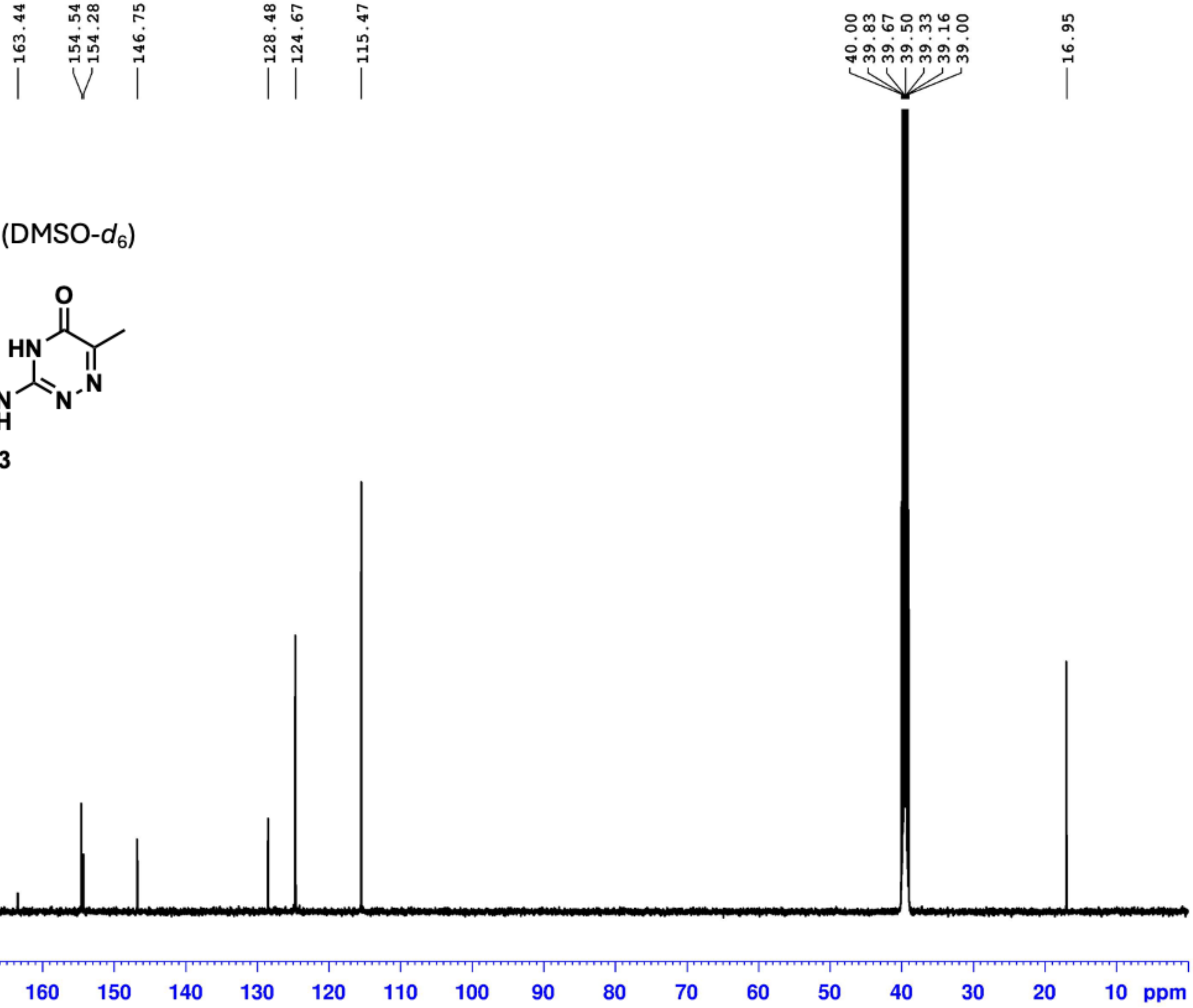

**34** $^{1}$H NMR (DMSO-$d_6$)

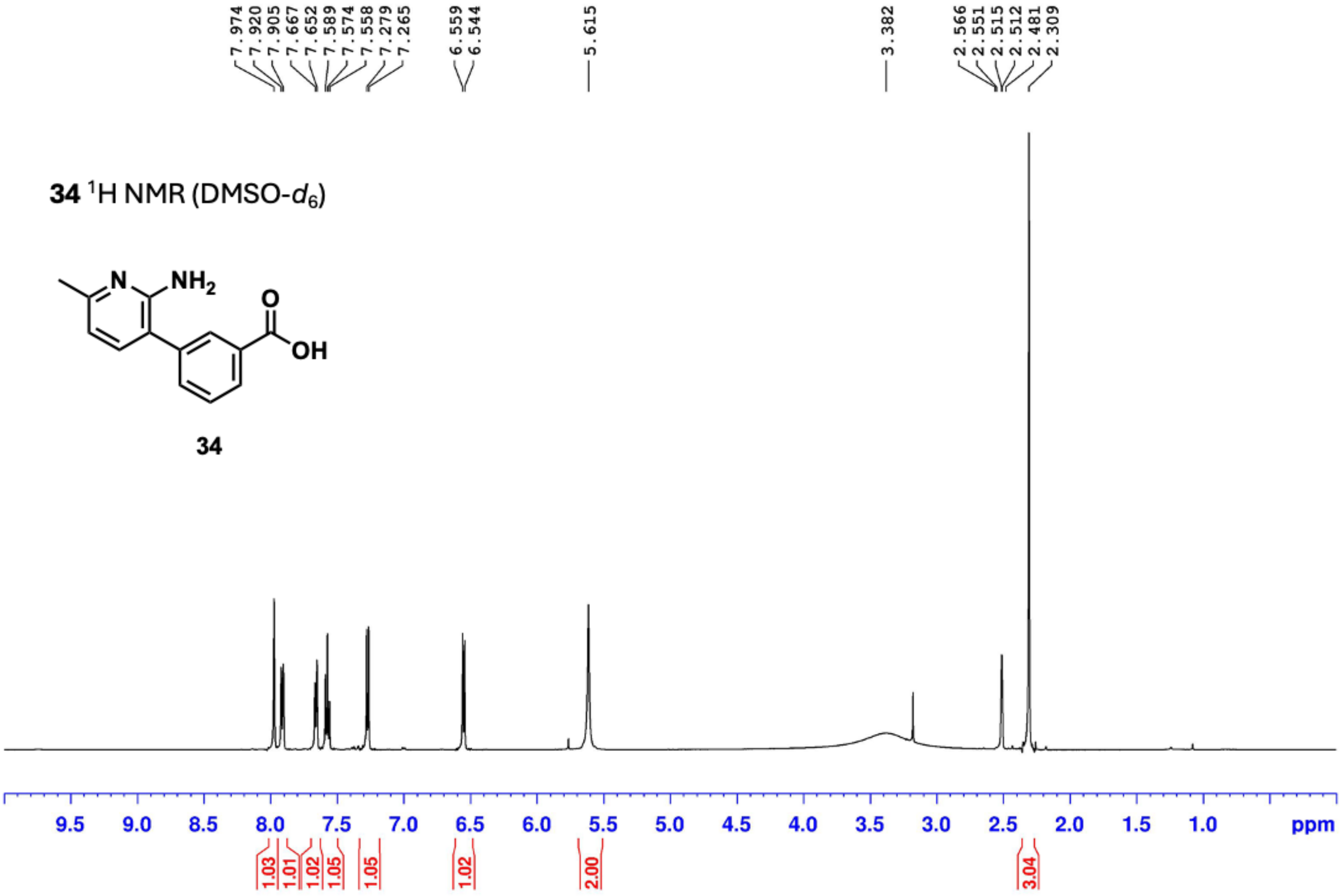


**34**

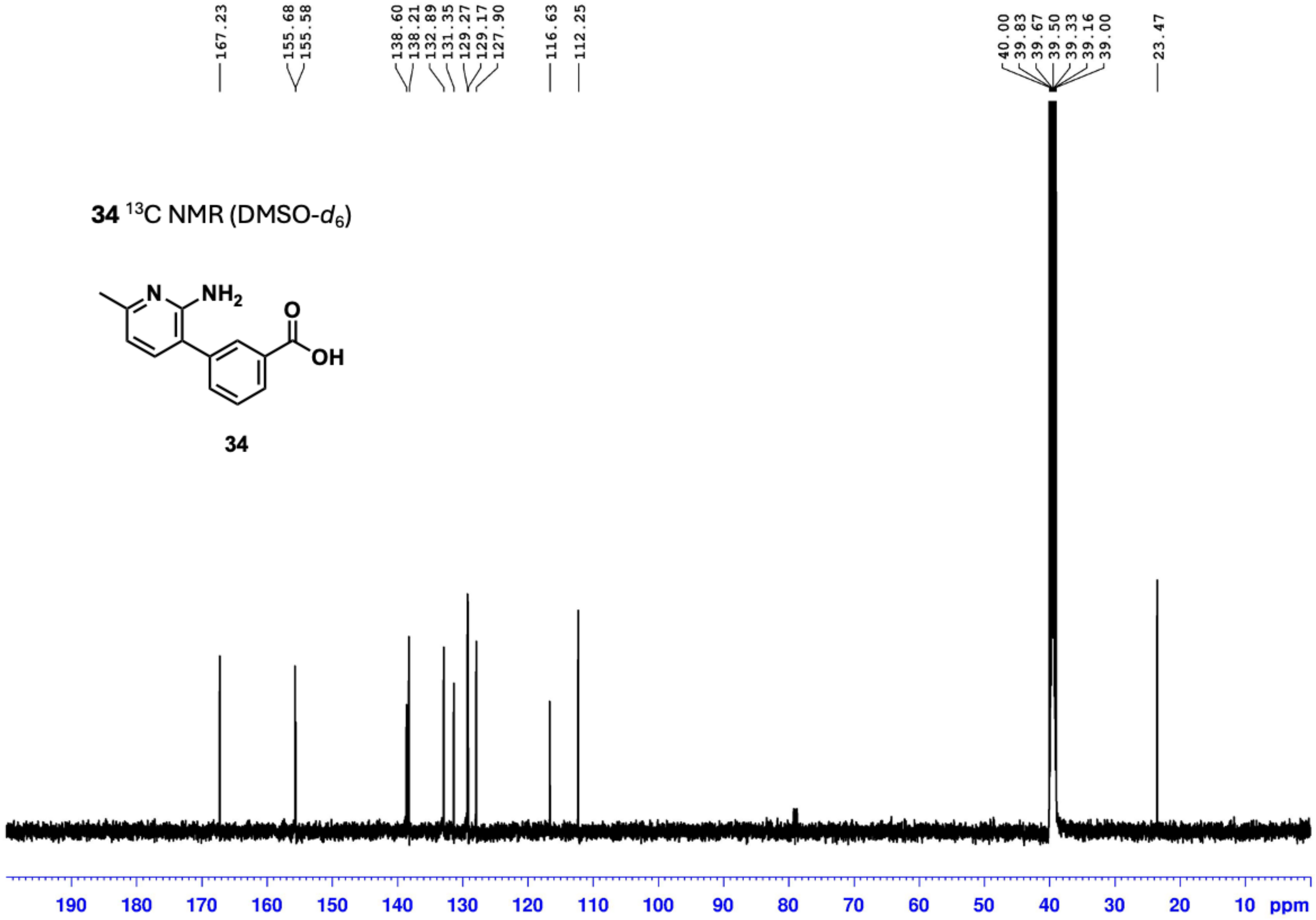

34 13C NMR (DMSO-d6)
NH2
N
O
OH
34
167.23
155.68
155.58
138.60
138.21
132.89
131.35
129.27
129.17
127.90
116.63
112.25
40.00
39.83
39.67
39.50
39.33
39.16
39.00
23.47
190 180 170 160 150 140 130 120 110 100 90 80 70 60 50 40 30 20 10 ppm

**16** $^1$H NMR ($CD_3OD$)

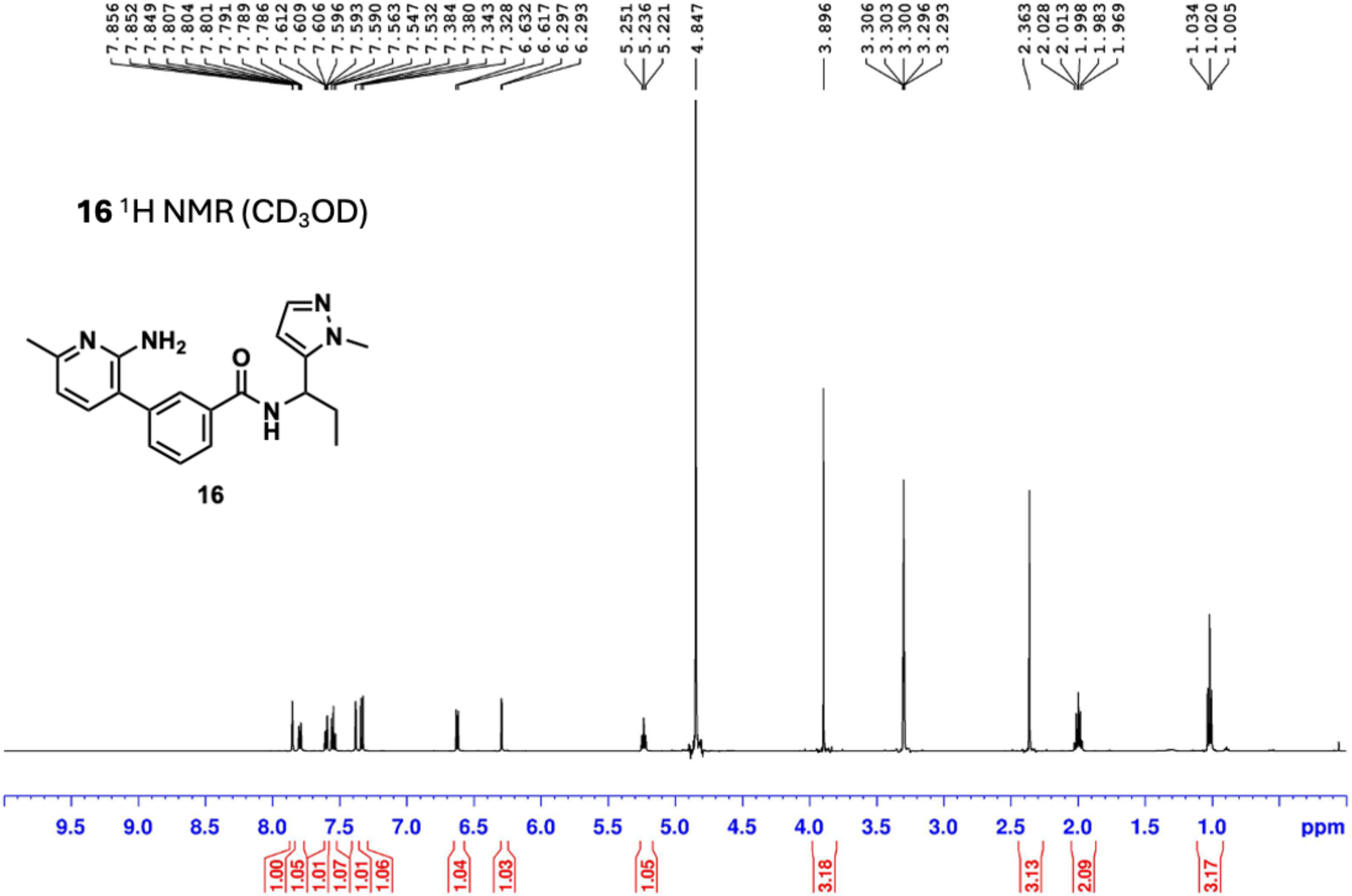


**16**

**16** $^{13}$C NMR ($CD_3OD$)

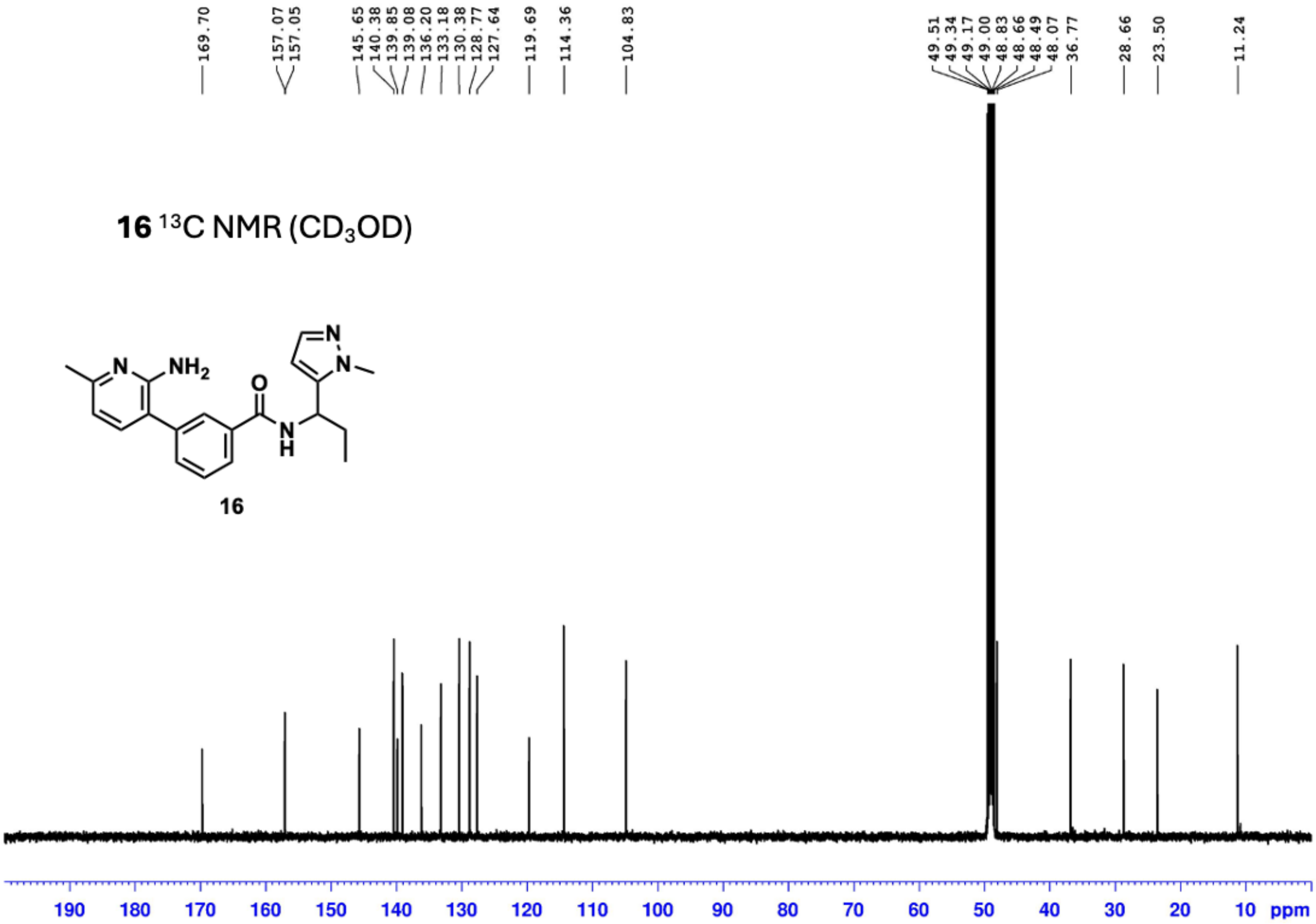

**16**
